# Genetic interaction between *Adgrg6* and *Sox9* reveals a feedforward mechanism for postnatal spinal stability

**DOI:** 10.64898/2026.04.06.716681

**Authors:** Valeria Aceves, Ziyi Xu, Cheng-Hai Zhang, Jongkil Kim, Joshua Ito, Kylie Mathers, Andrew B. Lassar, Zhaoyang Liu, Ryan S. Gray

## Abstract

Adolescent idiopathic scoliosis (AIS) is the most common spinal disorder in children and is best understood as a complex polygenic condition. Genome-wide association studies (GWAS) have identified several AIS risk loci, including regions near *ADGRG6* and *SOX9*, yet the functional mechanisms underlying AIS heritability remain poorly defined. Here, we use spatial transcriptomics to characterize altered gene expression in the spine of a conditional *Adgrg6* mutant mouse model of AIS (*Adgrg6-cKO*), revealing reduced expression of *Sox9* and several components of extracellular matrix organization in the intervertebral disc. We further show that SOX9 occupies regions of open chromatin within the *Adgrg6* locus in cells isolated from the mouse intervertebral disc. Finally, we demonstrate a strong genetic interaction between *Adgrg6-cKO* and a hypomorphic *Sox9* allele that increases both the penetrance and severity of AIS-like pathology in mice. Collectively, these findings support a self-reinforcing feedforward regulatory circuit, where *Adgrg6* and *Sox9* are co-regulated to maintain extracellular matrix gene expression in the annulus fibrosus and paraspinal tissues. These findings provide mechanistic insight into the functional significance of AIS-associated GWAS loci near *ADGRG6* and *SOX9* and establish combined *Adgrg6–Sox9* insufficiency as a tractable model of polygenic scoliosis susceptibility.

- *Adgrg6* maintains *Sox9* and extracellular matrix gene expression in the intervertebral disc
- SOX9 directly occupies open chromatin at the *Adgrg6* locus in SOX9⁺ skeletal cells
- *Adgrg6* and *Sox9* genetically interact to increase scoliosis incidence and severity
- Combined loss of *Adgrg6* and *Sox9* drives ectopic cartilage formation and reduced vertebral bone mineral density in the thoracic spine

## INTRODUCTION

Adolescent idiopathic scoliosis (AIS) is the most common pediatric musculoskeletal disorder, affecting approximately 3% of school-age children worldwide ^1,2^. AIS typically presents as spine curvature around the onset of puberty, without obvious axial patterning or vertebral body fusion defects. Notably, the need for surgical correction or bracing is more frequent in females than in males. Population-based genome-wide association studies (GWAS) meta-analysis have identified several AIS susceptibility loci, predominantly within non-coding regions of the genome, supporting a complex polygenic etiology ^3–8^. In addition, a polygenic burden of rare variants across extracellular matrix genes likely contributes significantly to AIS risk ^4,9,10^. Taken together, an emerging model suggests that AIS results from the combined effects of complex polygenic variation, hormonal influences, and mechanical stresses encountered during rapid adolescent growth.

Maturation and homeostasis of a healthy, functional spine require the coordinated integration of multiple musculoskeletal tissues, including bone, cartilage, connective tissue, muscle, and the peripheral nervous system. Consistent with this, AIS-like spinal curvatures are observed in patients with connective tissue disorders featuring joint hypermobility, such as Marfan syndrome and subtypes of Ehlers-Danlos syndrome, as well as in neuromuscular conditions including muscular dystrophy, Rett syndrome and pediatric spinal cord injury. This suggests that subclinical defects in connective tissue integrity or neuromuscular regulation may underlie susceptibility to AIS. In support of this, animal models with targeted disruption of neural circuits involved in proprioception ^11^ or central pattern generation ^12,13^, spinal cord central canal physiology ^14–16^, muscle physiology ^17–20^ or connective tissue gene expression and mechanics ^21^ all recapitulate AIS-like phenotypes.

Generalized joint hypermobility is a recognized risk factor for AIS ^22^, and postnatal dysregulation of connective tissue homeostasis has emerged as a recurring theme in human genetic studies of scoliosis ^23^. Multiethnic GWAS have identified intronic variants in *GPR126/ADGRG6* as strongly associated with human AIS ^7,24^. ADGRG6 belongs to a large class of adhesion G-protein coupled receptors (aGPCRs) that act as mechanosensors through force-based mechanisms, whereby dissociation of the N-terminal fragment from the seven-transmembrane domain elicits downstream signal transduction ^25–27^. Our prior studies demonstrated that conditional loss of *Adgrg6* in mice recapitulates AIS, alters the biomechanical properties of tendons, and reduces SOX9 protein expression in the intervertebral disc (IVD) through disruption of CREB signaling ^21,28^.

*Sox9* is a key transcription factor that controls skeletal development and postnatal homeostasis of the spine ^29,30^. Notably, the postnatal loss of *Sox9* causes reduced skeletal growth, scoliosis, and leads to altered IVD gene expression, including a significant reduction in *Adgrg6/Gpr126* expression ^30^. GWAS have identified variants in the *SOX9* locus associated with human AIS ^8,31^, and coding variants in the transactivation middle (TAM) domain have been independently implicated in human scoliosis ^32,33^. Consistent with this, our prior studies showed that a microdeletion in the *Sox9* TAM domain causes late-onset scoliosis in mice, accompanied by reduced expression of SOX9, ADGRG6, and several ECM proteins in the annulus fibrosus of the IVD ^33^.

The complex polygenic architecture underlying AIS susceptibility and severity remains poorly understood. To address this, we performed unbiased spatial transcriptomic profiling and RNA *in situ* hybridization in the conditional *Adgrg6* mouse model of AIS ^21,28^, documenting reduced expression of *Sox9* and ECM genes in the IVD. We further analyzed chromatin accessibility and SOX9 occupancy in *Sox9*-expressing skeletal cells, revealing that SOX9 binds putative regulatory regions within *Adgrg6*. Finally, we show that genetic interactions between conditional *Adgrg6* and *Sox9* TAM domain mutations increase the penetrance and severity of scoliosis in mice. Taken together with our prior work ^21^, these findings indicate that *Adgrg6* and *Sox9* operate in a feedforward regulatory loop to maintain gene programs essential for postnatal spinal stability.

## RESULTS

### Spatial Transcriptomics Reveals Reduced Expression of Genes Related to Cartilage Development and Extracellular Matrix Organization in the Intervertebral Discs of the *Adgrg6-cKO* Mutant Mouse Model of AIS

We previously showed that conditional loss of *Adgrg6* in osteochondral progenitors (Col2a1-Cre; *Adgrg6^f/f^*; hereafter *Adgrg6-cKO*) or in connective tissue (ScxCre; *Adgrg6^f/f^*) models adolescent idiopathic scoliosis (AIS) in mice ^21^. To generate a comprehensive, unbiased map of how *Adgrg6* contributes to the molecular regulation of the spine, we performed spatial transcriptomics on P20 spine sections from wild-type and *Adgrg6-cKO* mice. Unsupervised clustering of the entire spine section identified nine transcriptionally distinct populations, encompassing nucleus pulposus cells of the intervertebral disc (IVD), osteoblasts/osteoclasts, paraspinal muscle and adipose tissue, paraspinal connective tissue, and patches of dorsal root ganglion. Because osteoblast-lineage–driven CRE recombination of *Adgrg6* does not induce scoliosis in mice ^21^, all downstream differential expression analyses were restricted to capture spots overlaying IVD tissues (Fig. 1a–d).

**Figure 1.**
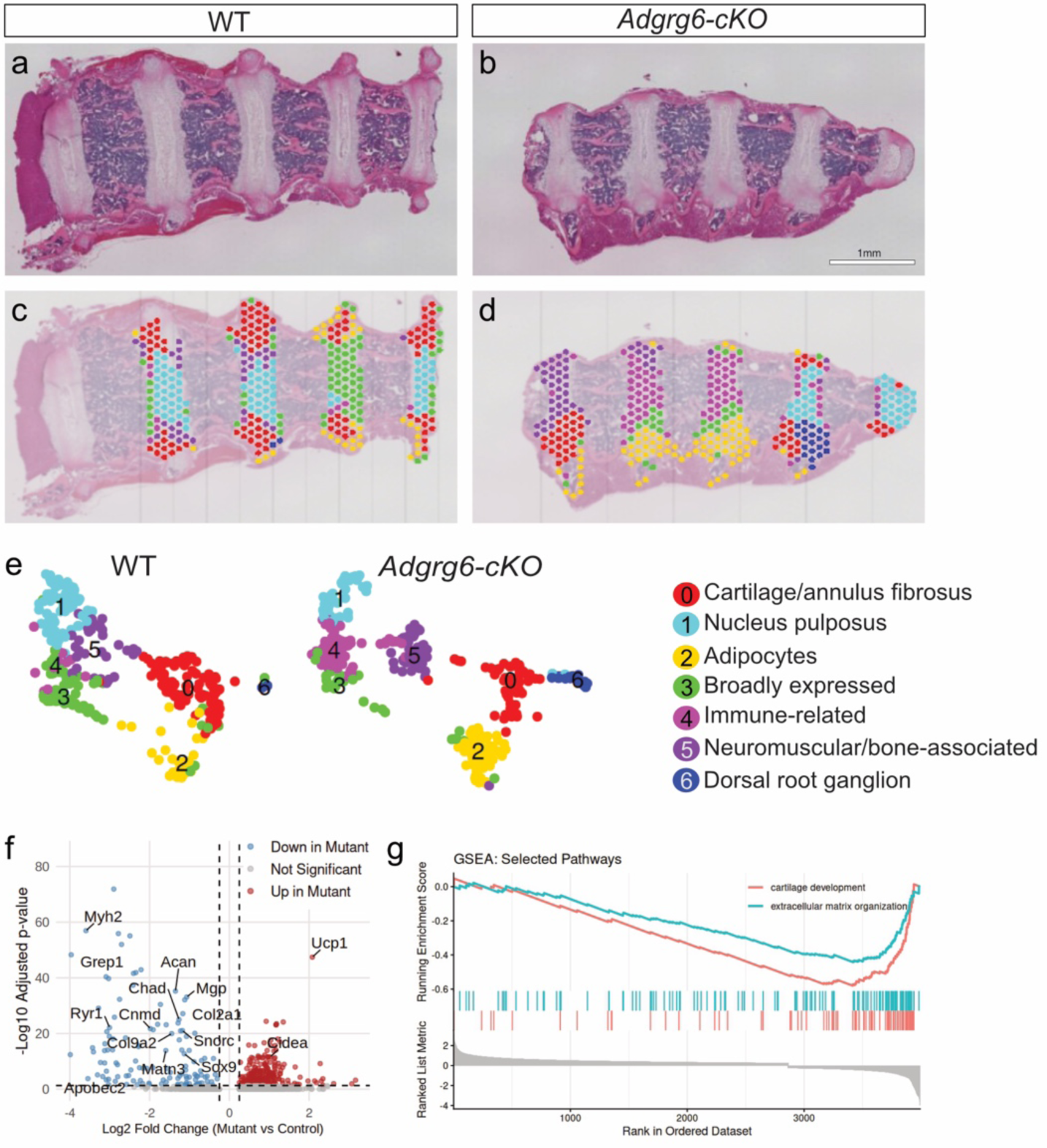
Spatial transcriptomics and differential expression of *Adgrg6-cKO* intervertebral disks (IVD). Hematoxylin and Eosin (H&E) staining of IVD sections from control and *Adgrg6-cKO* mice (a–b). Spatial DimPlots projecting transcriptomic clusters on control and *Adgrg6-cKO* samples (c–d). Uniform Manifold Approximation and Projection (UMAP) visualization of the first 9 principal components (e). Cells are colored by cluster identity and annotated by cell type (right). Volcano plot showing global differential expression (DE) between control and *Adgrg6-cKO* conditions (f). Significance was determined using a Wilcoxon Rank Sum test (adjusted p-value < 0.05, log2 fold change > 0.25). Gene Set Enrichment Analysis (GSEA) shows that cartilage development (red line, GO:0051216) and extracellular matrix organization (teal line, GO:0030198) are significantly downregulated in the *Adgrg6-cKO* compared to control (g). Scale bar = 1 mm.

The spine section is a structurally heterogeneous tissue in which metabolically active paraspinal muscle and adipose depots are in direct proximity to the fibrous connective tissues of the IVD. Because high-UMI transcripts released from these adjacent tissues can diffuse into neighboring capture spots, we implemented a per-spot muscle-score quality control filter as part of our standard preprocessing pipeline, excluding spots with greater than 1% muscle-transcript contribution prior to clustering and differential expression analysis. To validate that this filtering strategy was correctly identifying contamination rather than genuine IVD signal, we performed orthogonal RNA *in situ* hybridization for canonical muscle markers (*Myh2, Ryr1, Apobec2*) and immunofluorescence for UCP1 protein in *Adgrg6-cKO* and wild-type spine sections. Neither muscle transcripts nor UCP1 protein showed detectable expression within the IVD in either genotype (Supp. Fig. 1a–h), confirming that capture spots enriched for these signals reflected transcript diffusion from adjacent paraspinal tissues rather than genuine IVD expression differences. We note that *Ryr1* was of independent interest given recent reports linking *RYR1* mutations to proprioceptive defects and scoliosis in mice and humans ^10,20^; however its absence from IVD tissue by *in situ* analysis suggests that its differential expression is more related to technical challenges of using spatial transcriptomics in heterogenous tissue sections rather than any real interaction with *Adgrg6* signaling. Taken together, these findings underscore that paraspinal tissues exhibit greater RNA release and diffusion than the dense fibrous IVD, and that muscle-score filtering is a necessary and effective QC step for spatial transcriptomics in heterogeneous spinal tissue.

Following quality control at the spot level (feature counts, UMI counts, and mitochondrial content), the muscle-filtered dataset comprised 730 spots and 17,480 genes (Control: 371 spots; Mutant: 359 spots), with comparable QC metrics across conditions indicating high-quality IVD data suitable for robust clustering and differential expression analyses. Approximately 9 principal components captured the major variation in the dataset, yielding clear separation into seven biologically coherent clusters (Fig. 1e, Supp. Fig. 2, Table 1): cartilage/annulus fibrosus (AF) tissue (e.g., *Clec3a*, *Chad*, *Col2a1*, *Comp*; Supp. Fig. 2); nucleus pulposus (NP) tissue (e.g., *Krt19*, *T*, *Krt8*, *Arhgef16*) ^34,35^ (Supp. Fig. 4); adipocytes (*Lgals12*, *Ppara*, *Tmem79*, *Cidec*); broadly expressed genes (*Stfa3*, *Usp1*, *Bbip1*, *Cks2*); immune-related genes (*Fanca*, *Malt1*, *Lin28a*, *Tonsl*); neuromuscular/bone-associated genes (*Mobp*, *Prg2*, *Casq1*, *Agt*); and dorsal root ganglion markers (*Caly*, *Hand1*, *Epha10*, *Gprin1*). Differential expression analysis within IVD-restricted, muscle-filtered capture spots revealed significant downregulation of genes associated with cartilage development (GO:0051216) and extracellular matrix (ECM) organization (GO:0030198) in *Adgrg6-cKO* samples (Fig. 1f–g, Table 2). Collectively, these data indicate that *Adgrg6* modulates postnatal gene expression in the IVD, including maintenance of *Sox9* expression and regulation of ECM genes such as *Acan*, *Col9a1*, *Col2a1*, *Matn3*, *Col11a2*, *Col27a1*, and *Col15a1*.

**Table 1.**
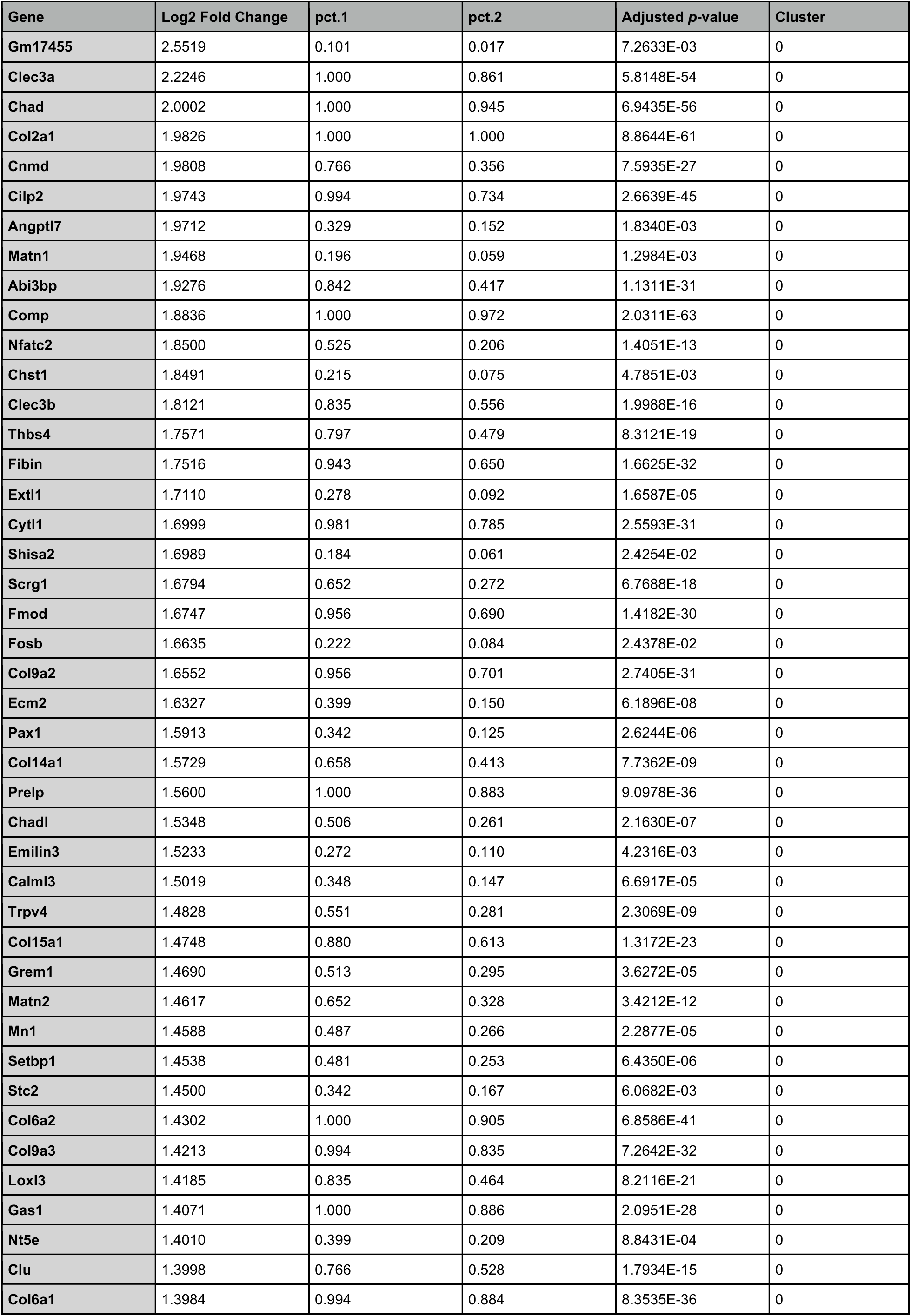
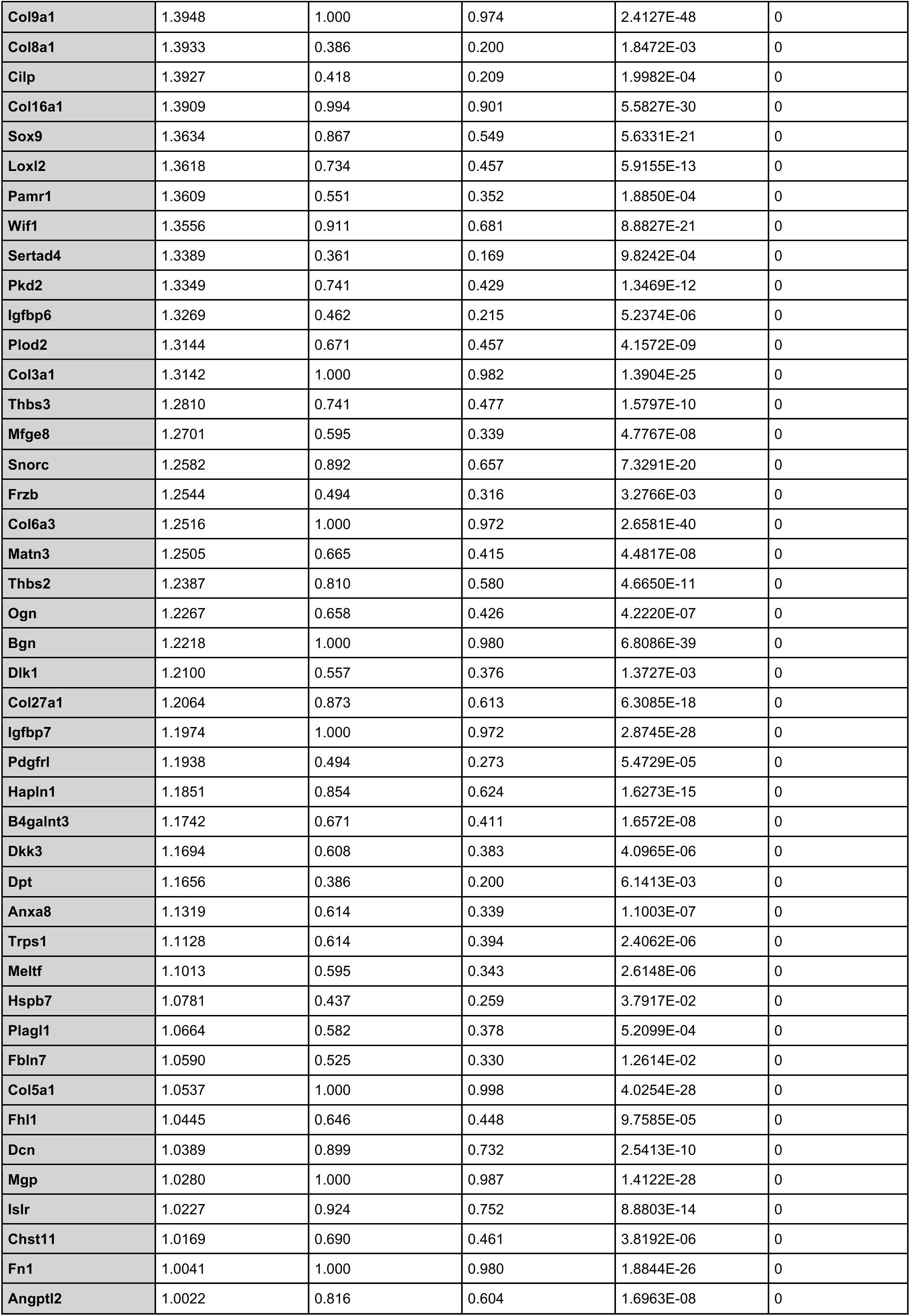
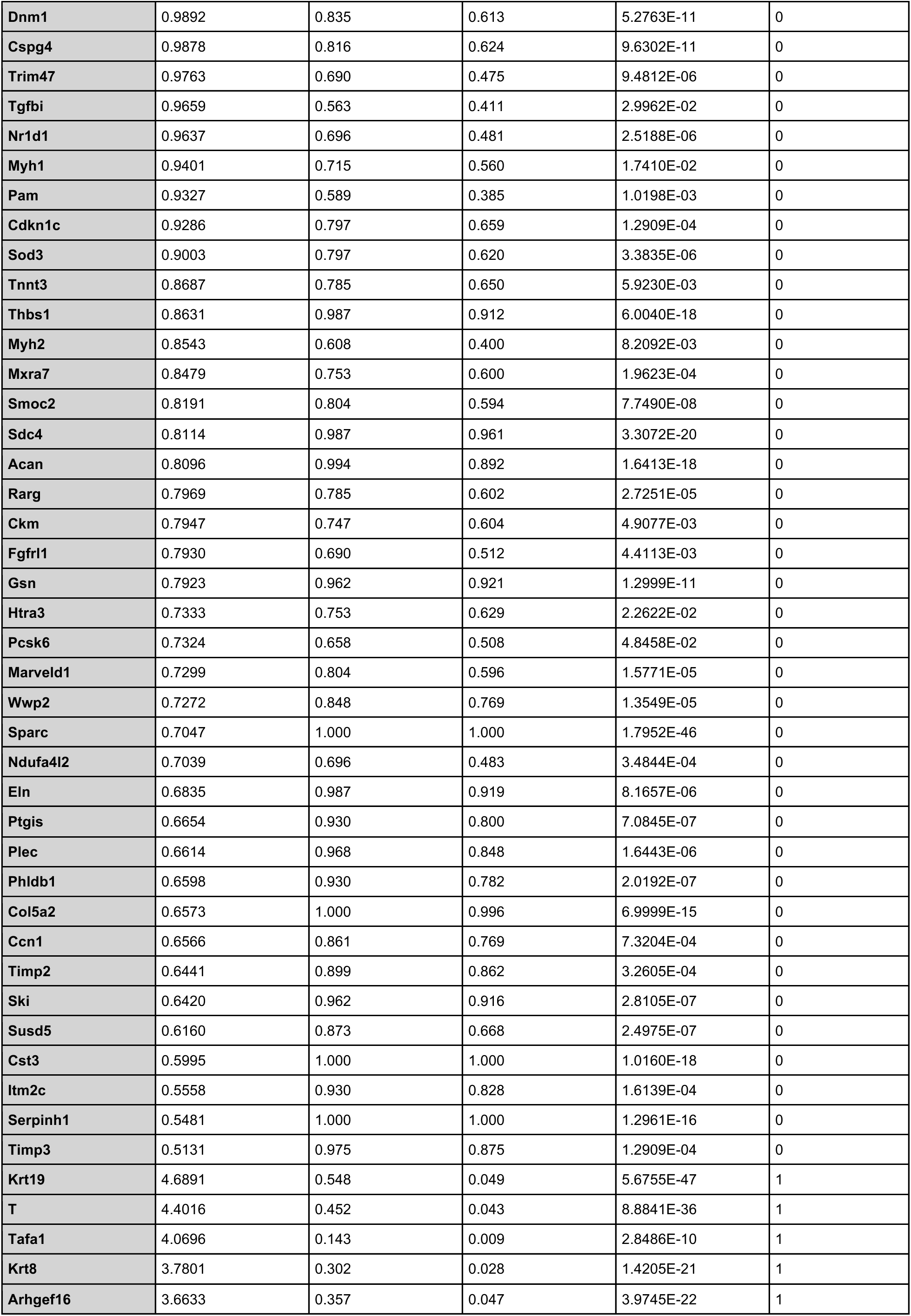
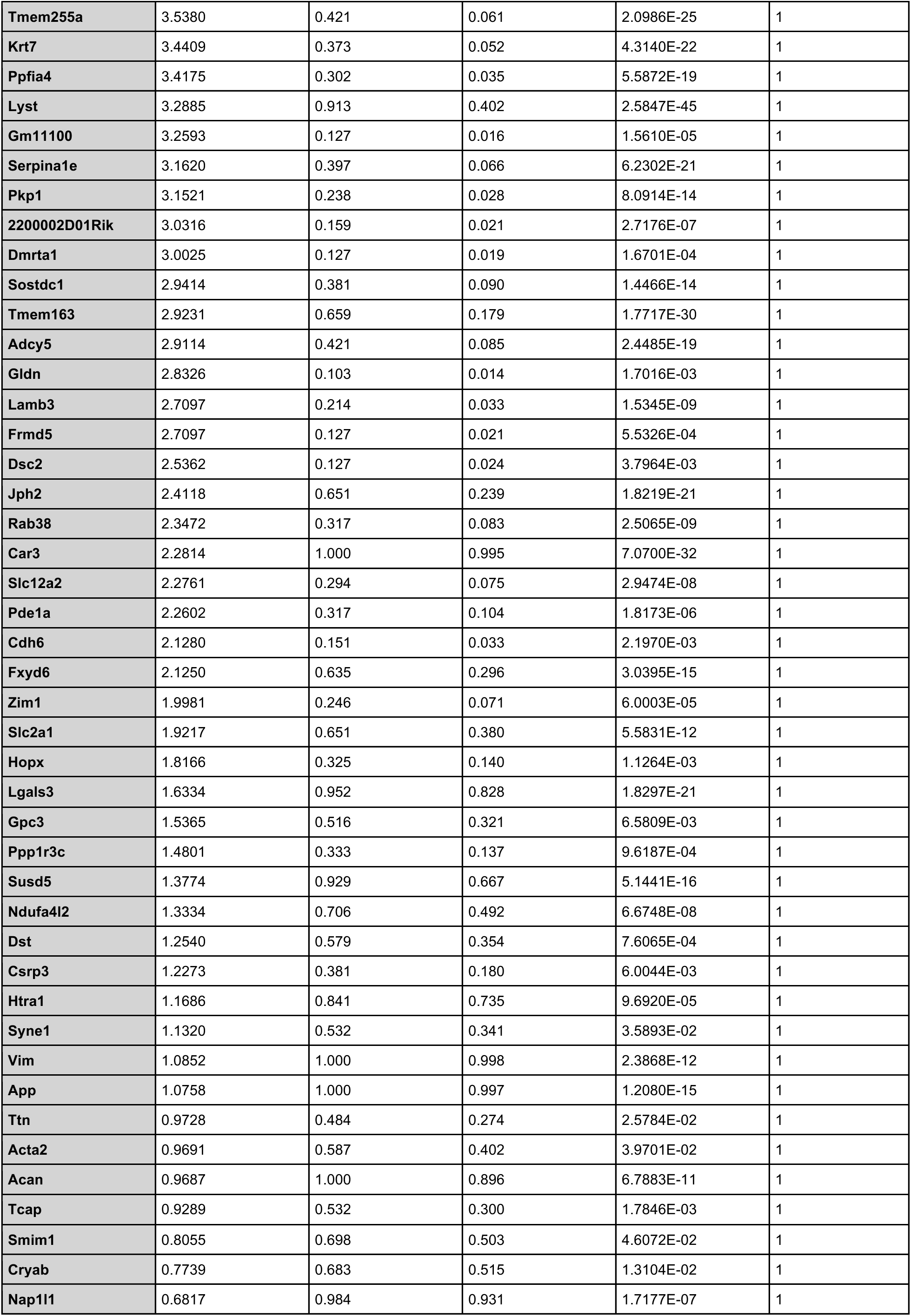
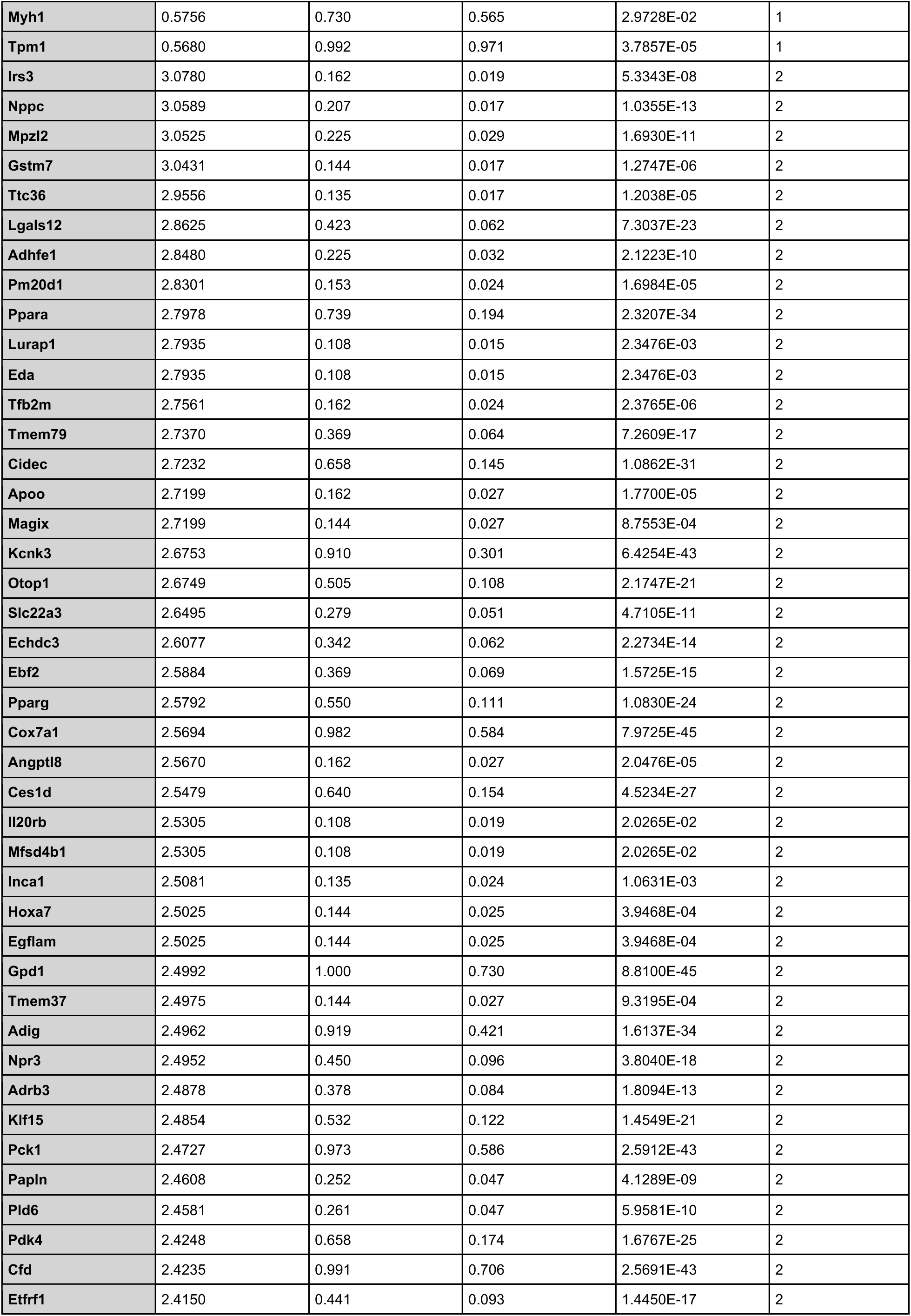
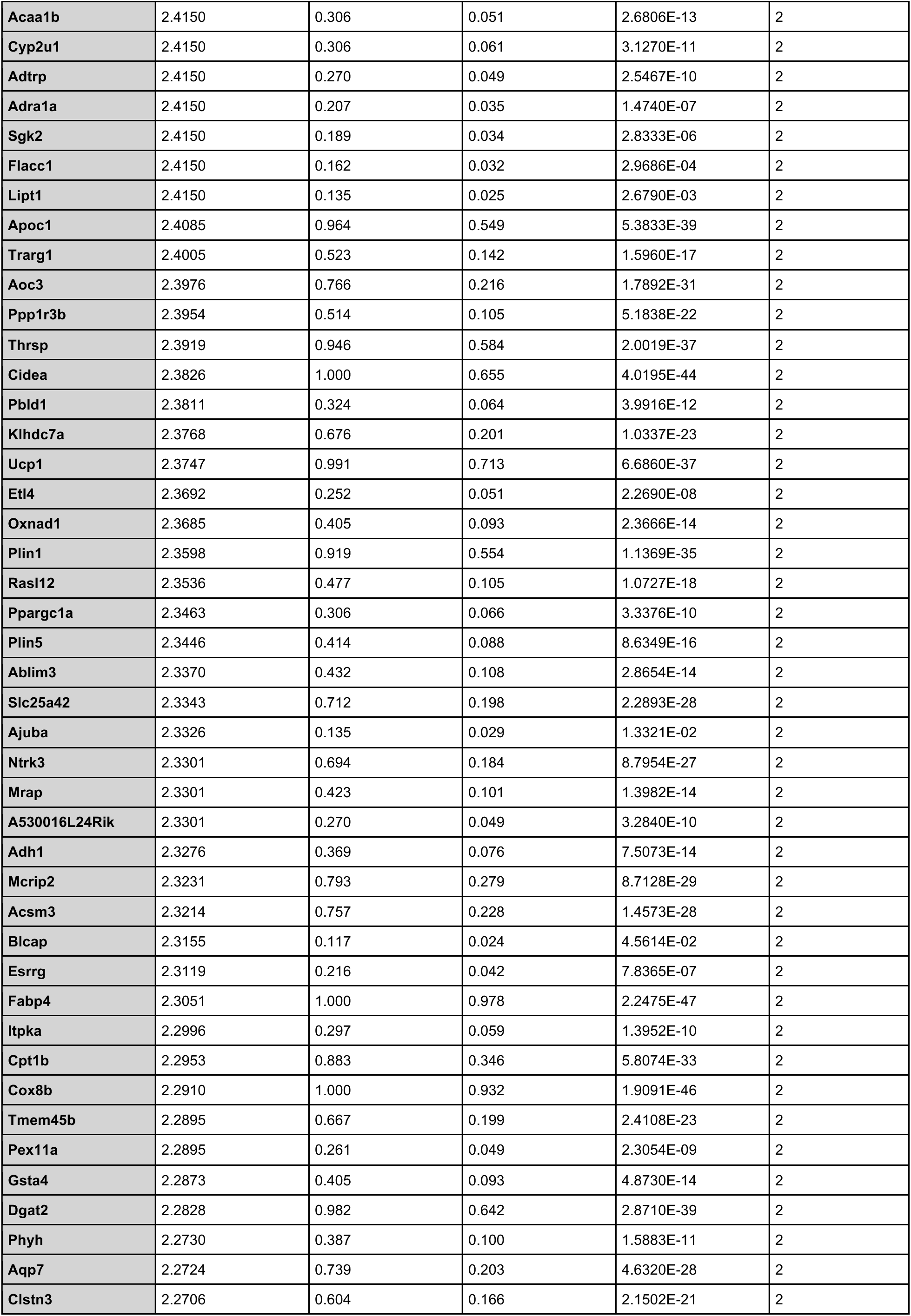
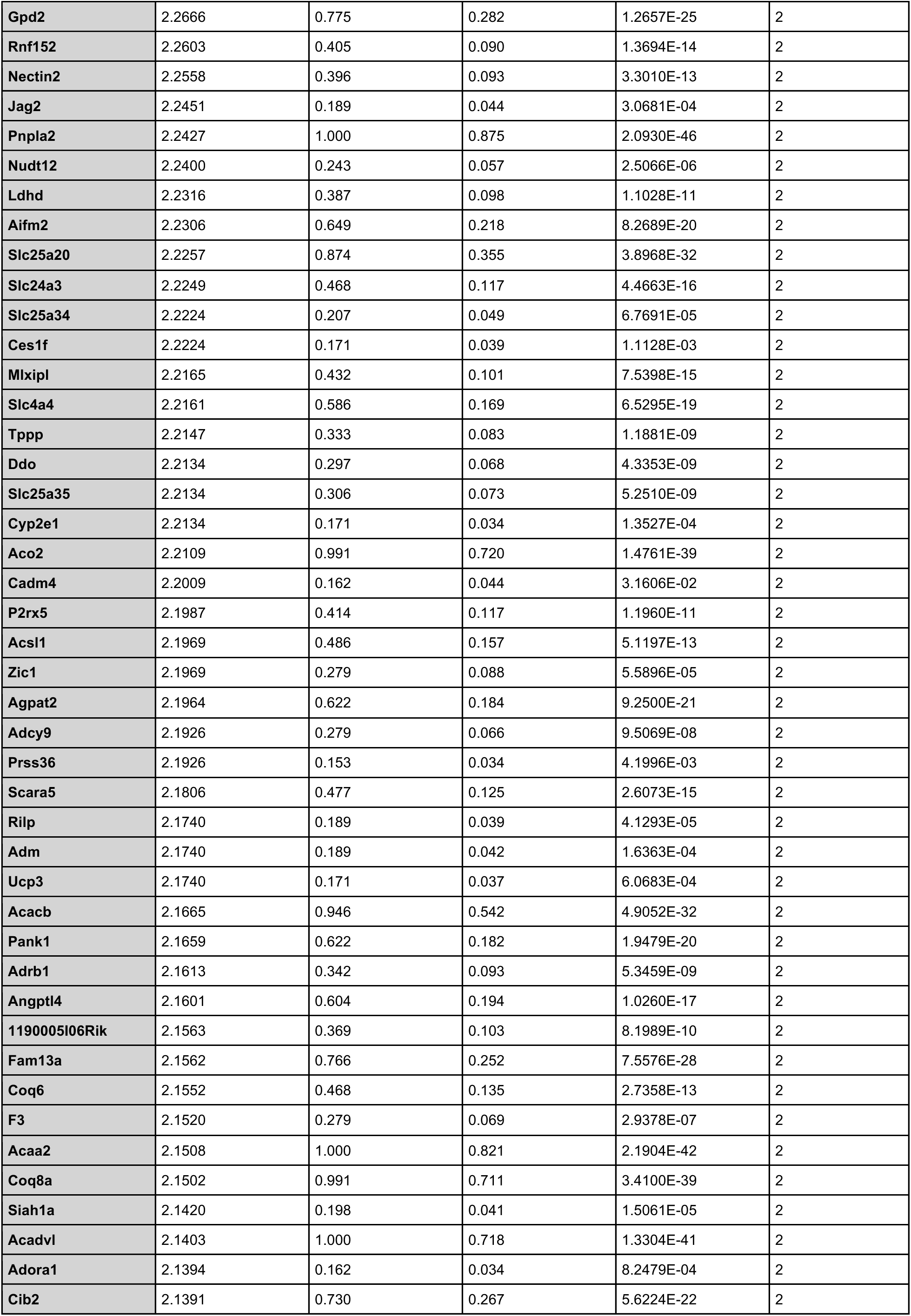
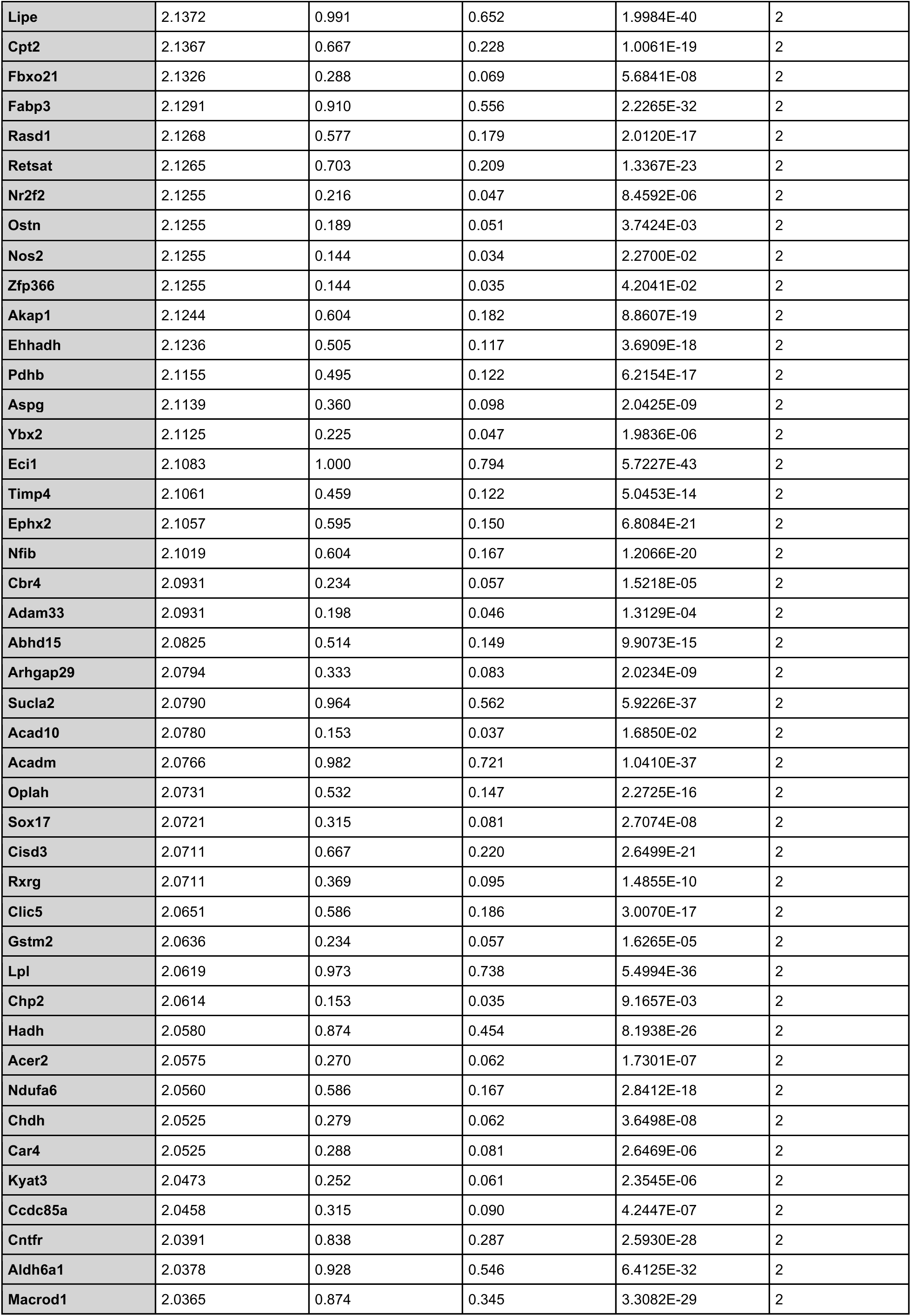
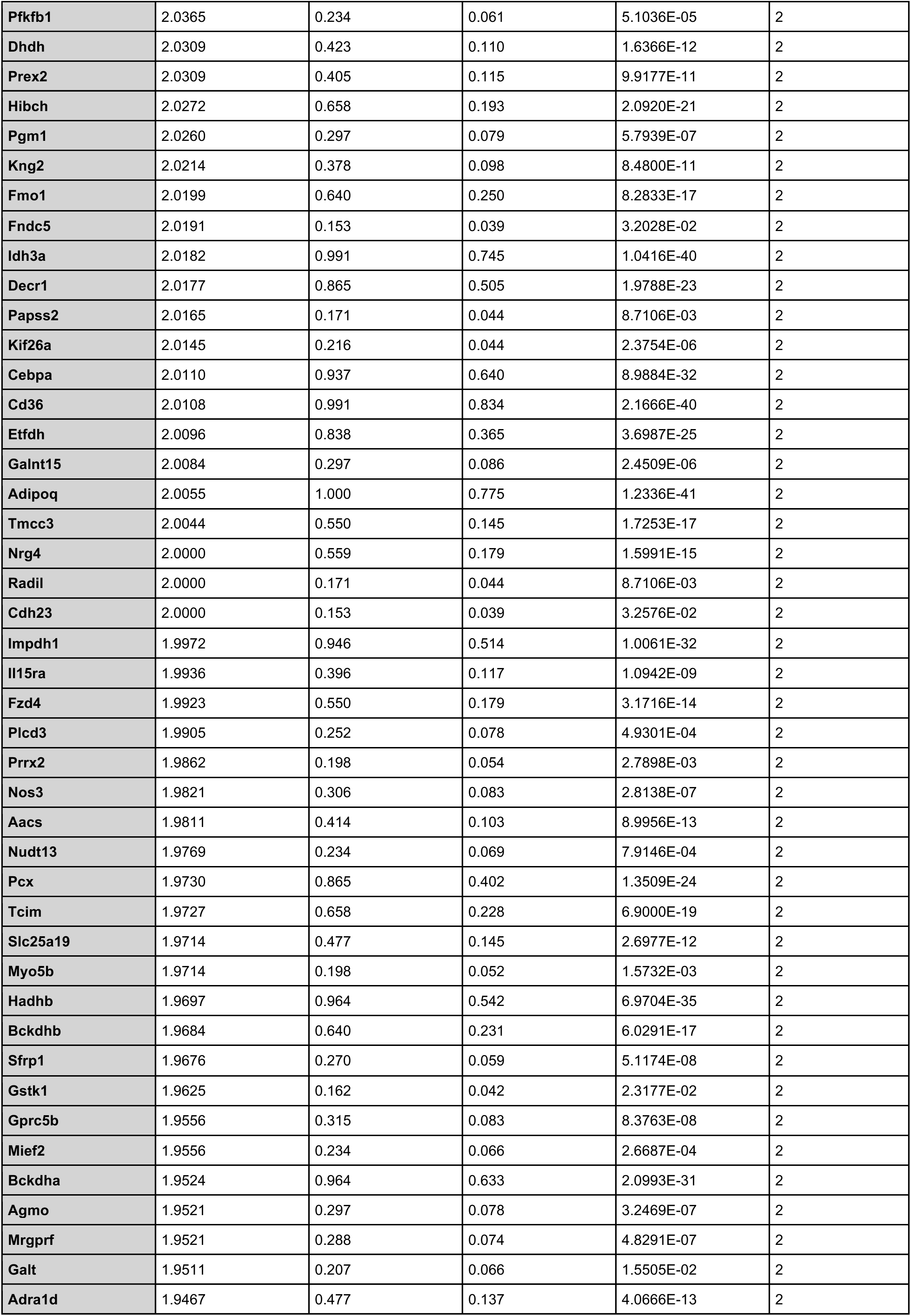
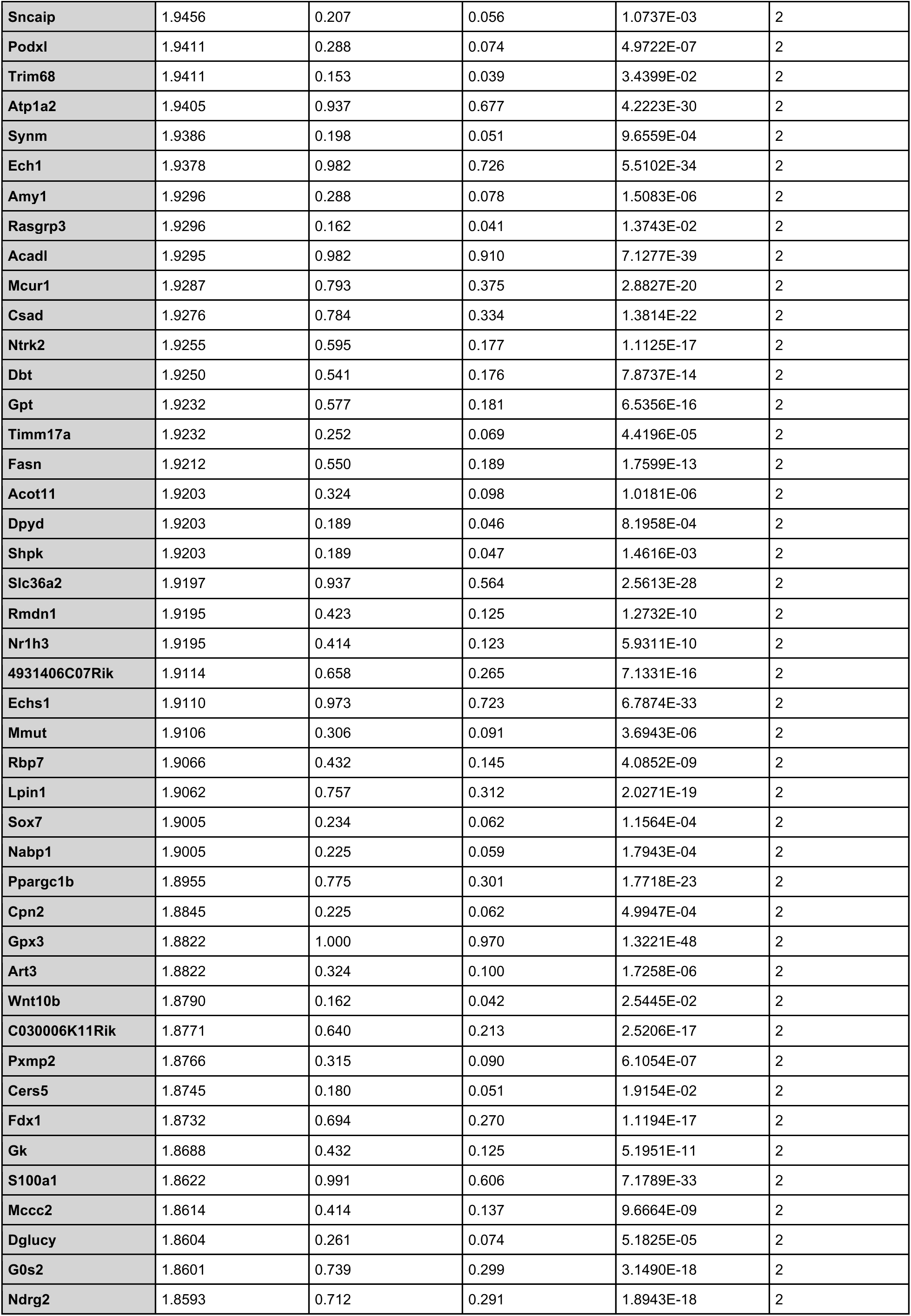
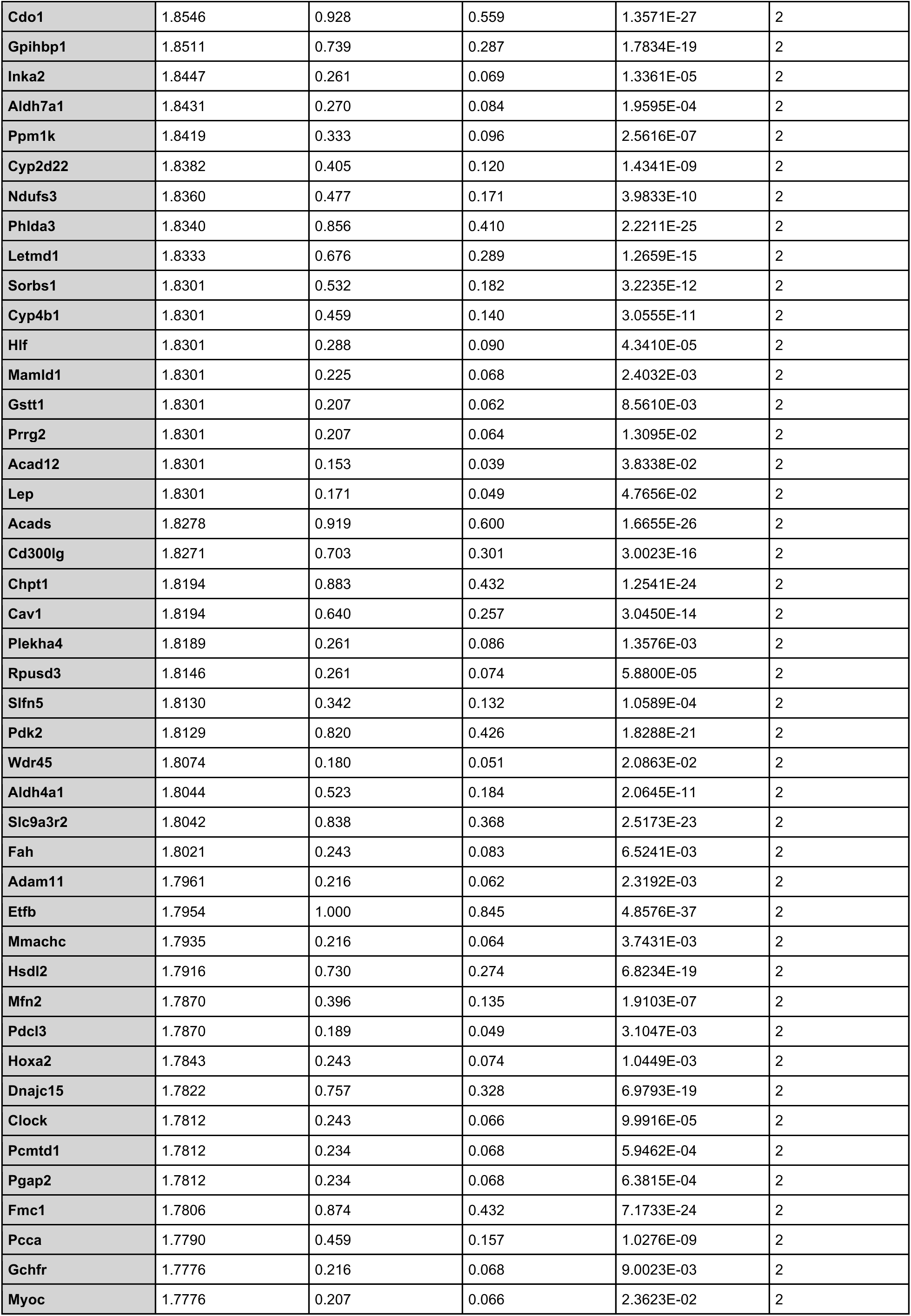
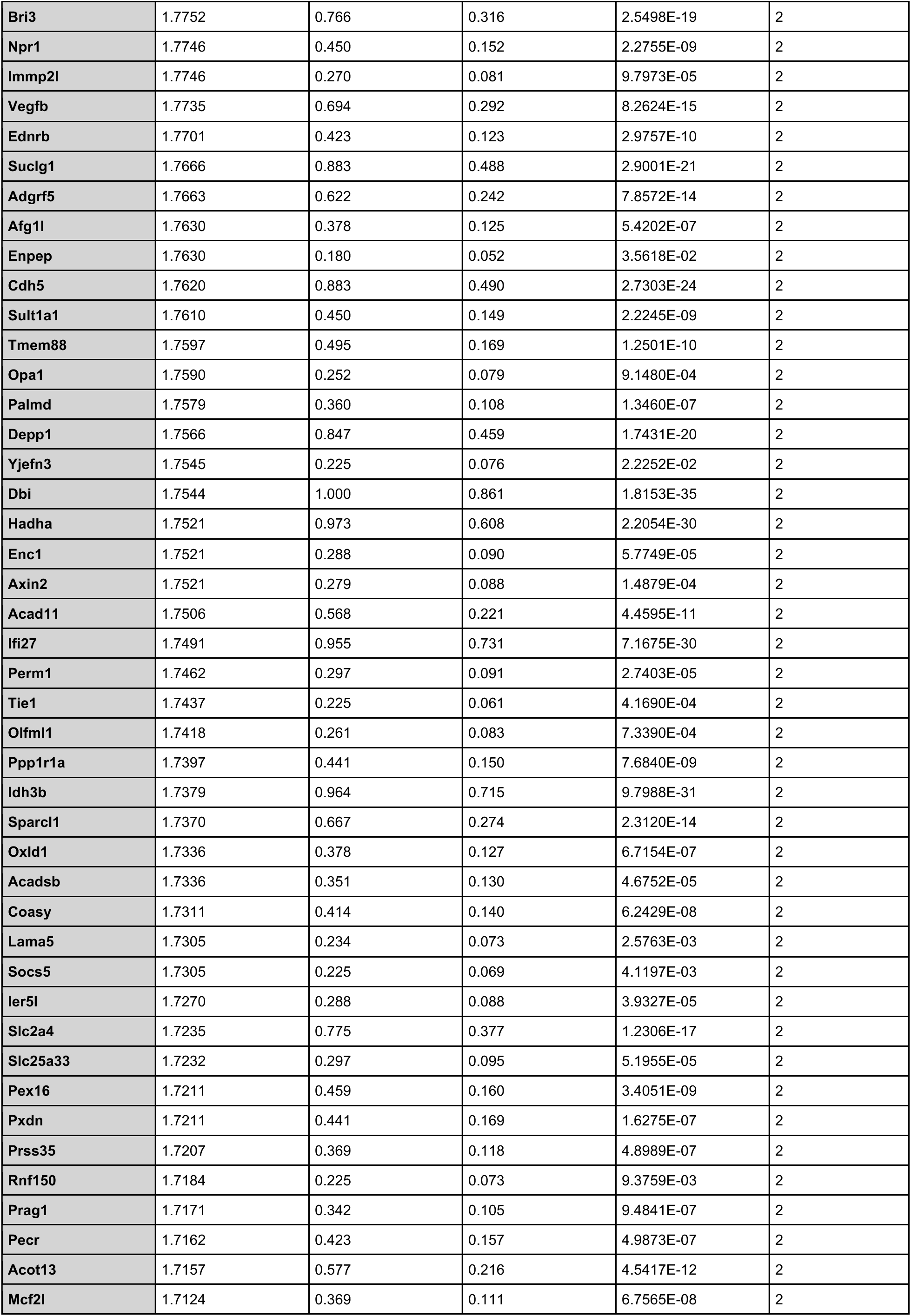
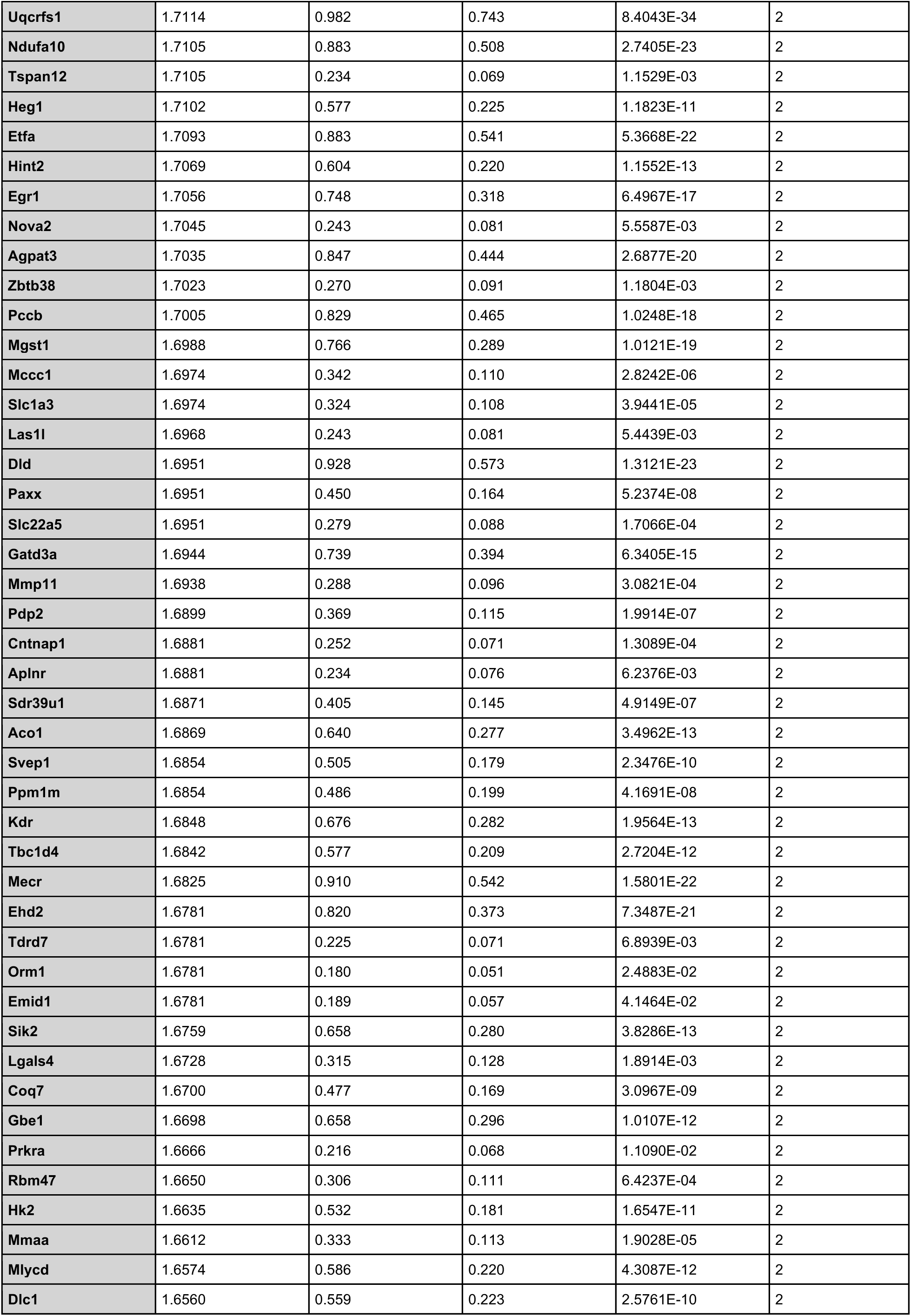
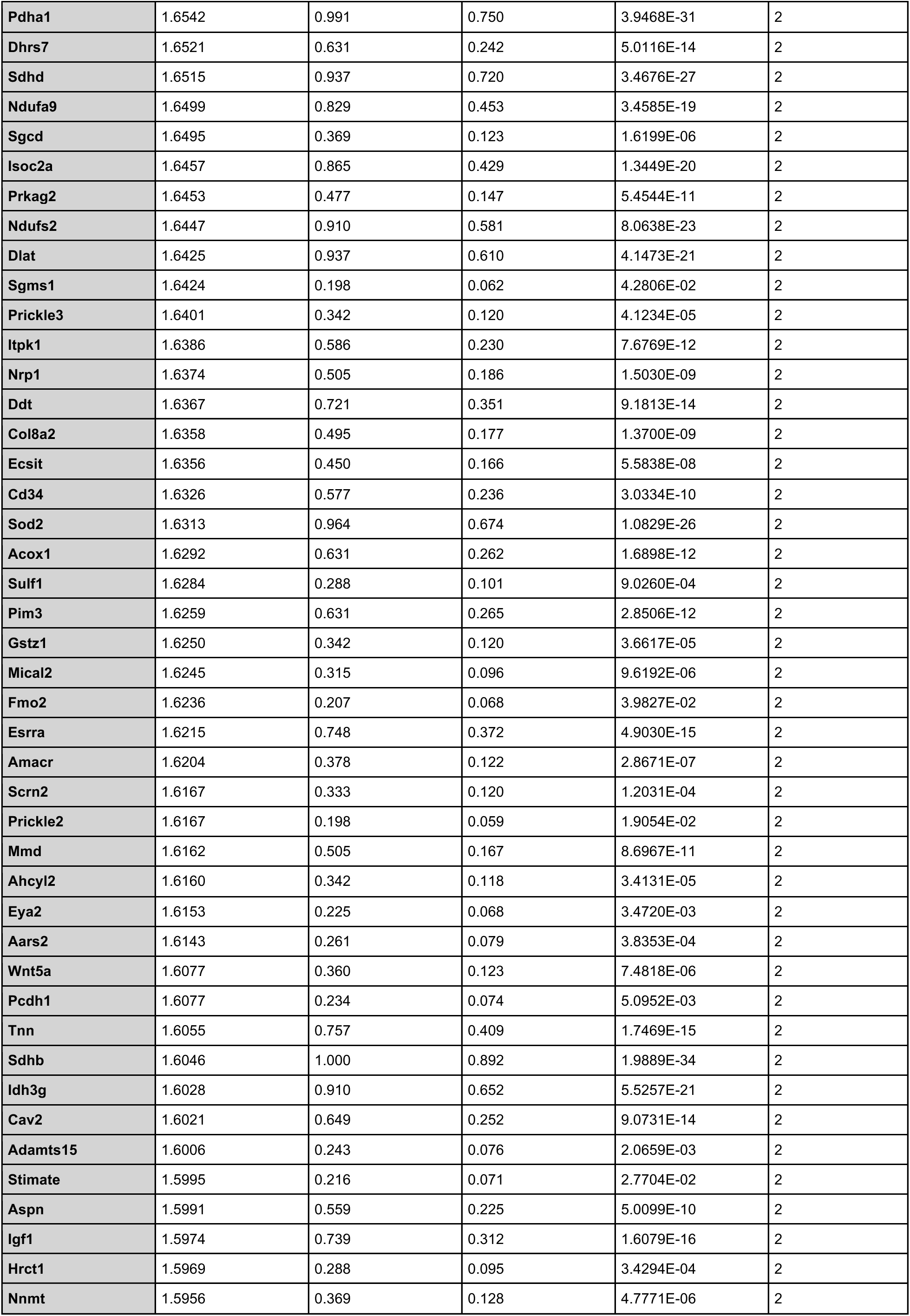
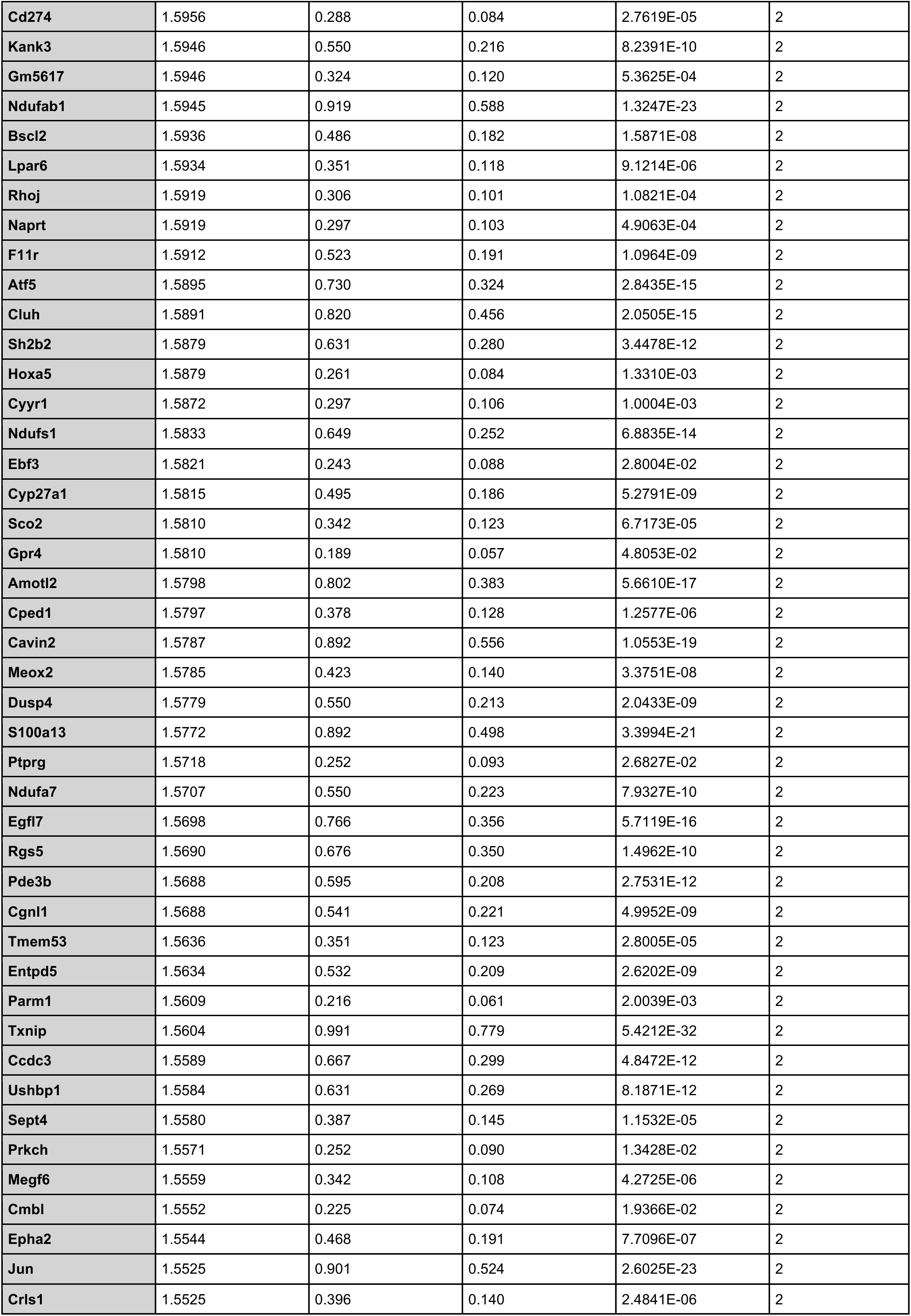
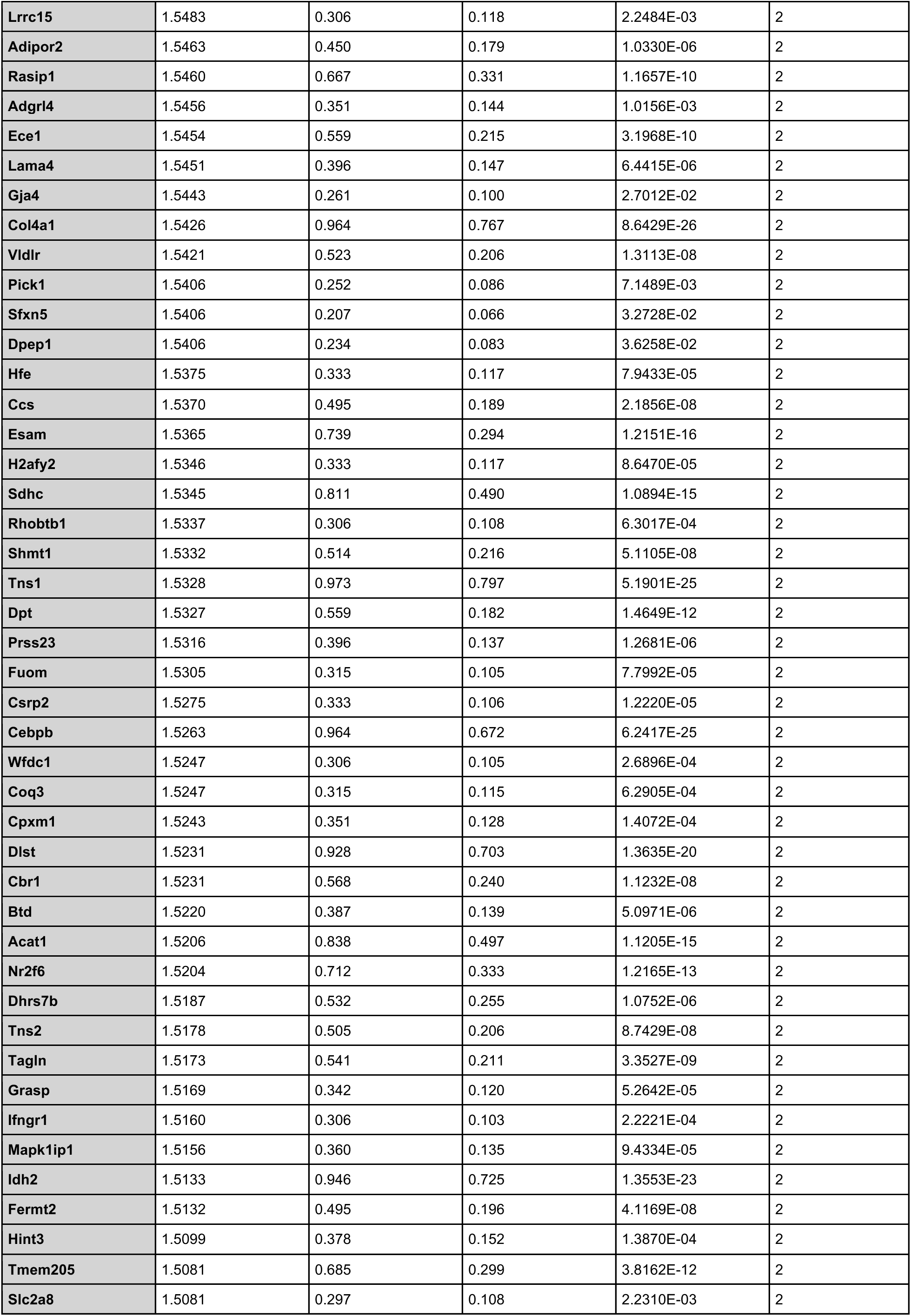
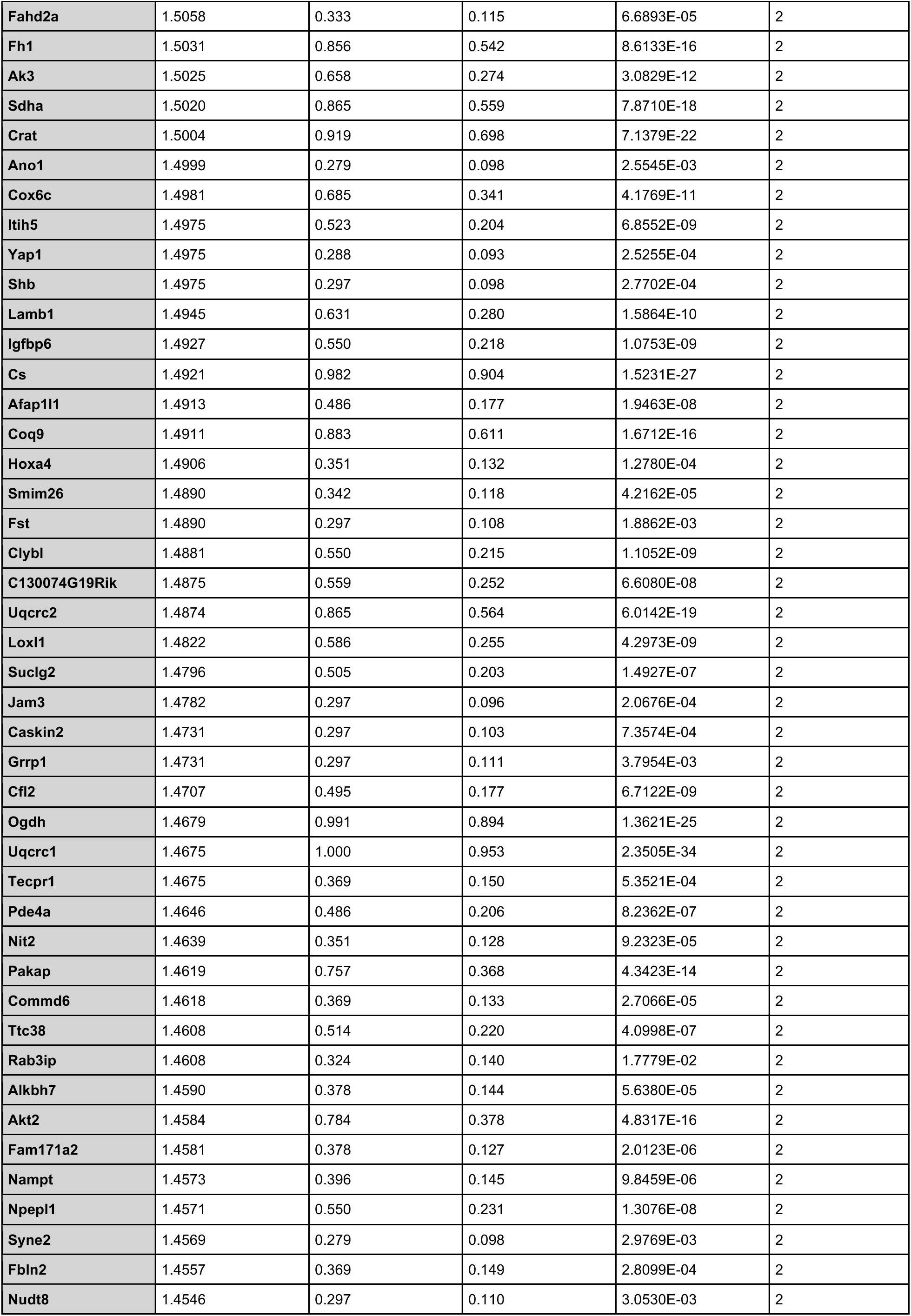
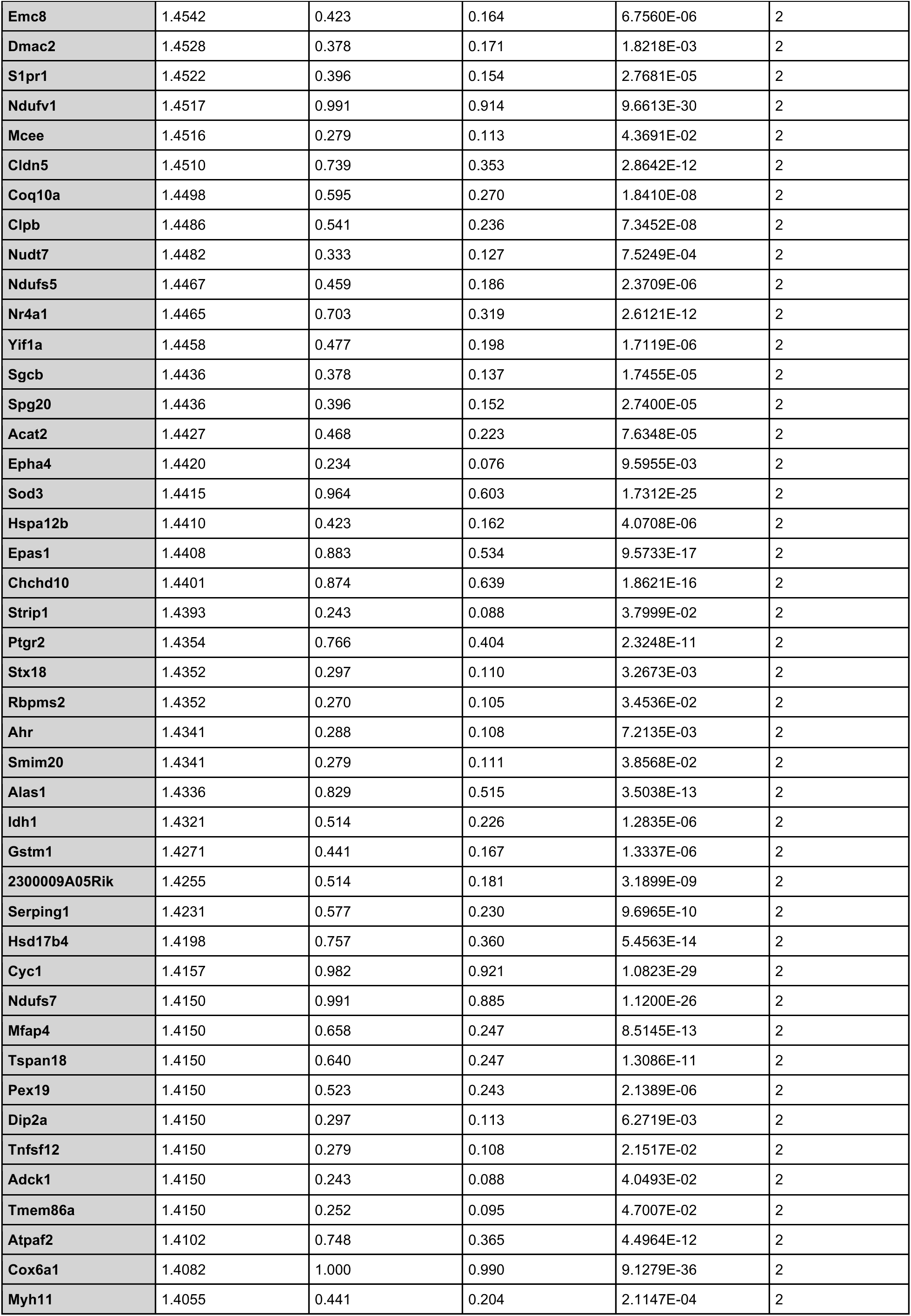
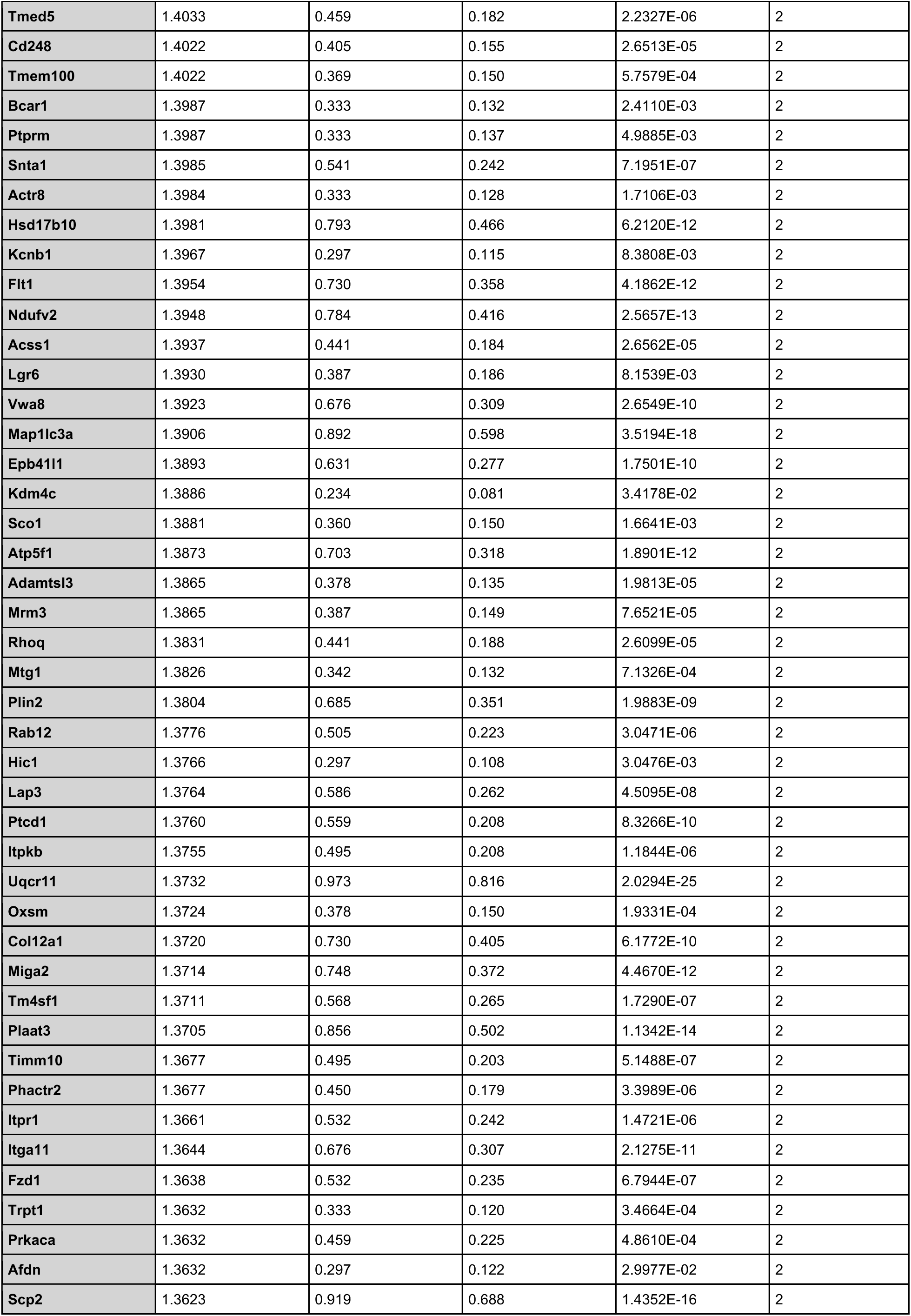
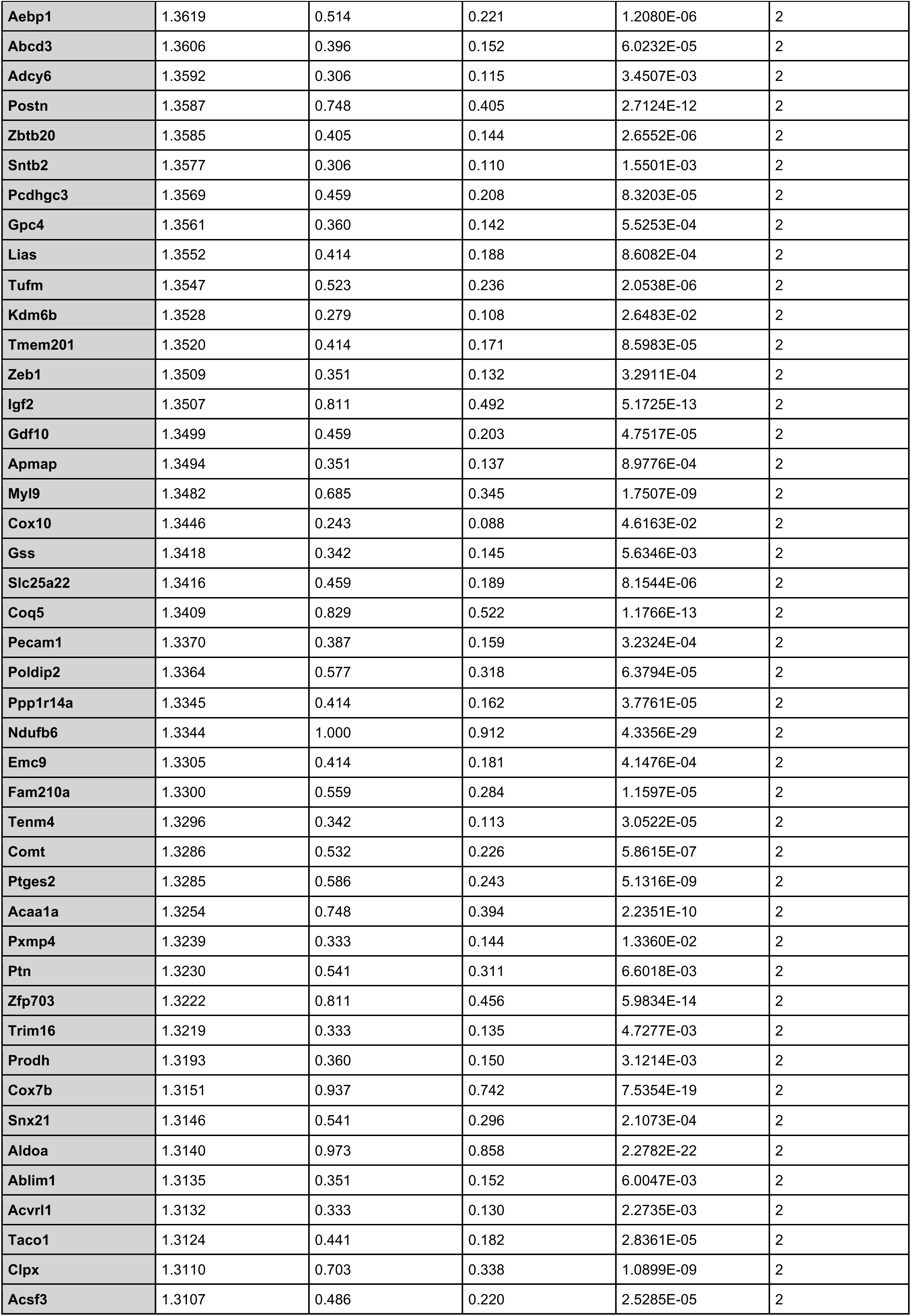
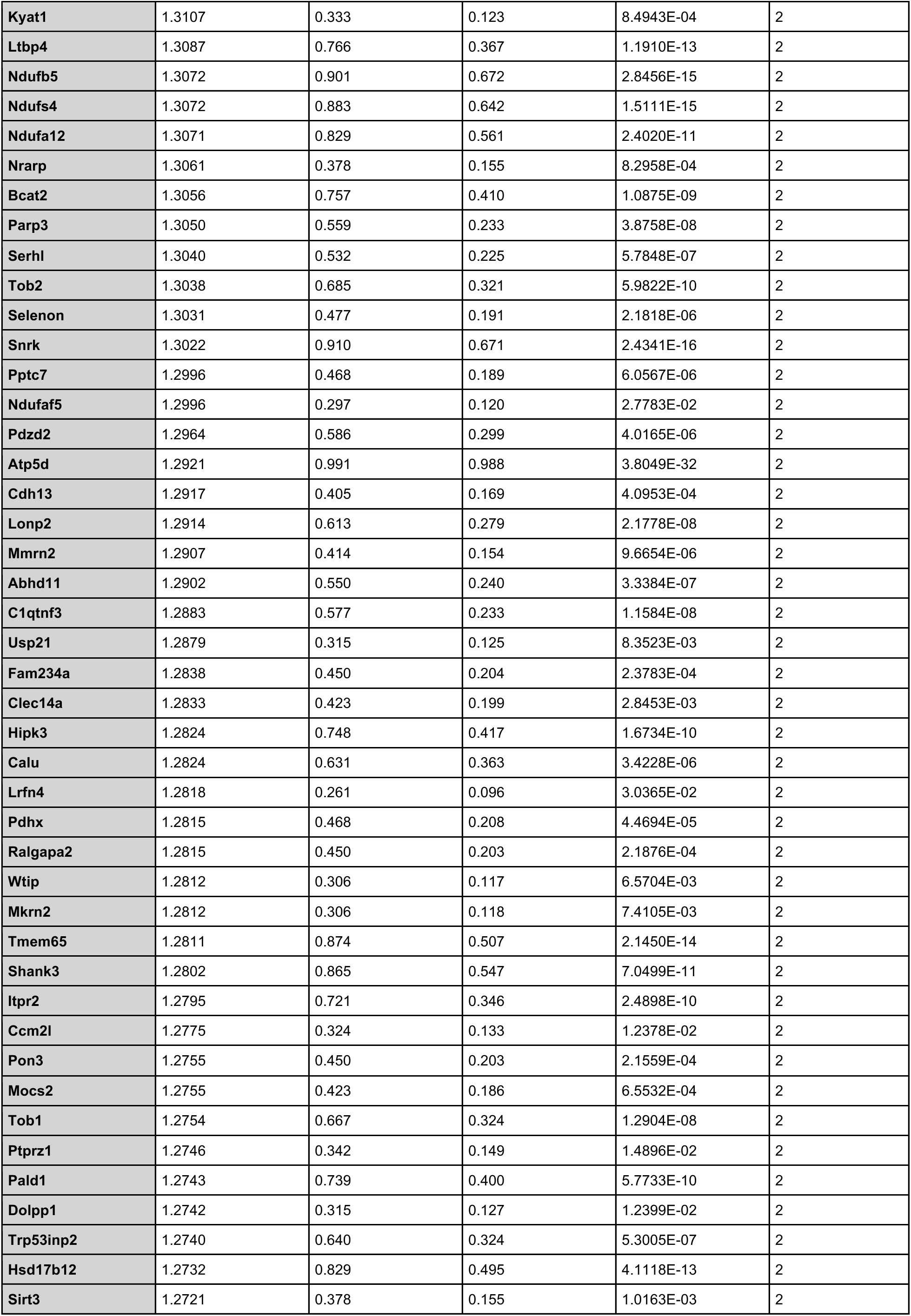
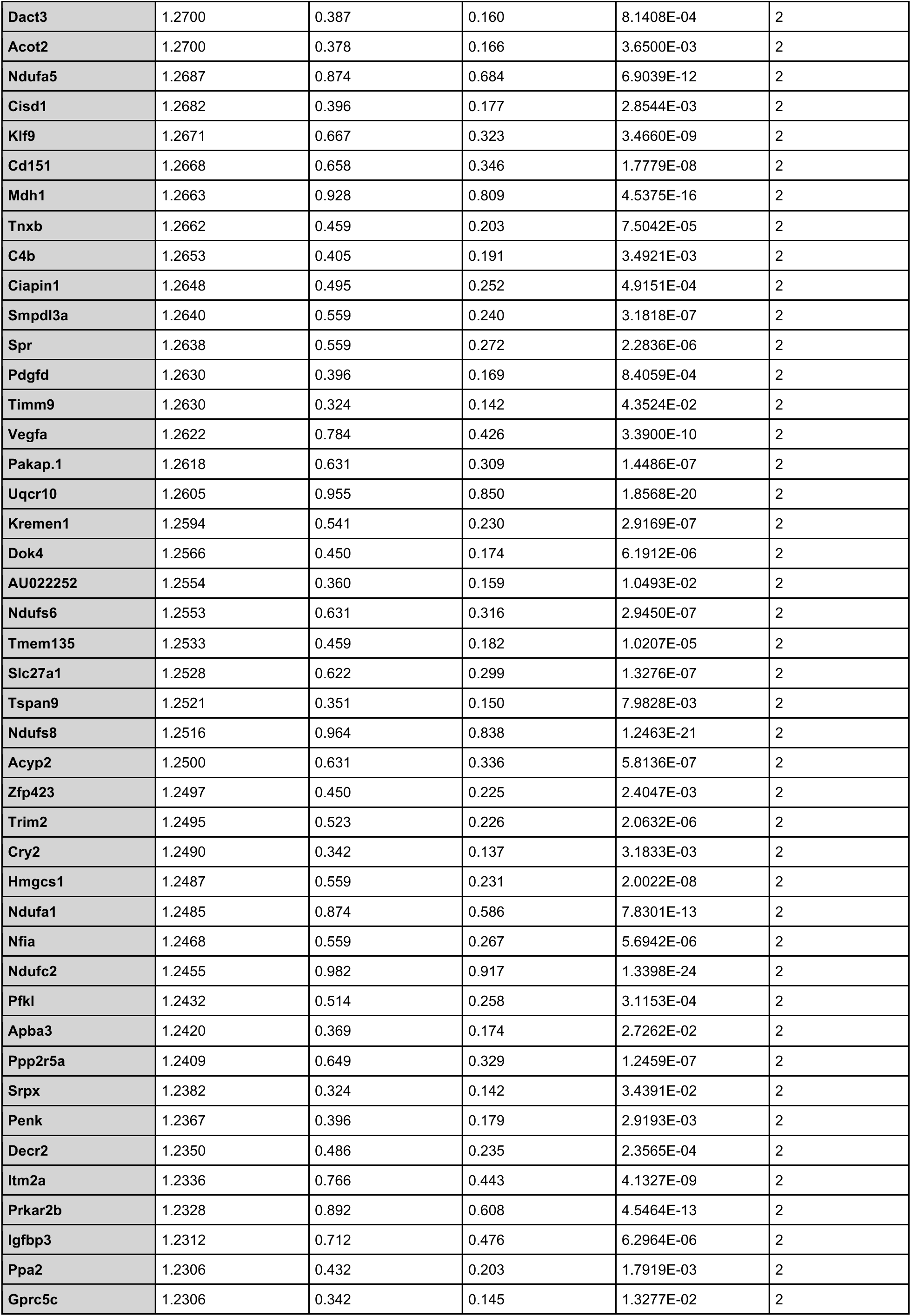
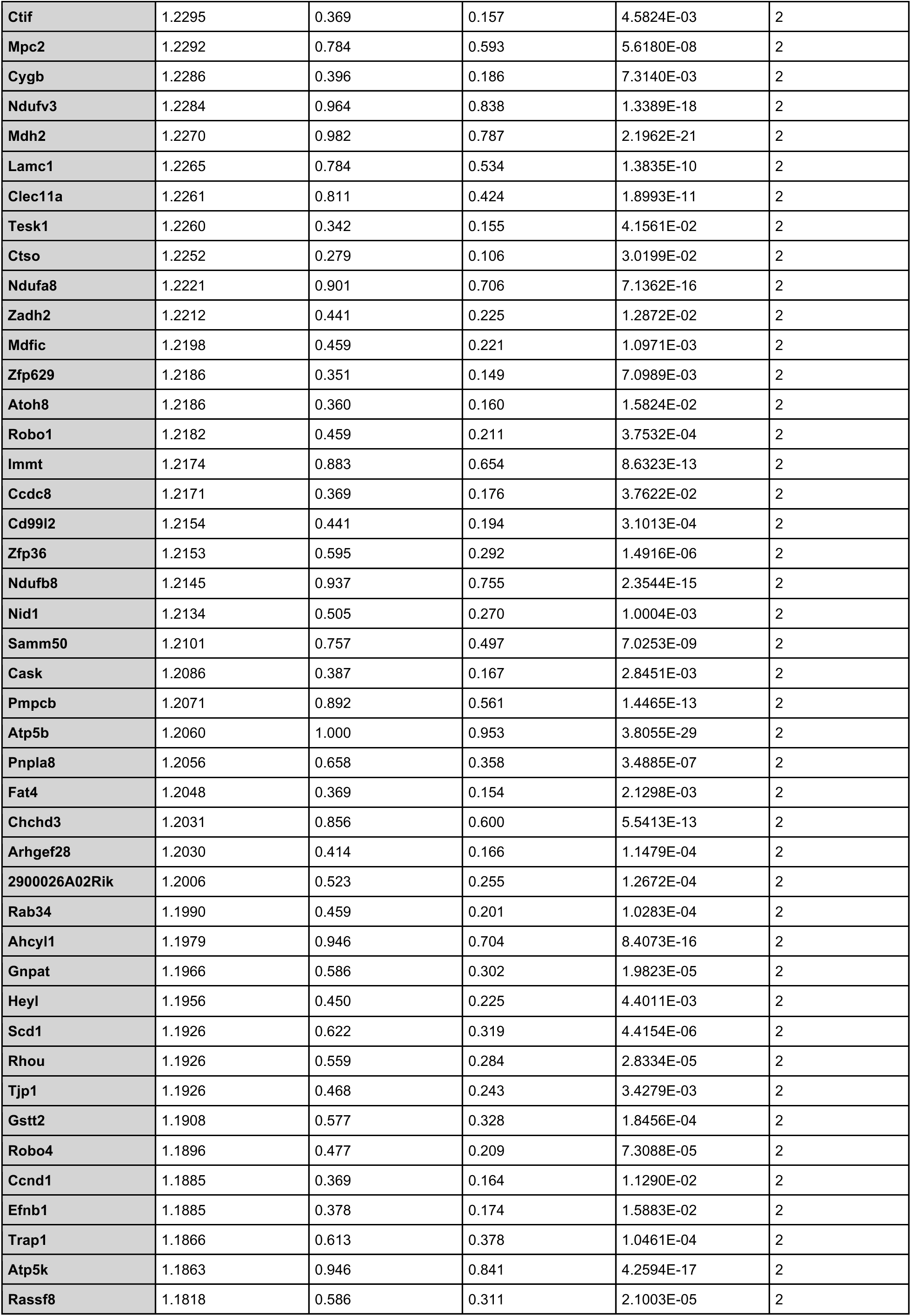
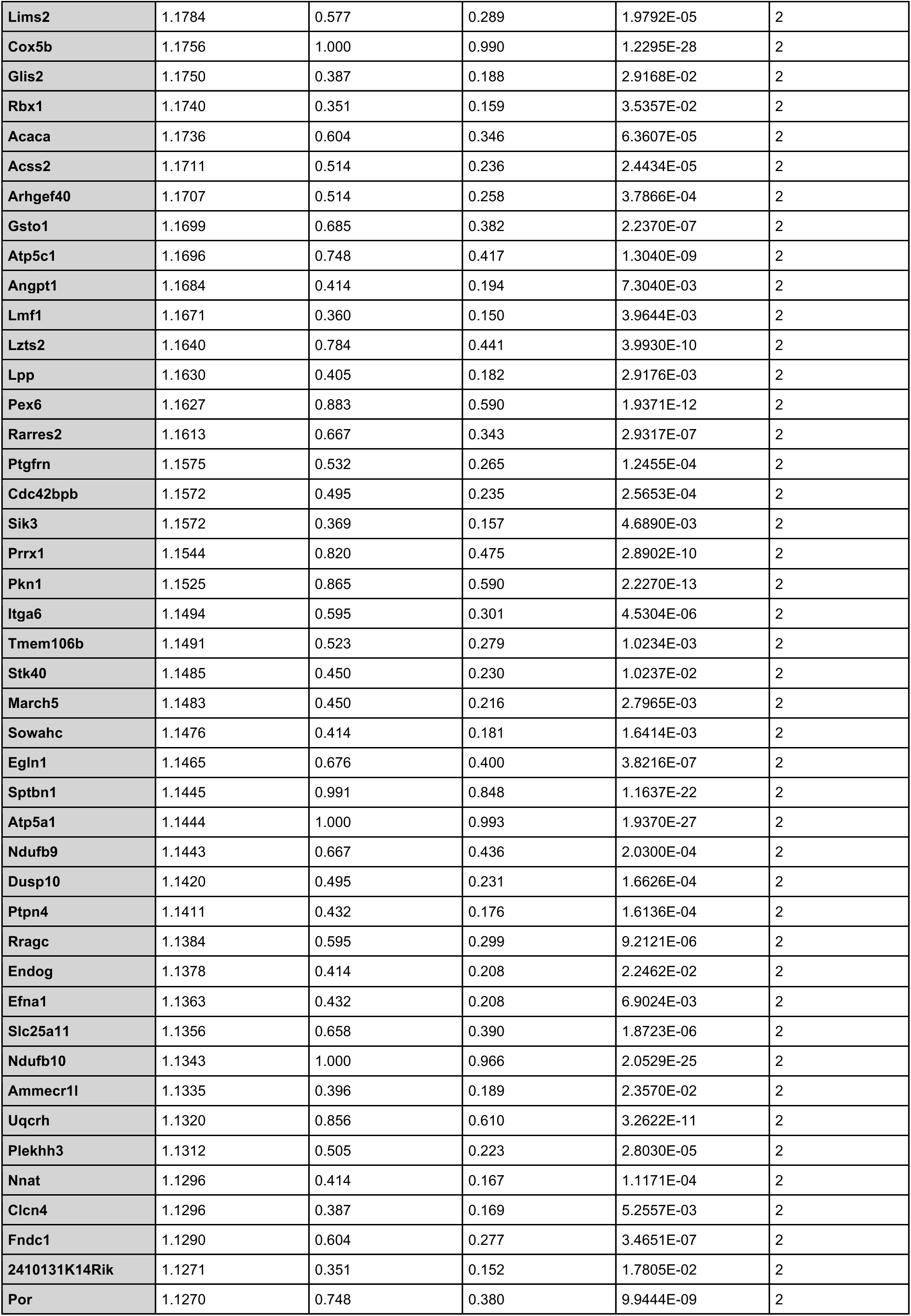
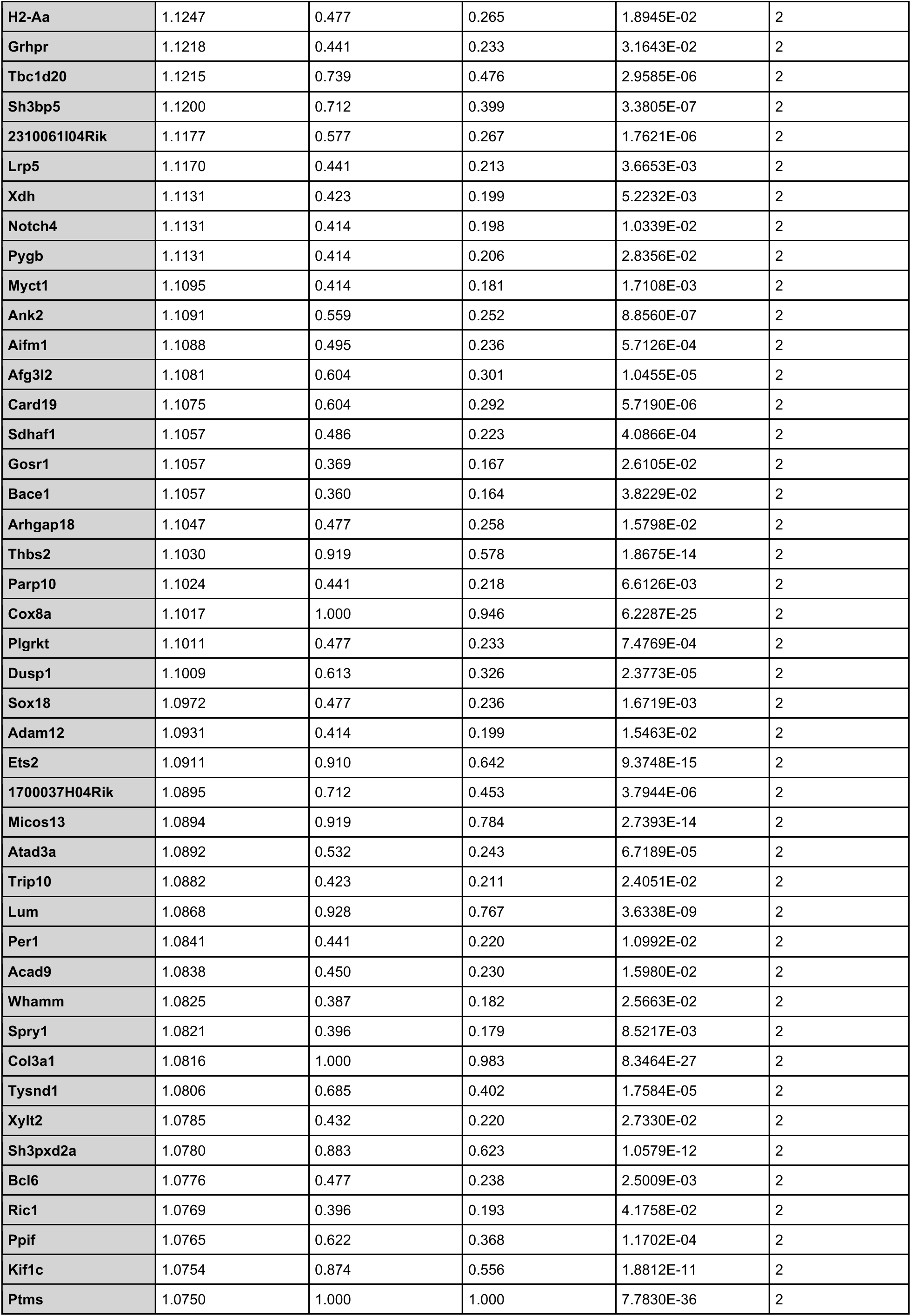
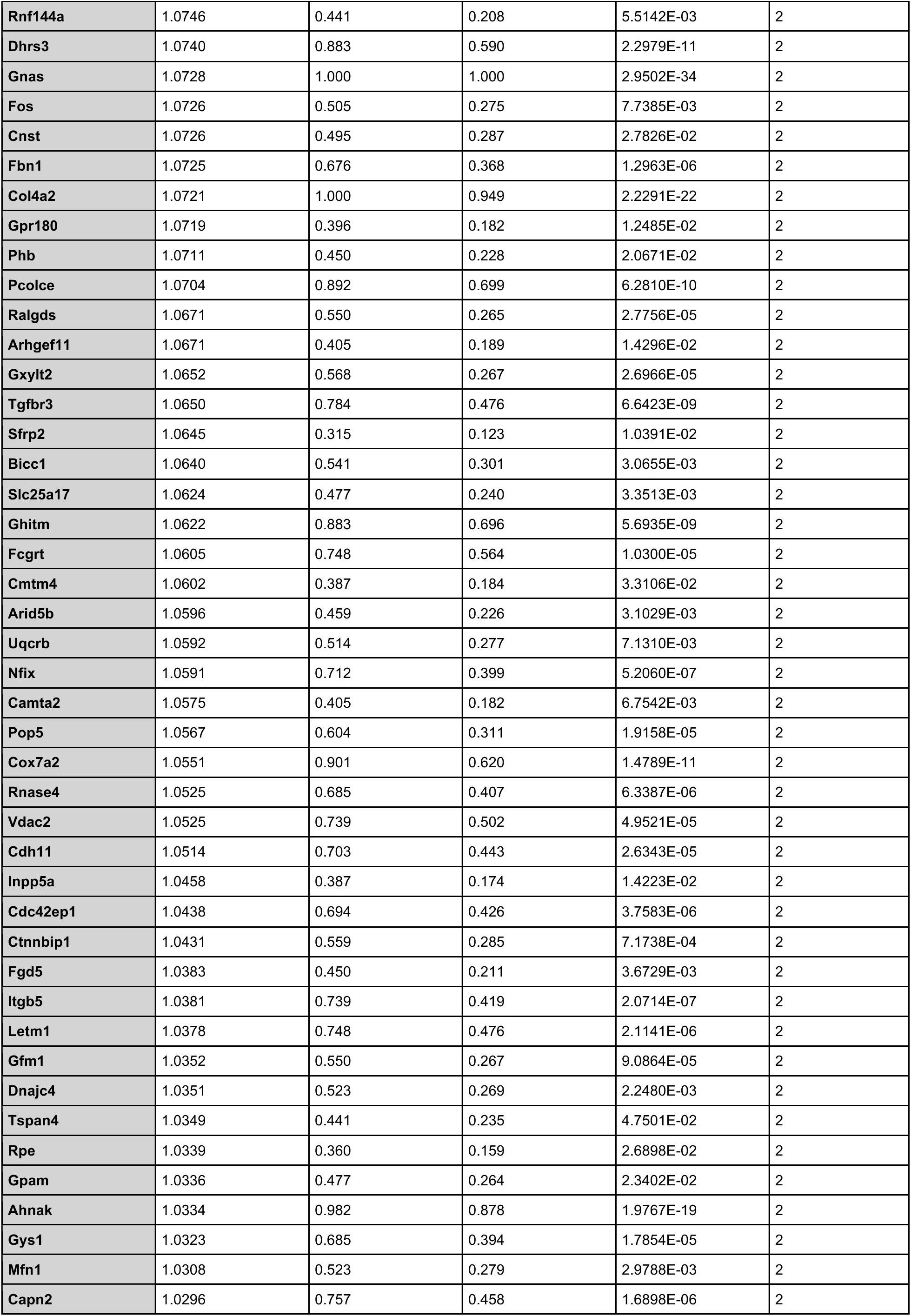
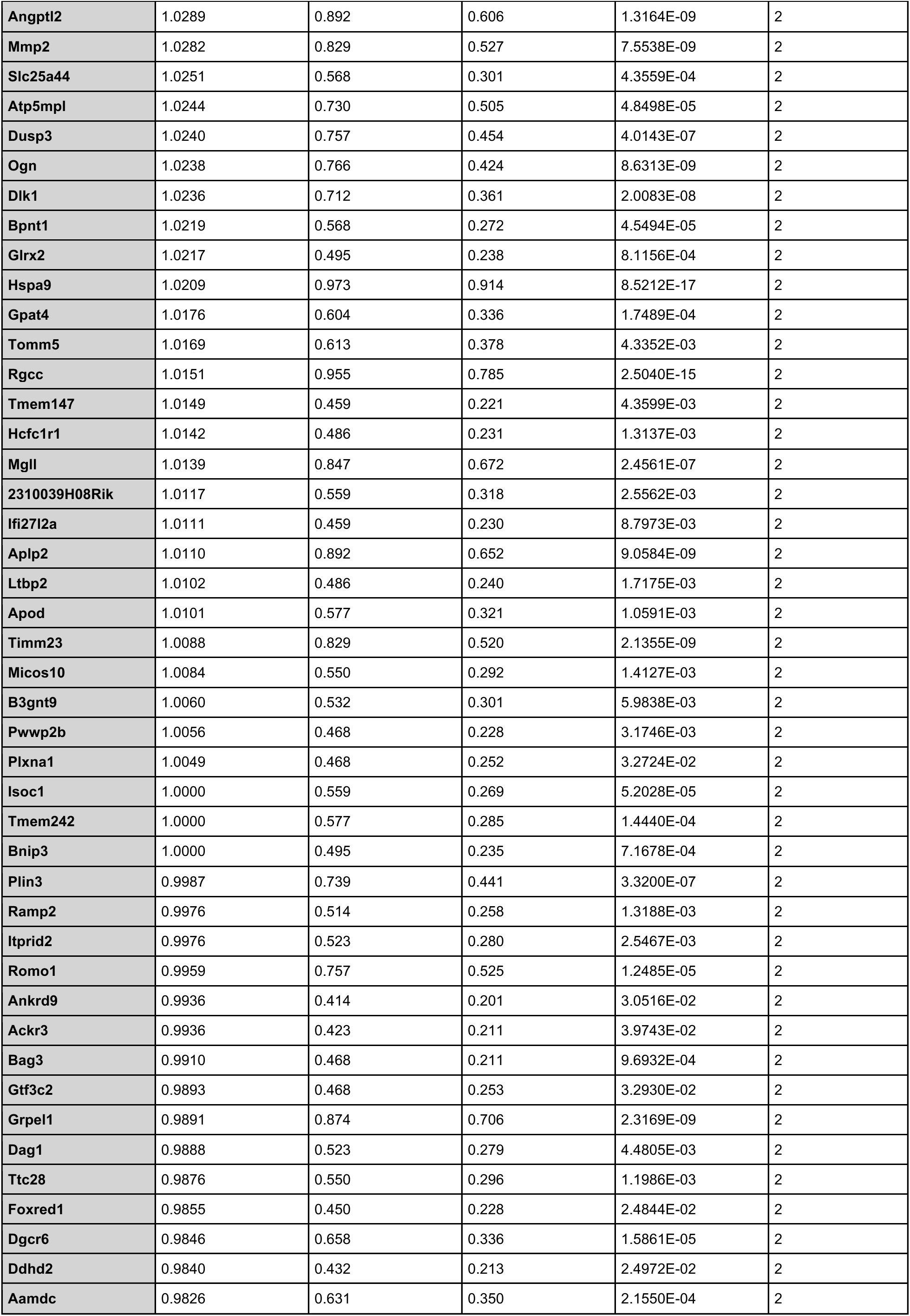
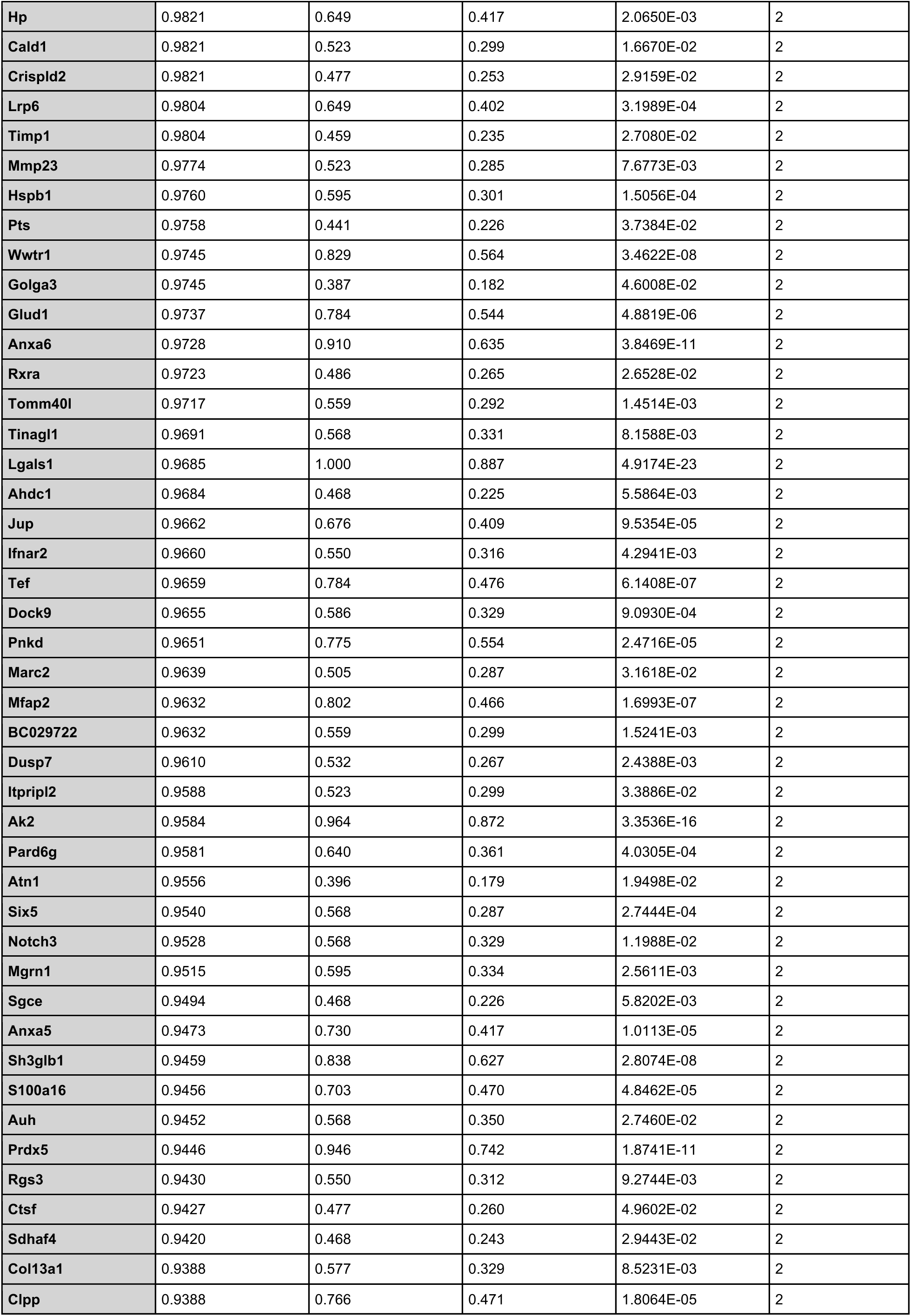
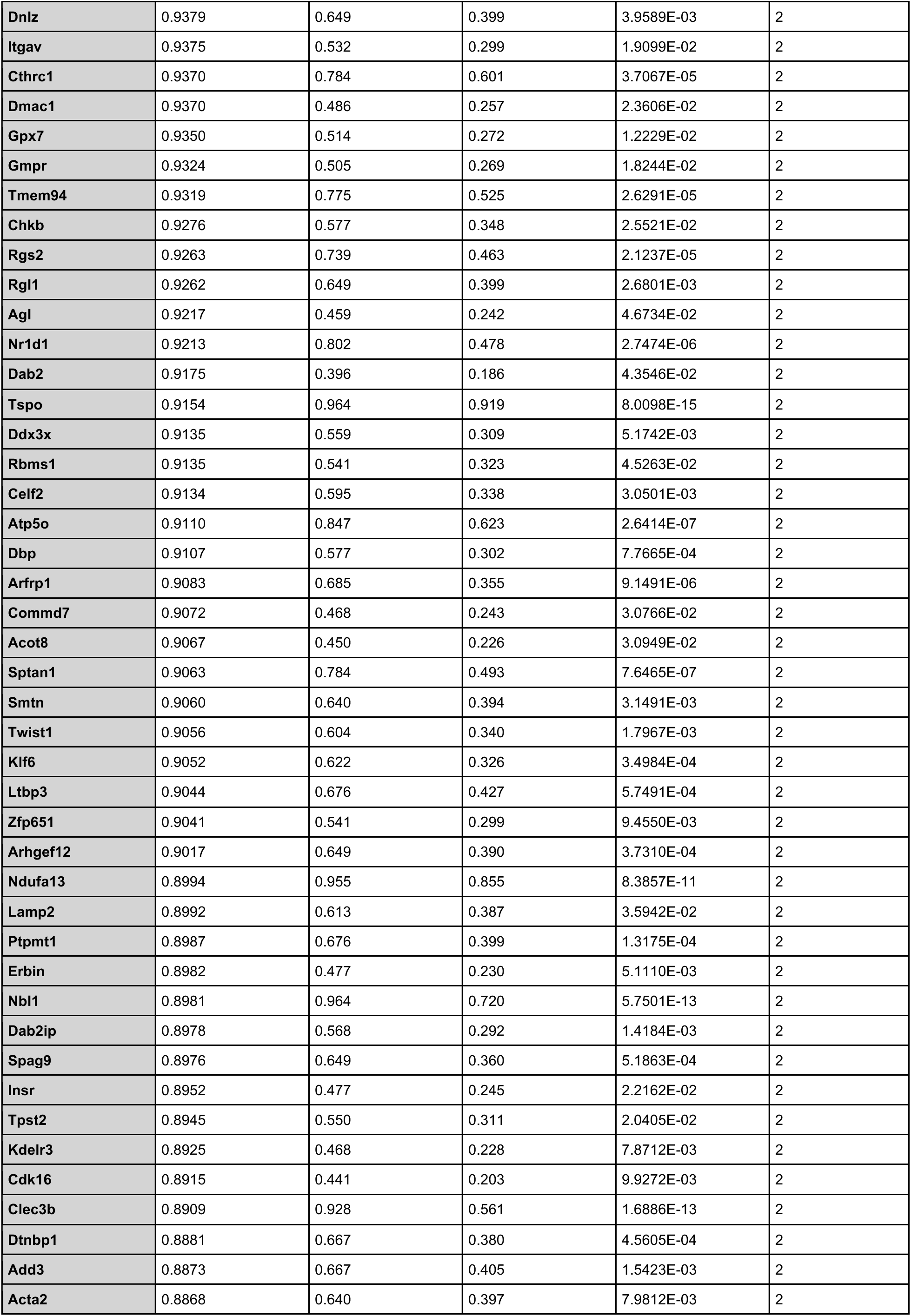
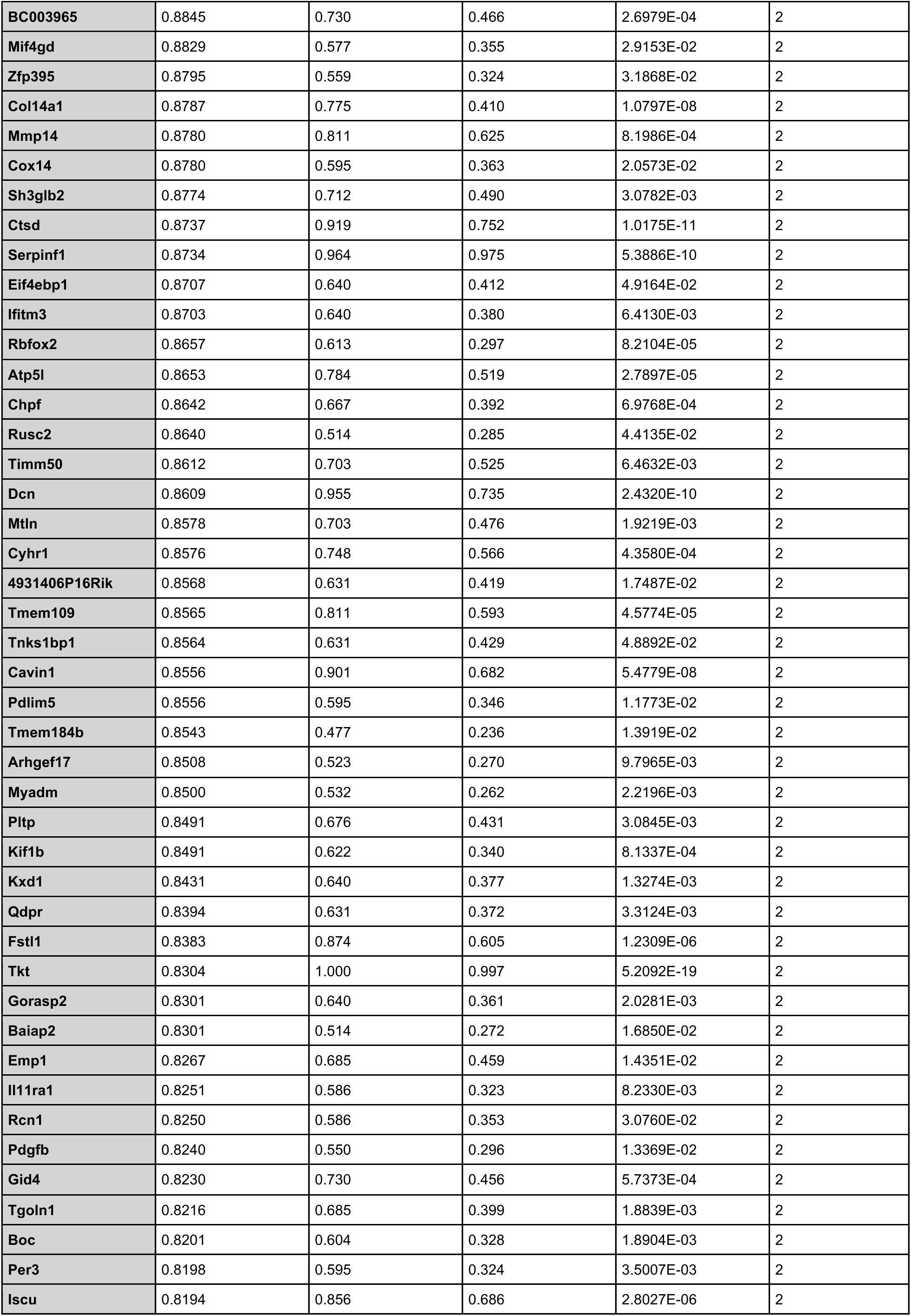
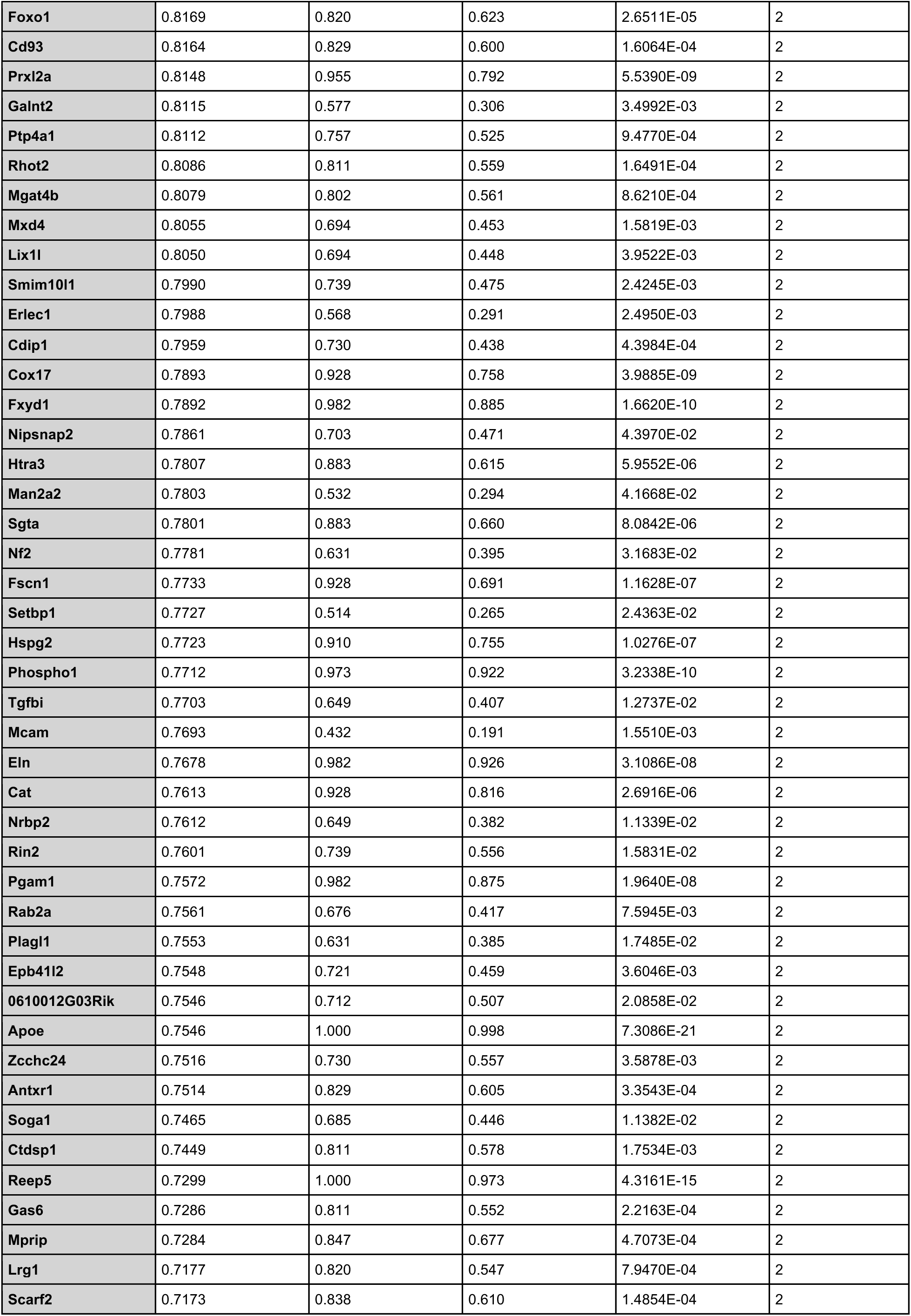
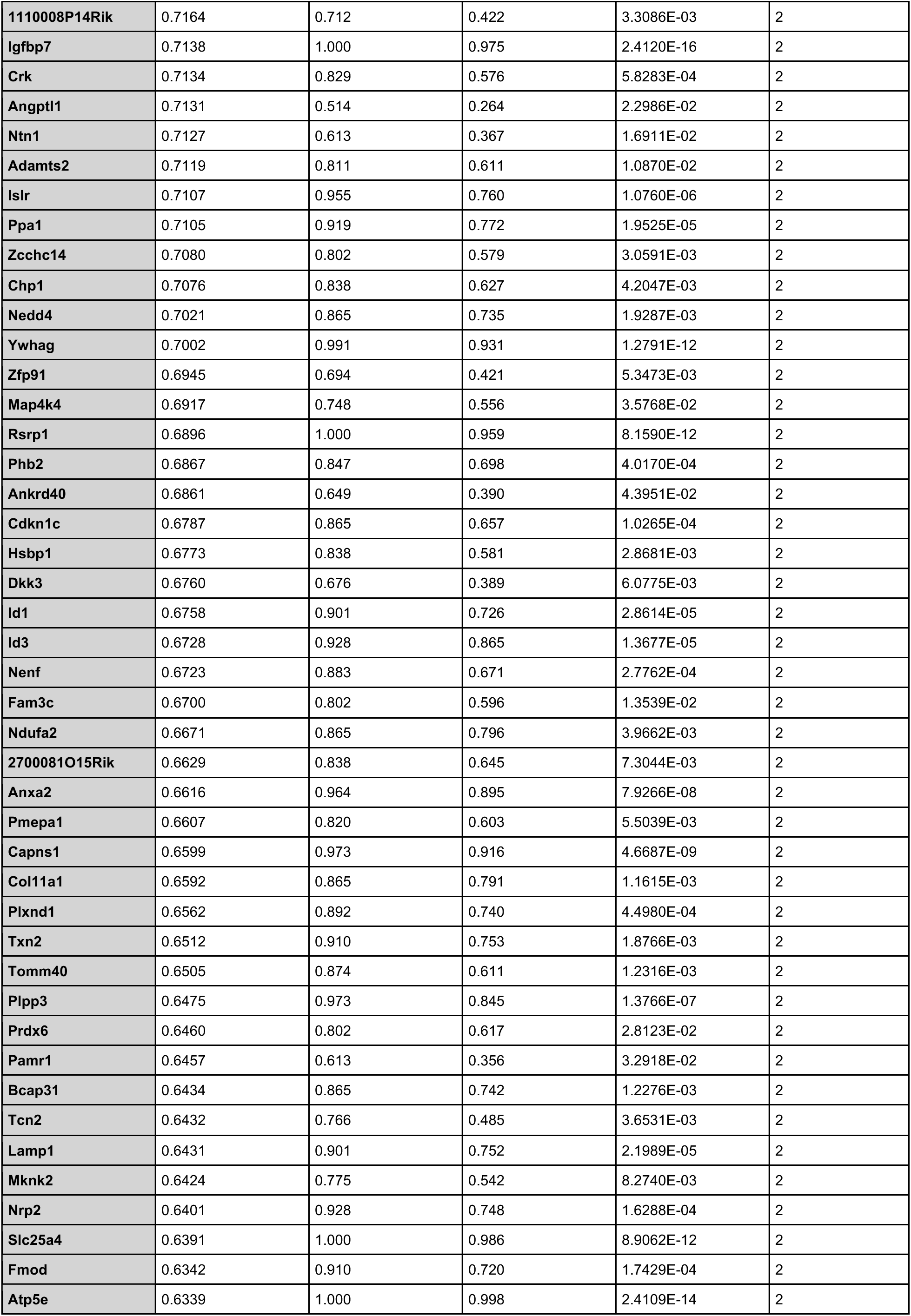
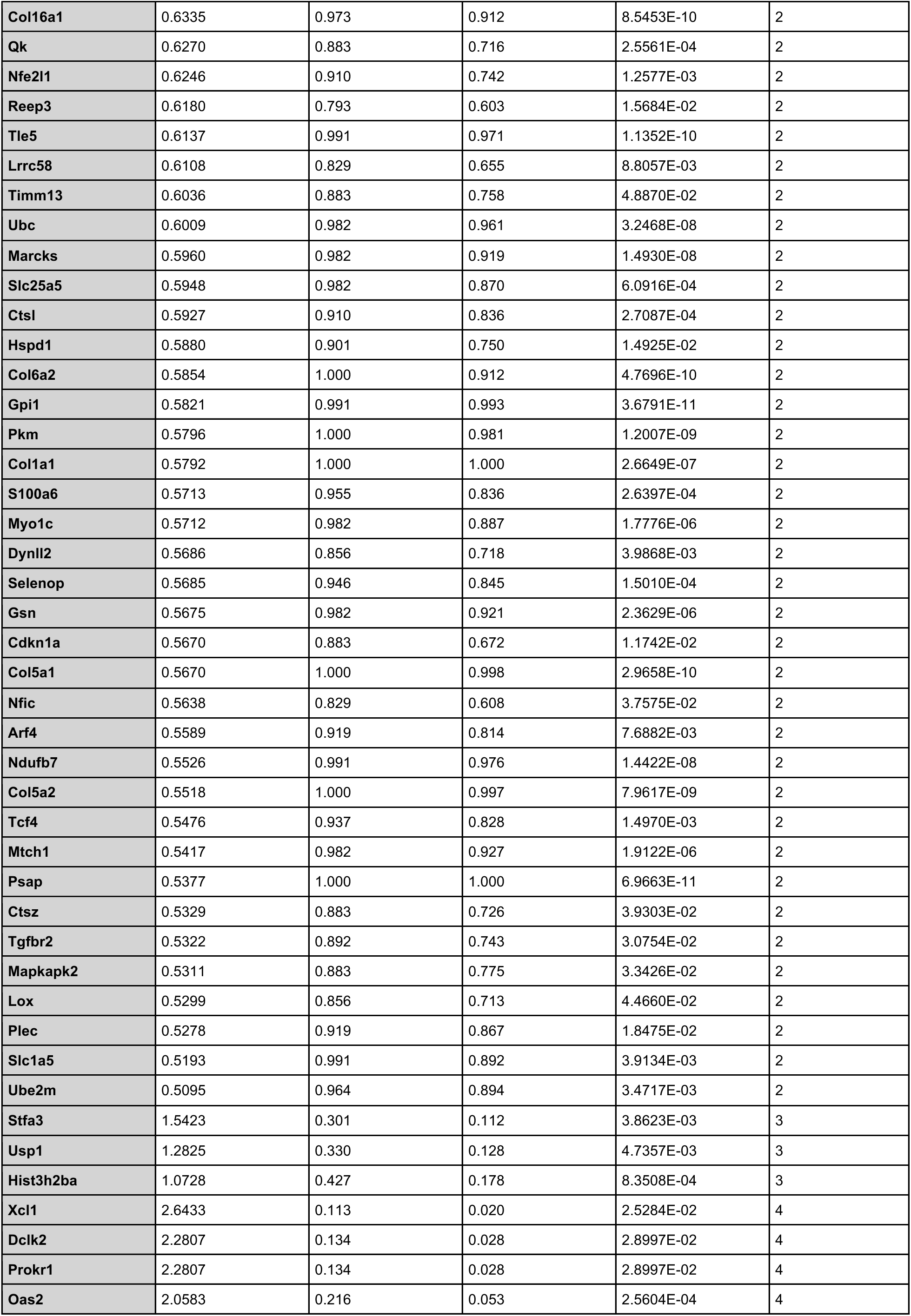
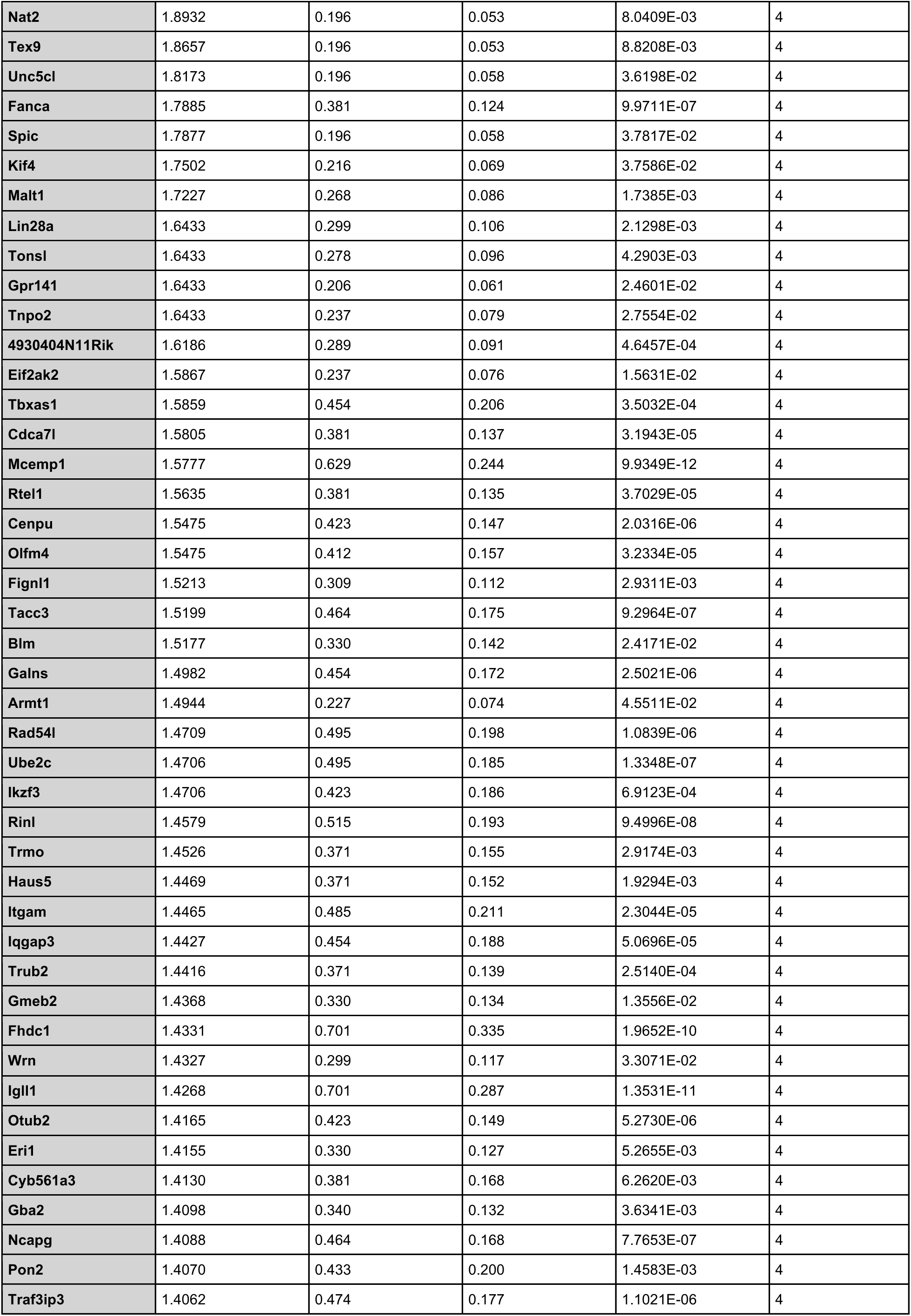
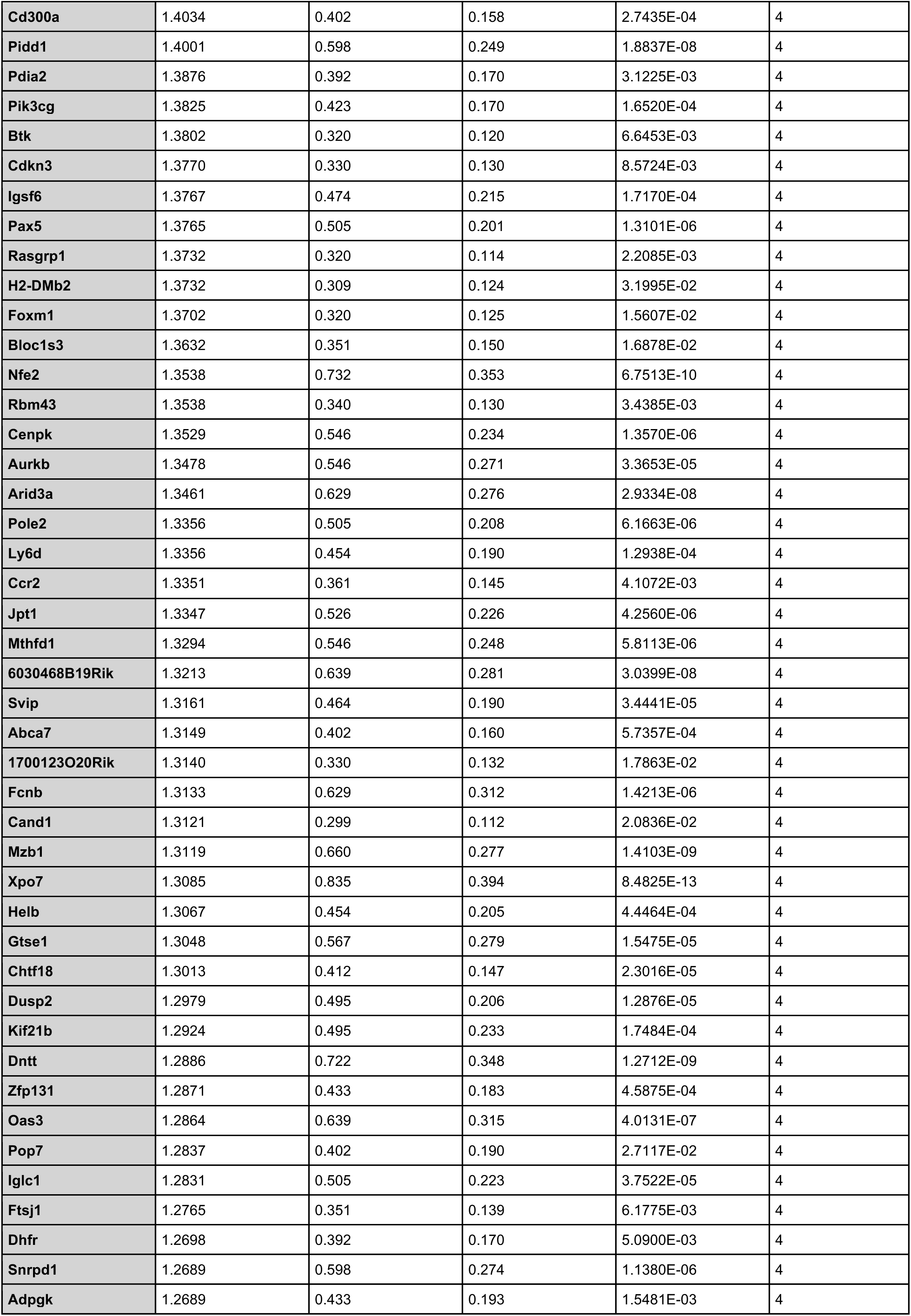
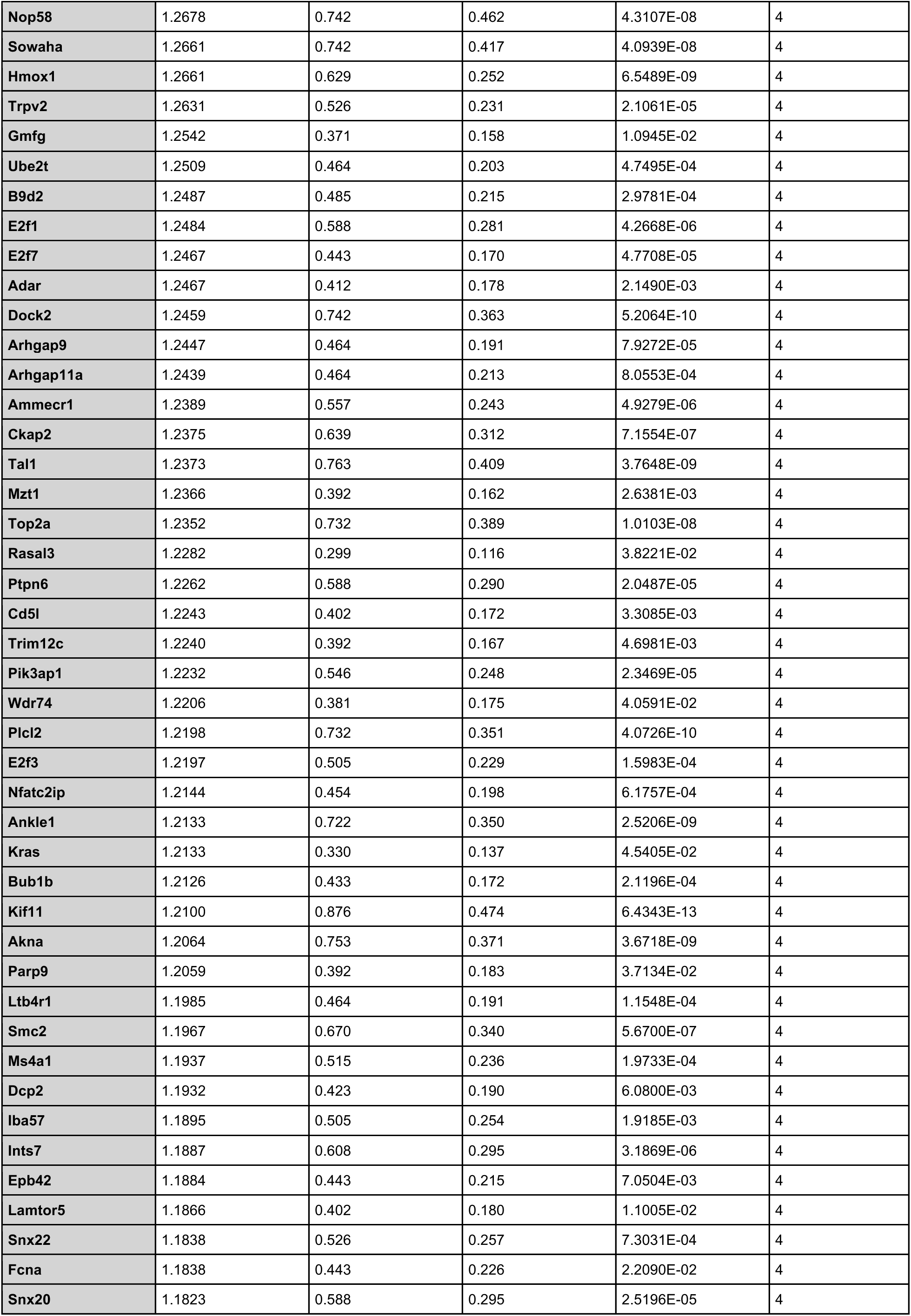
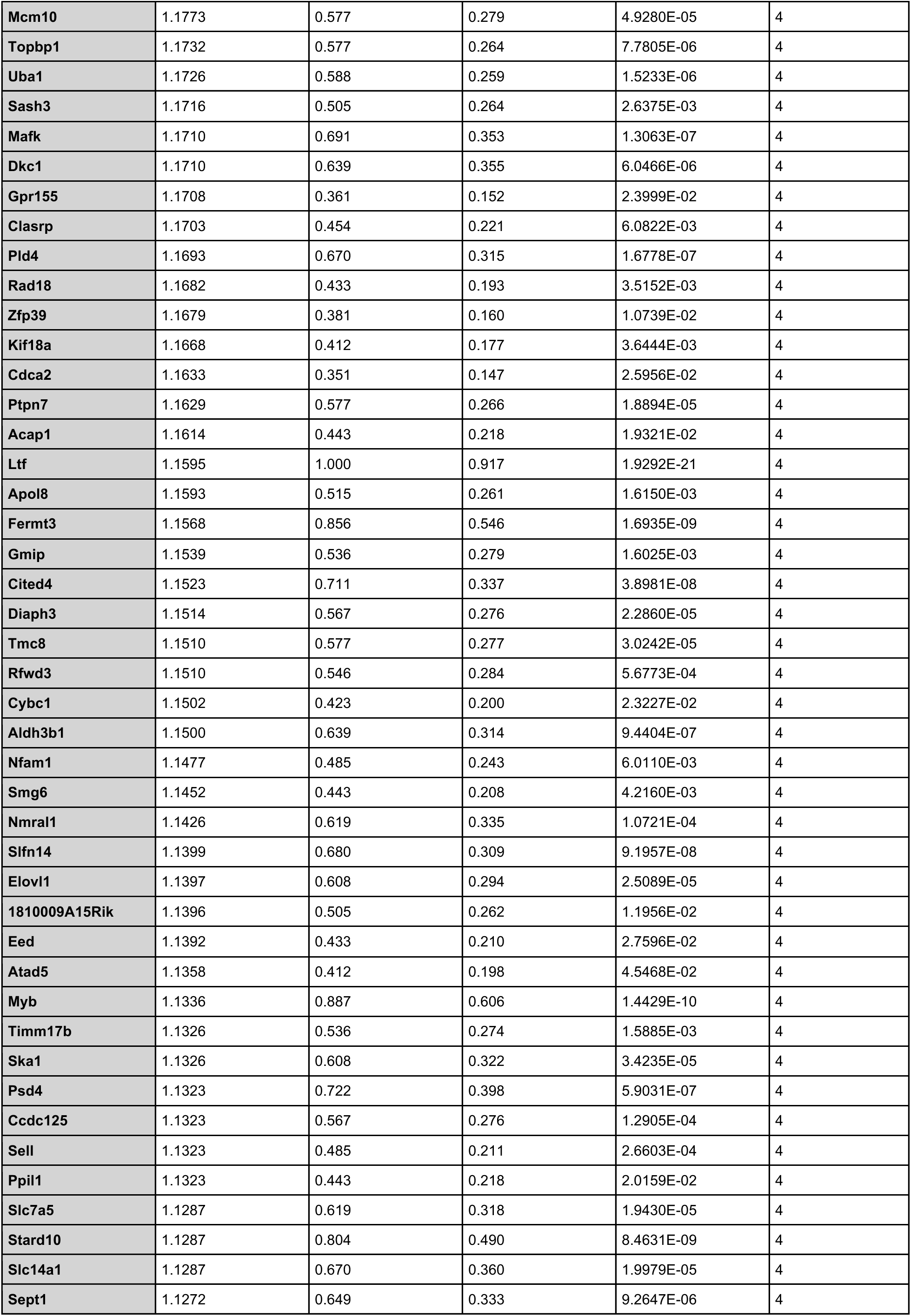
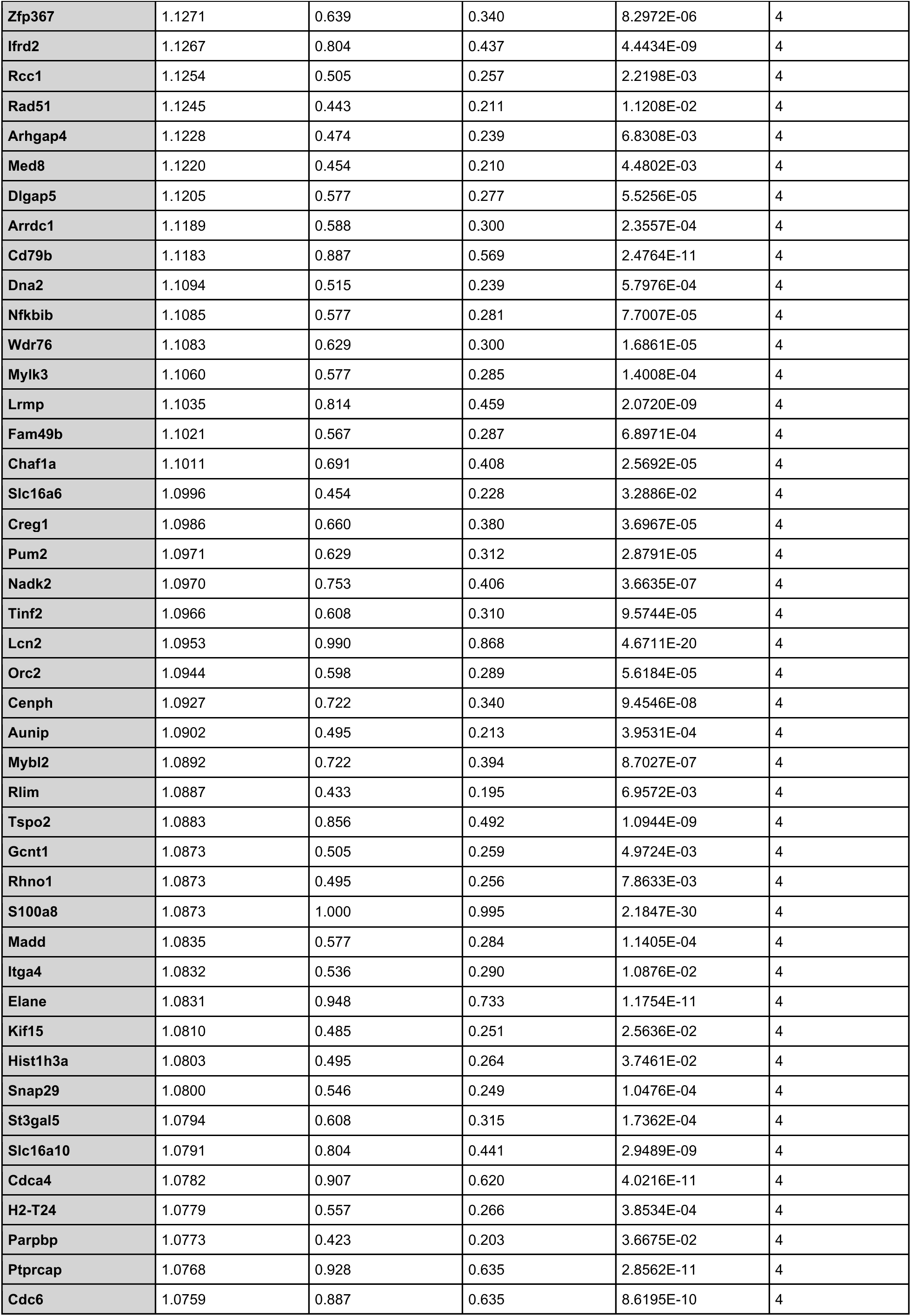
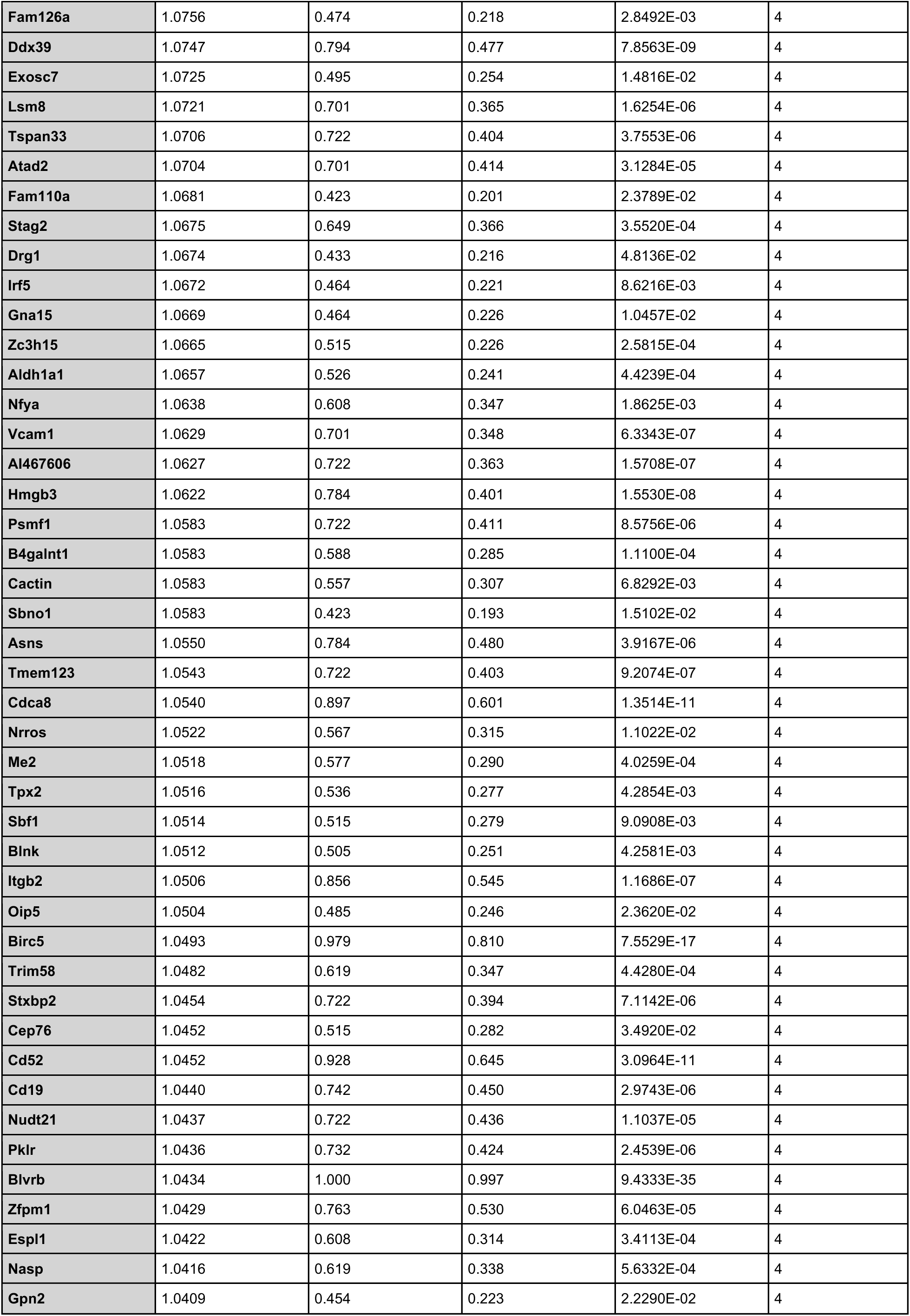
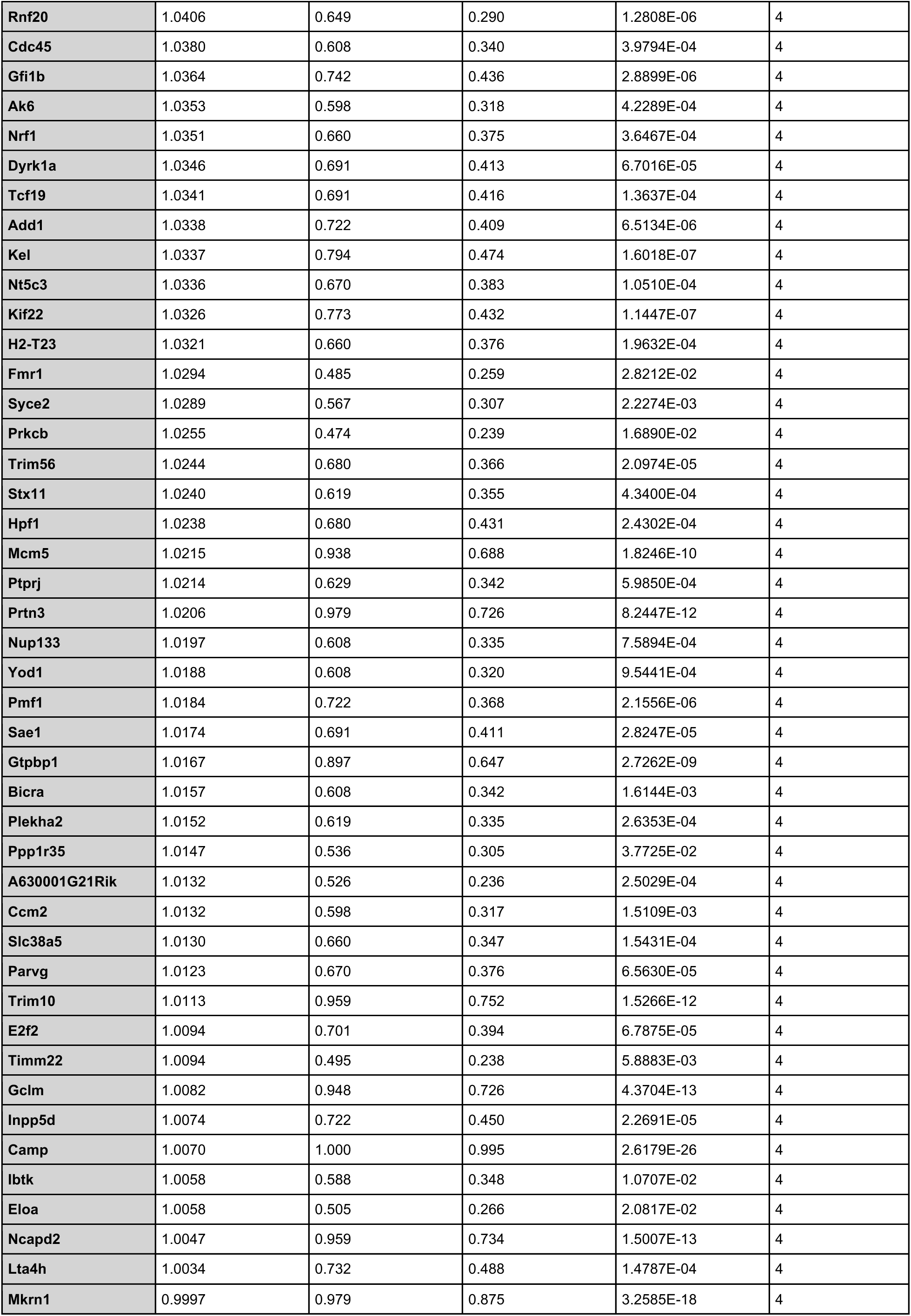
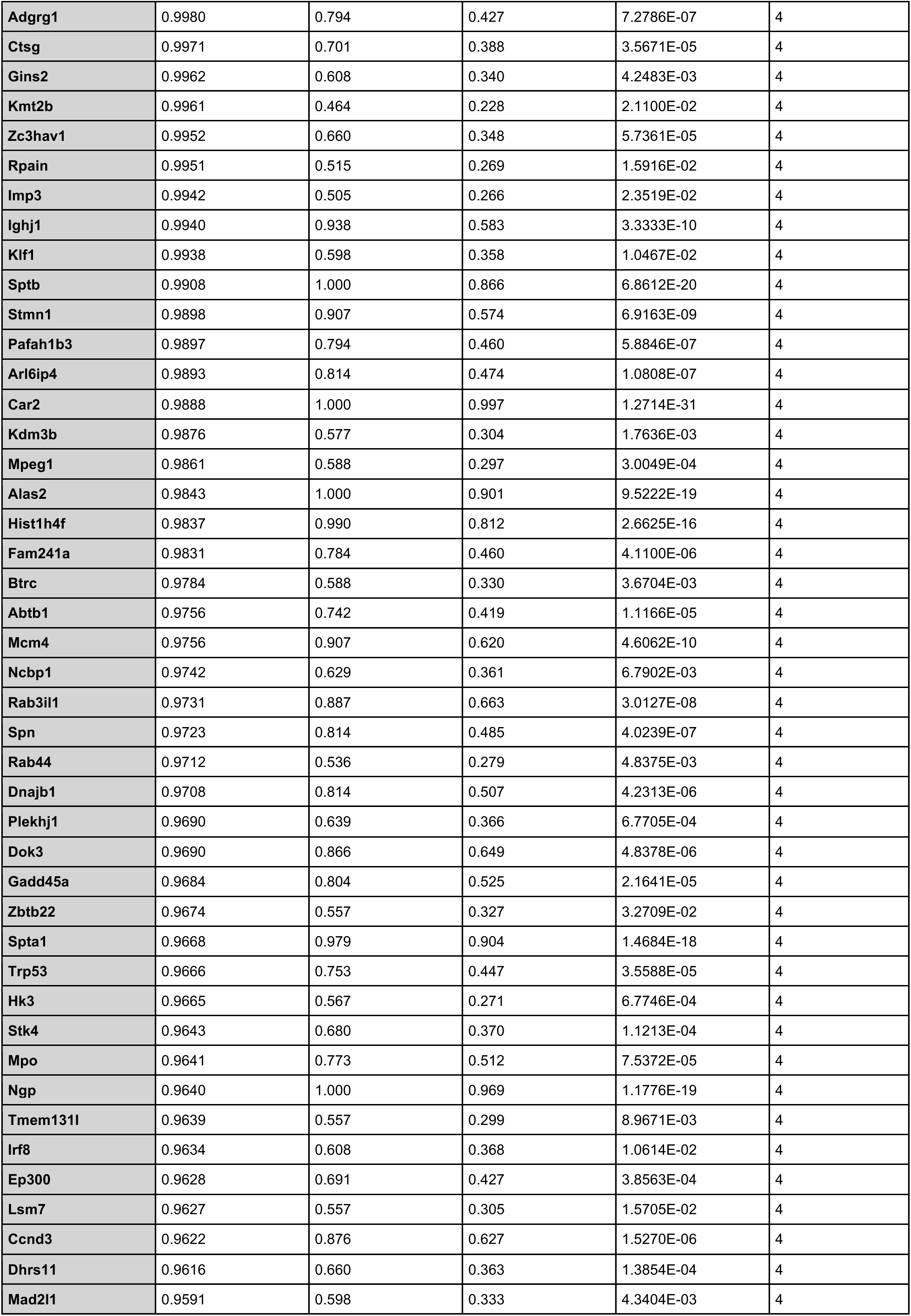
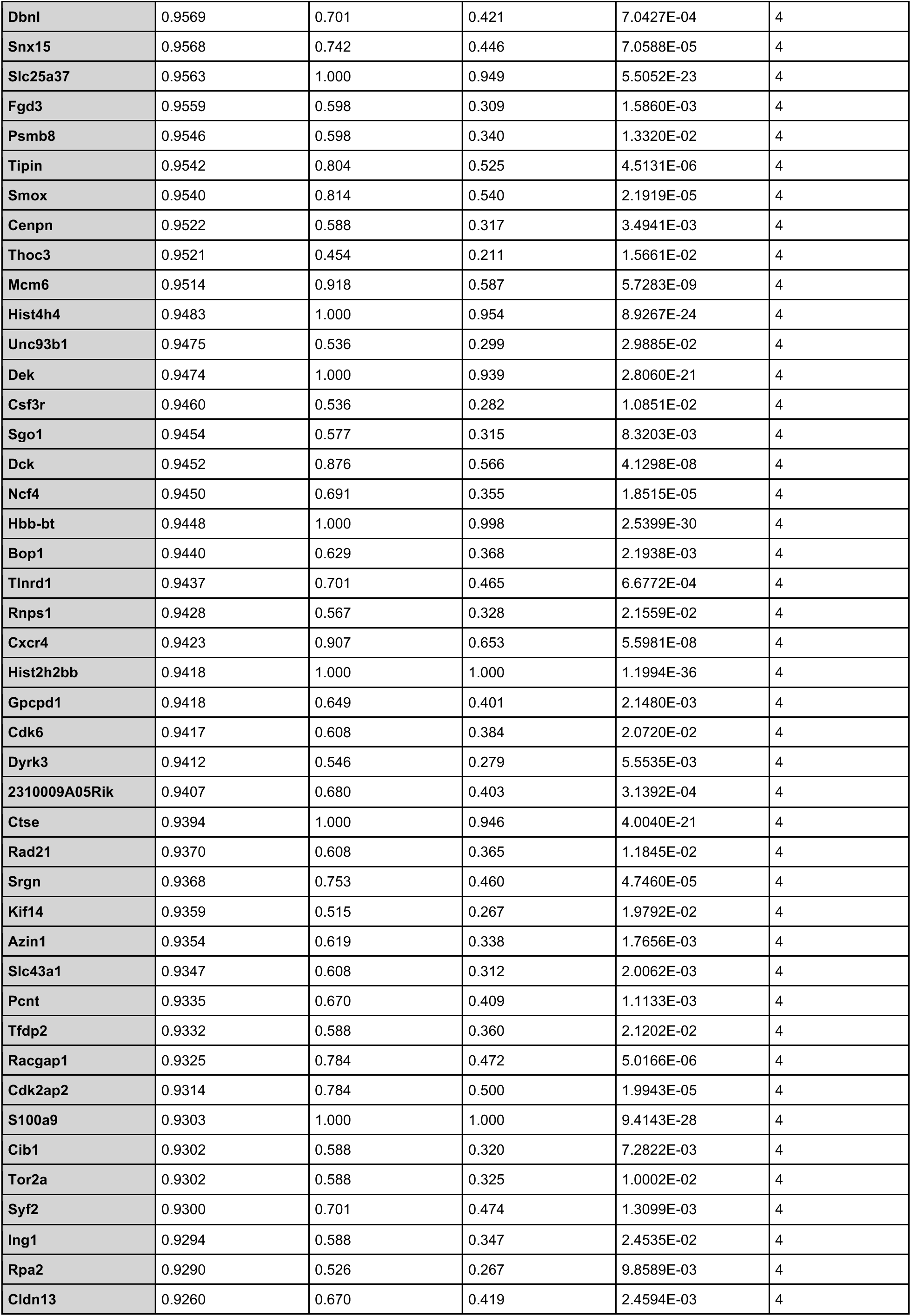
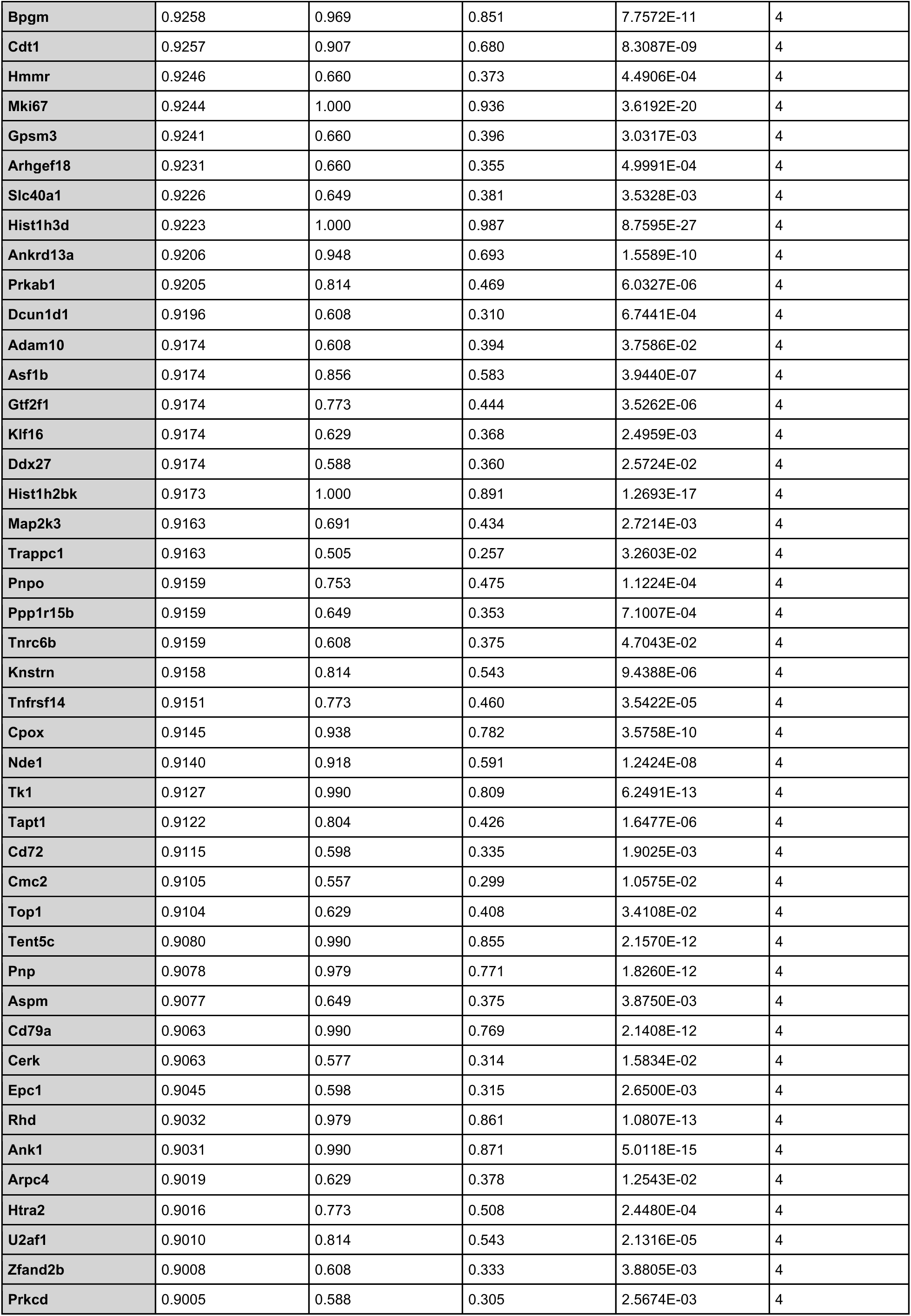
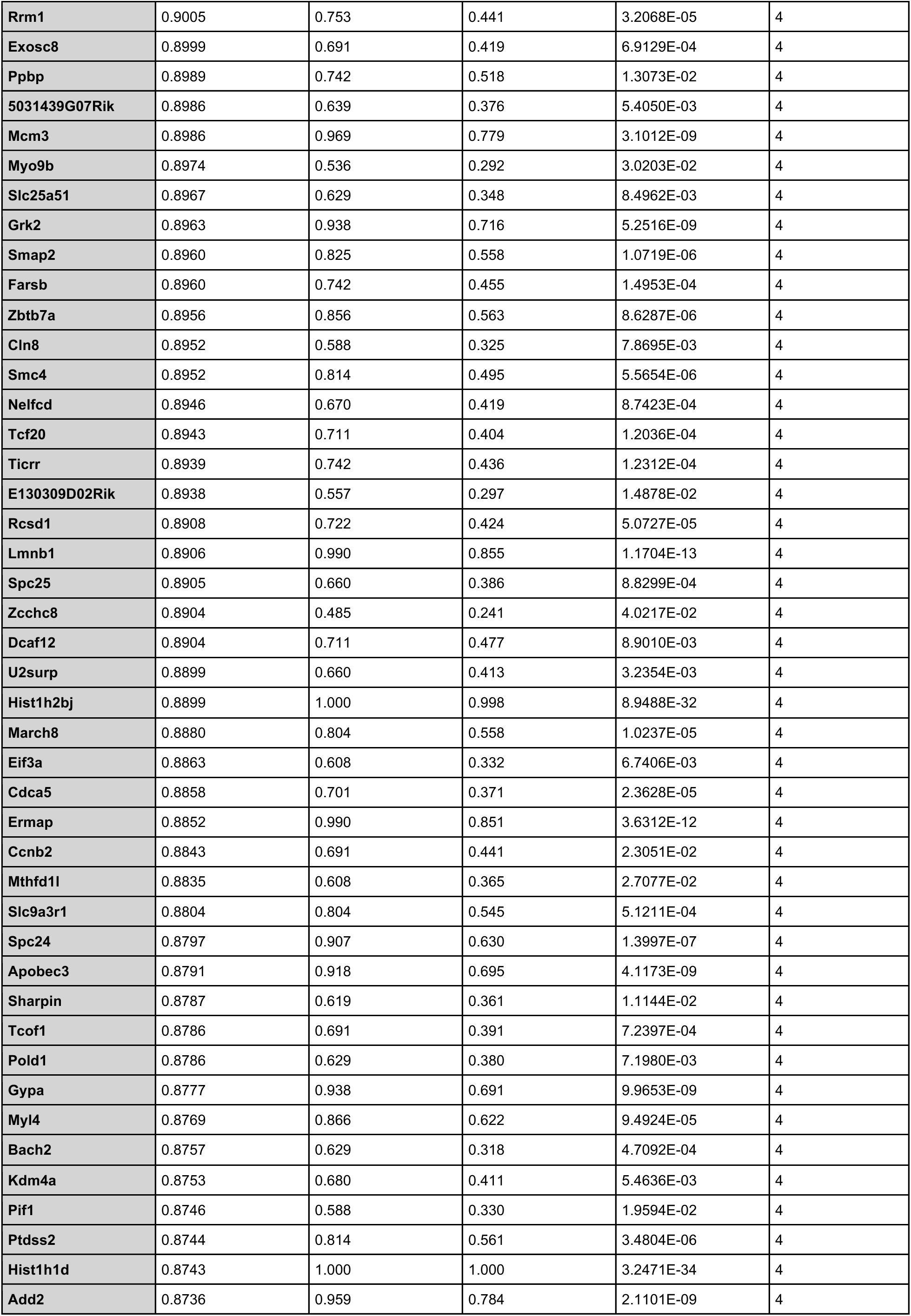
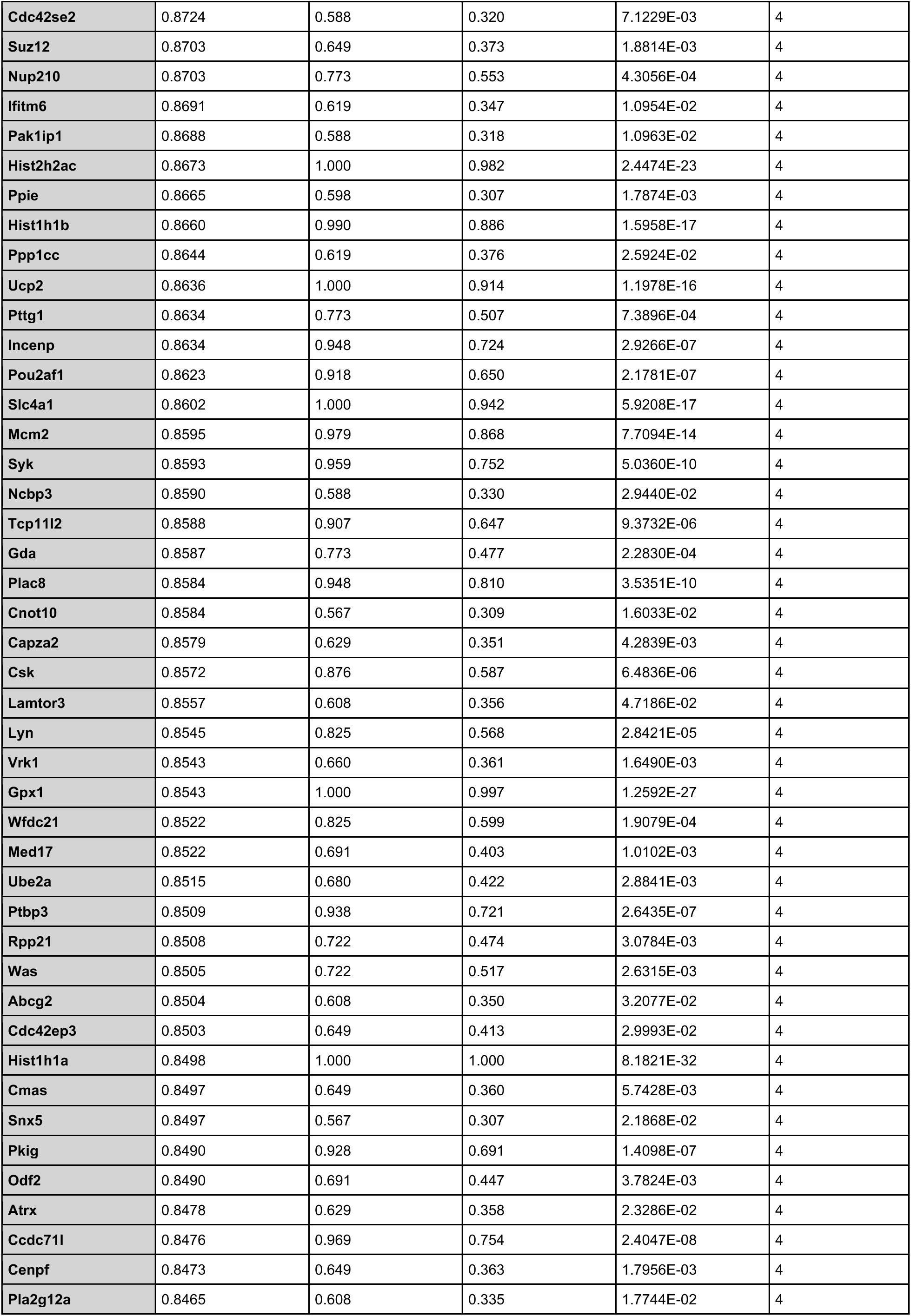
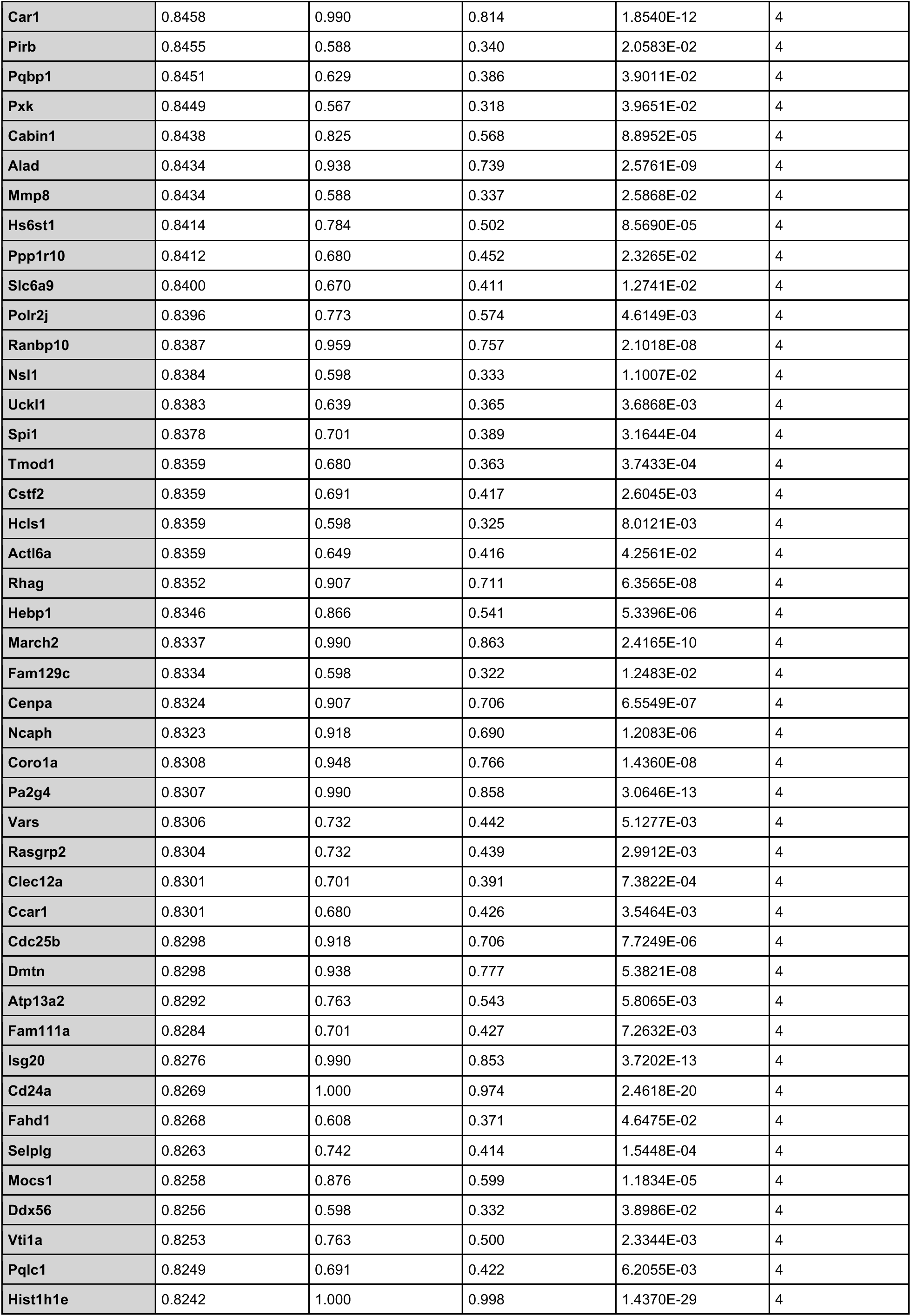
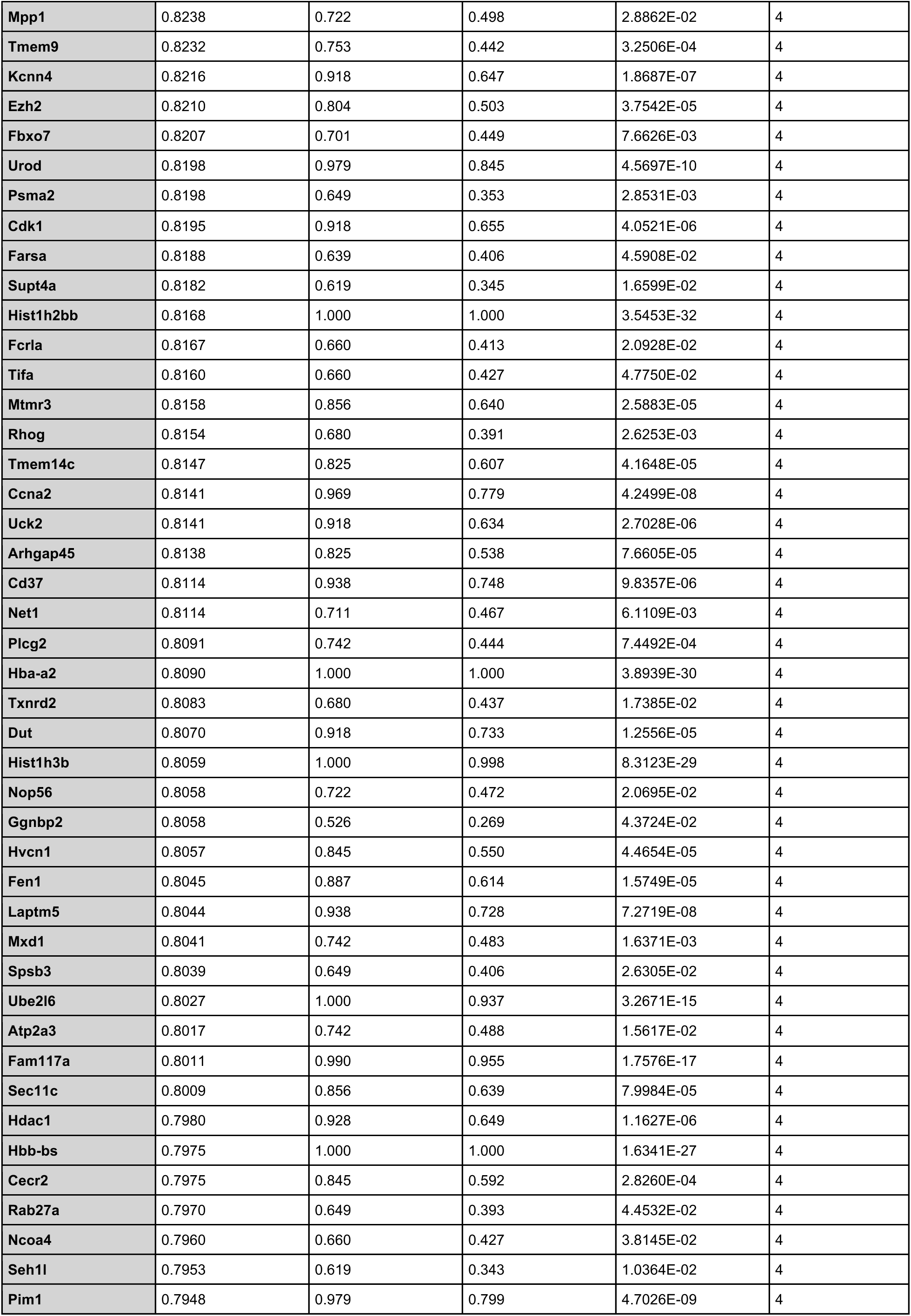
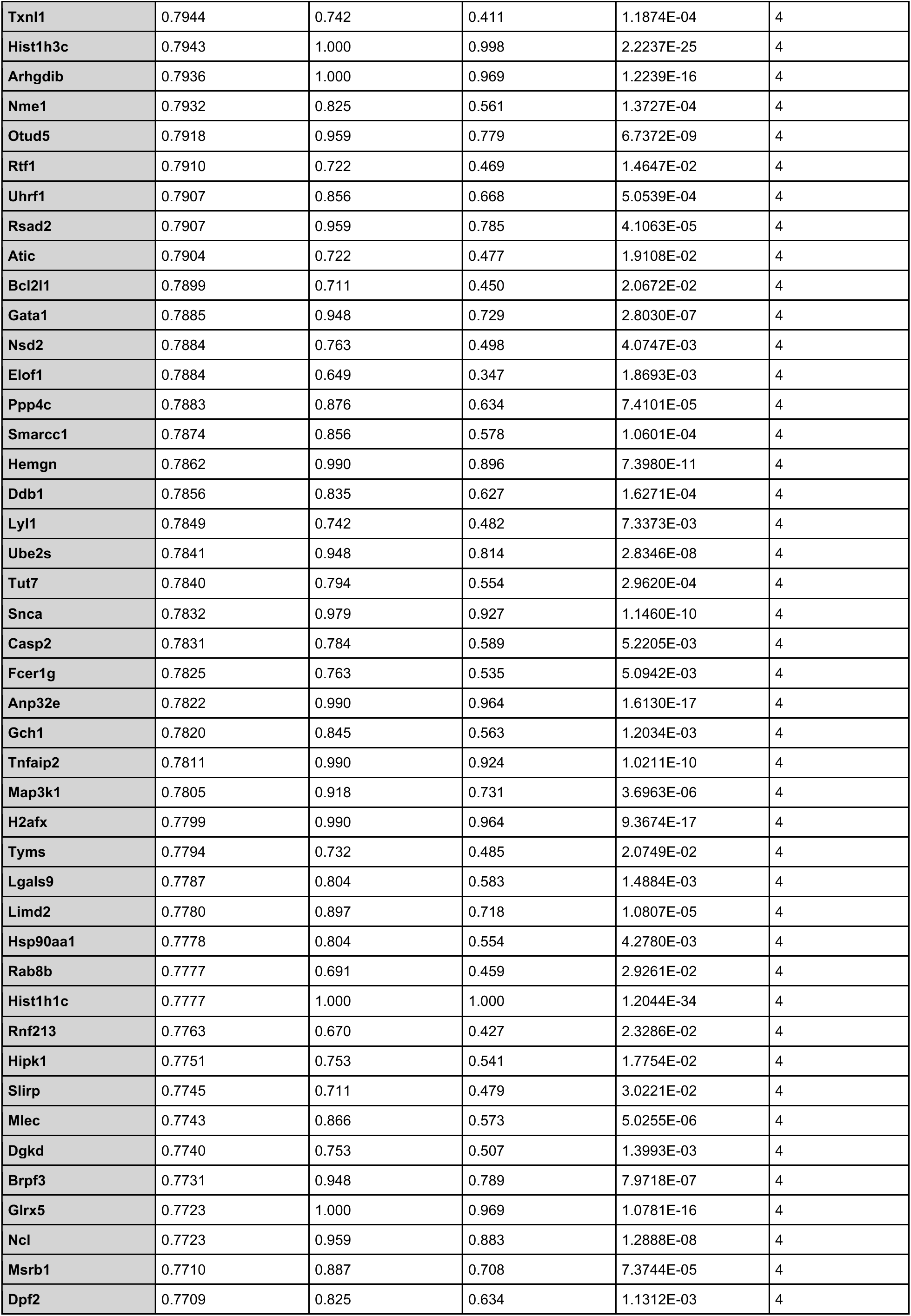
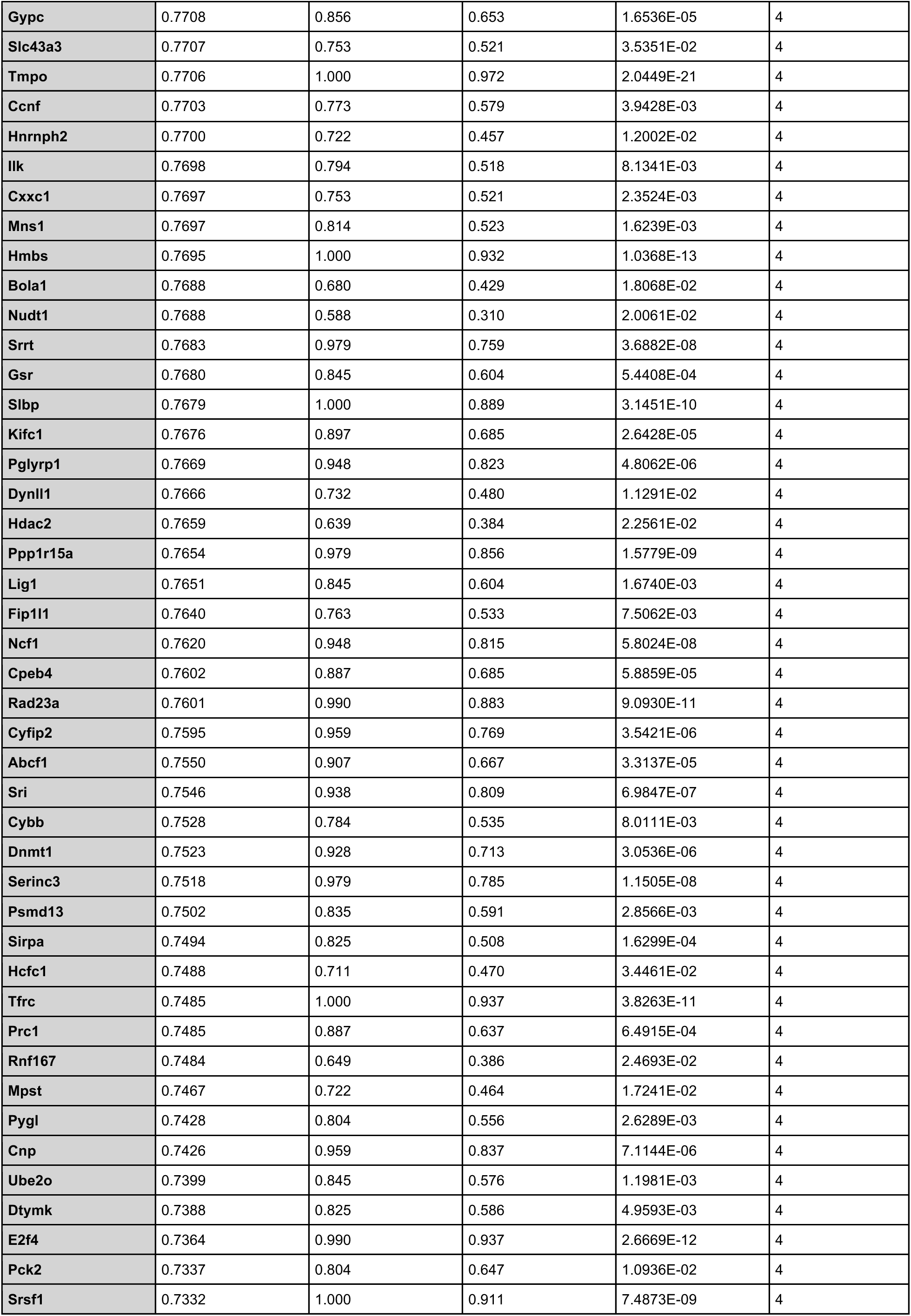
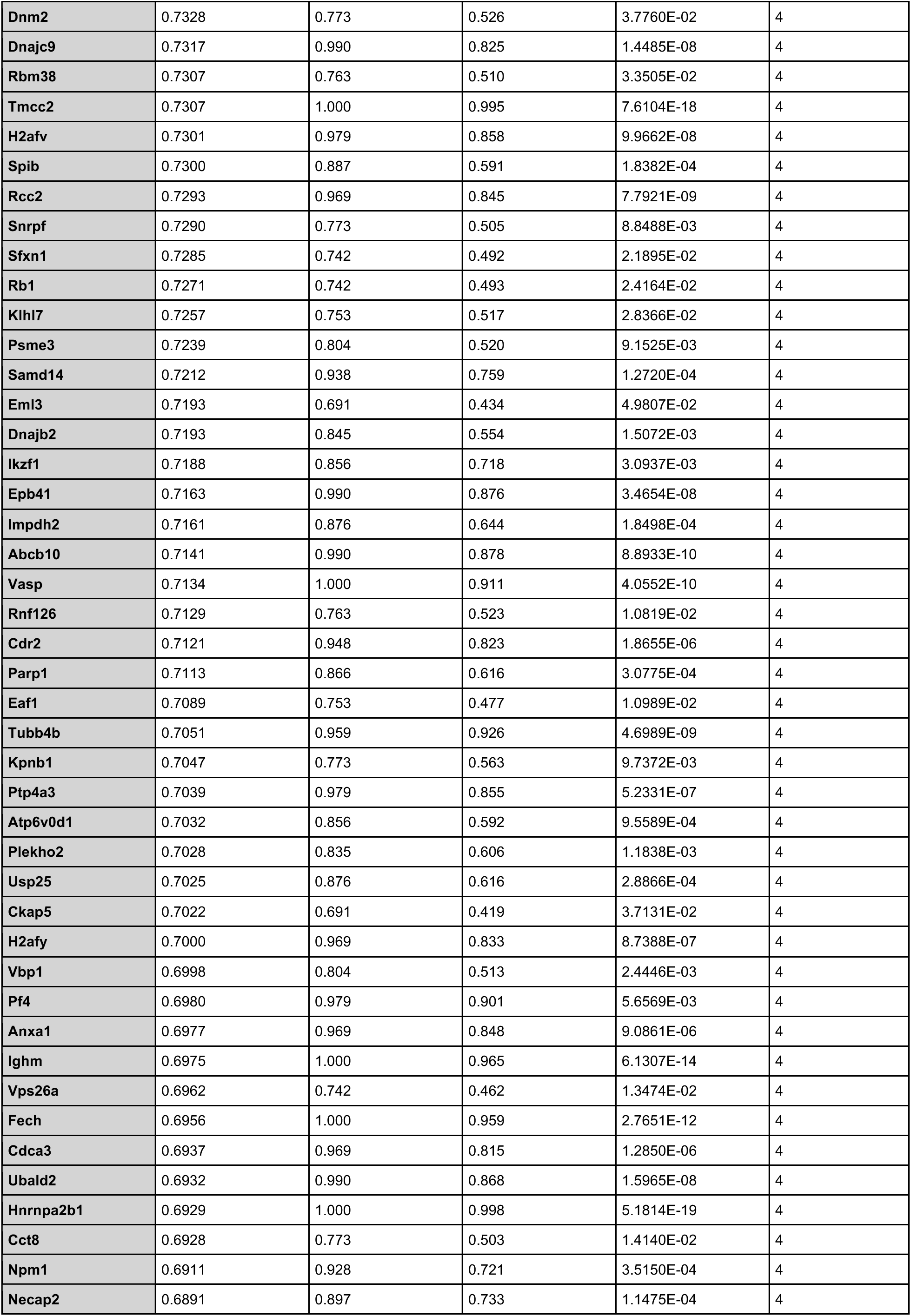
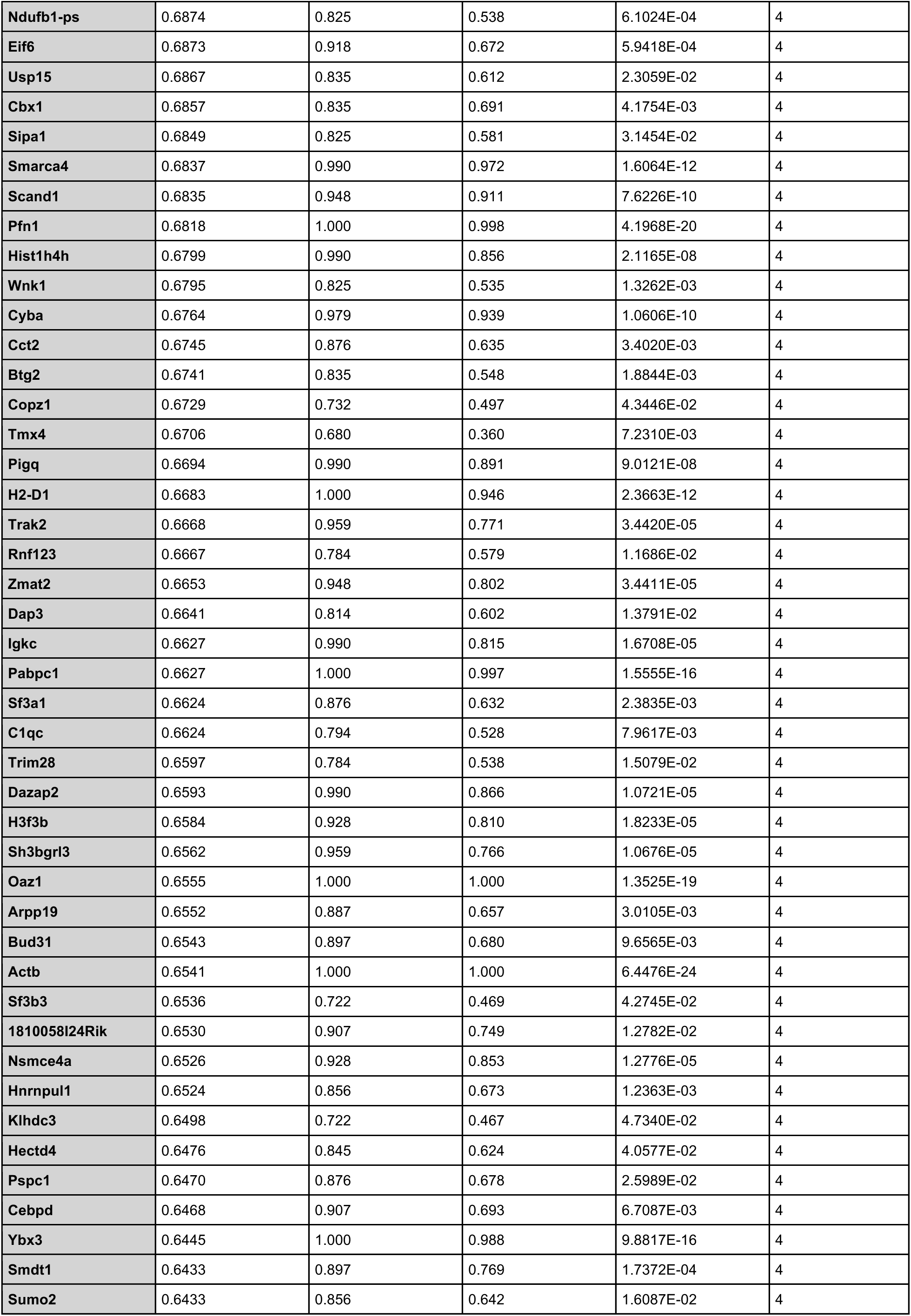
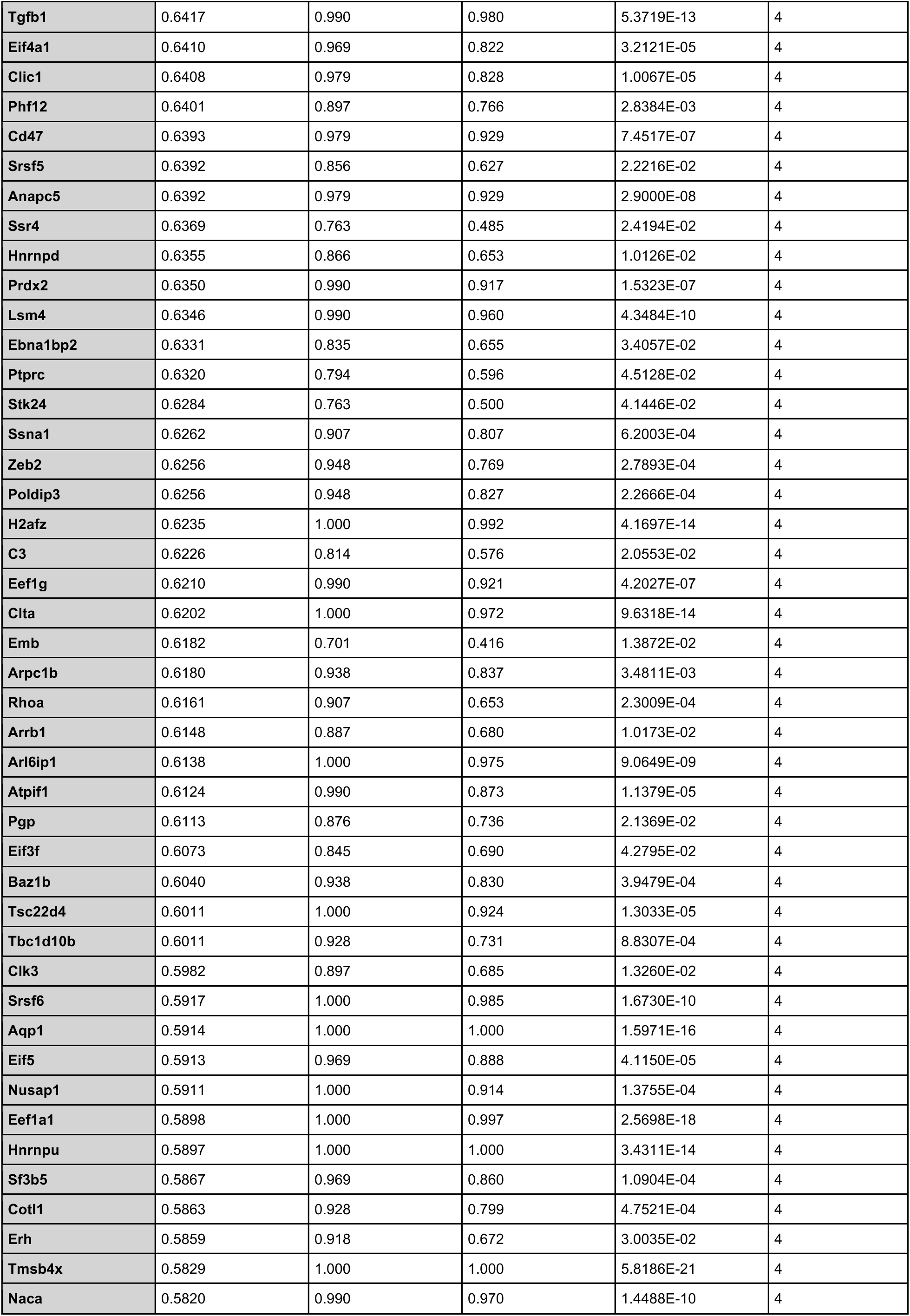
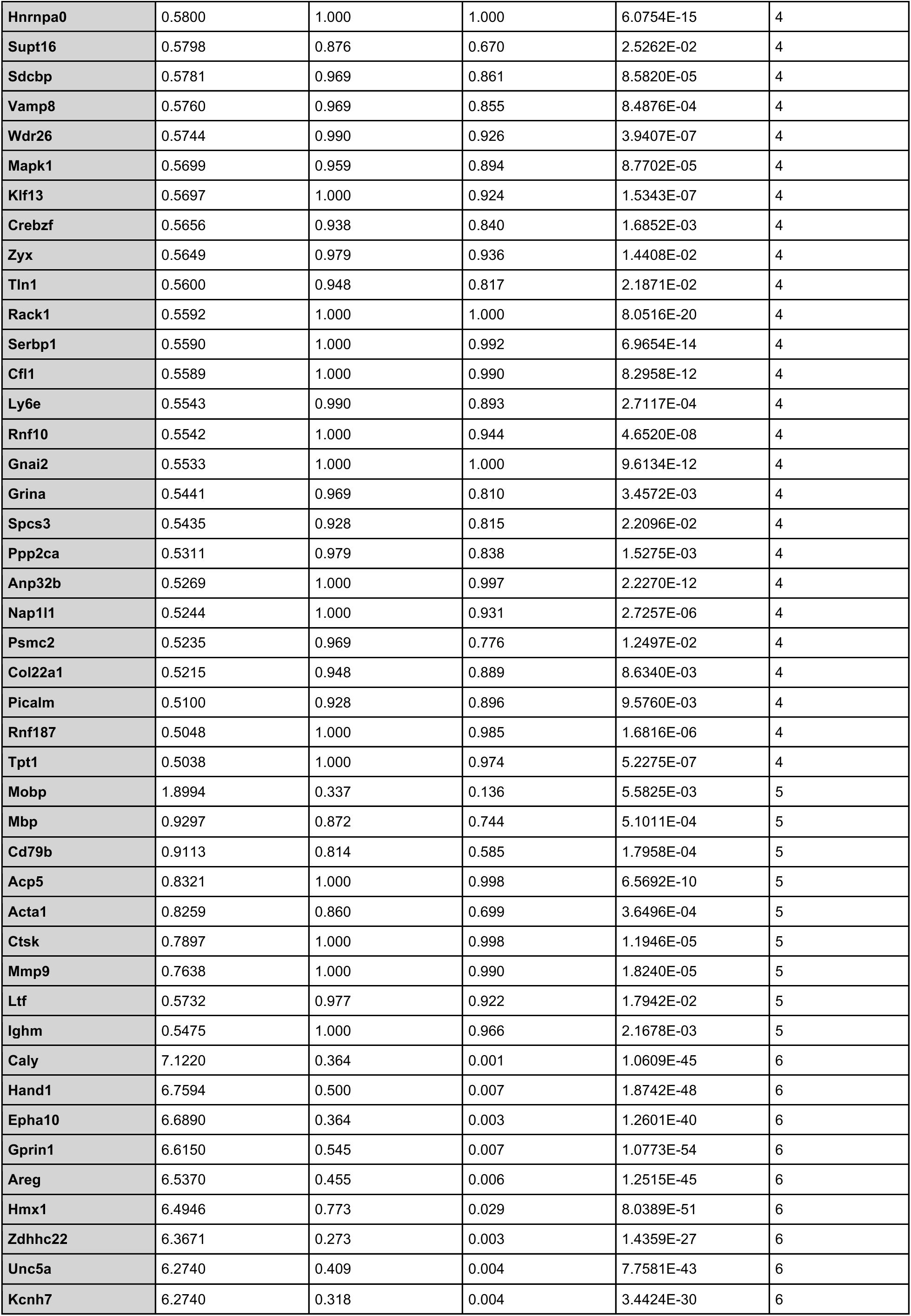
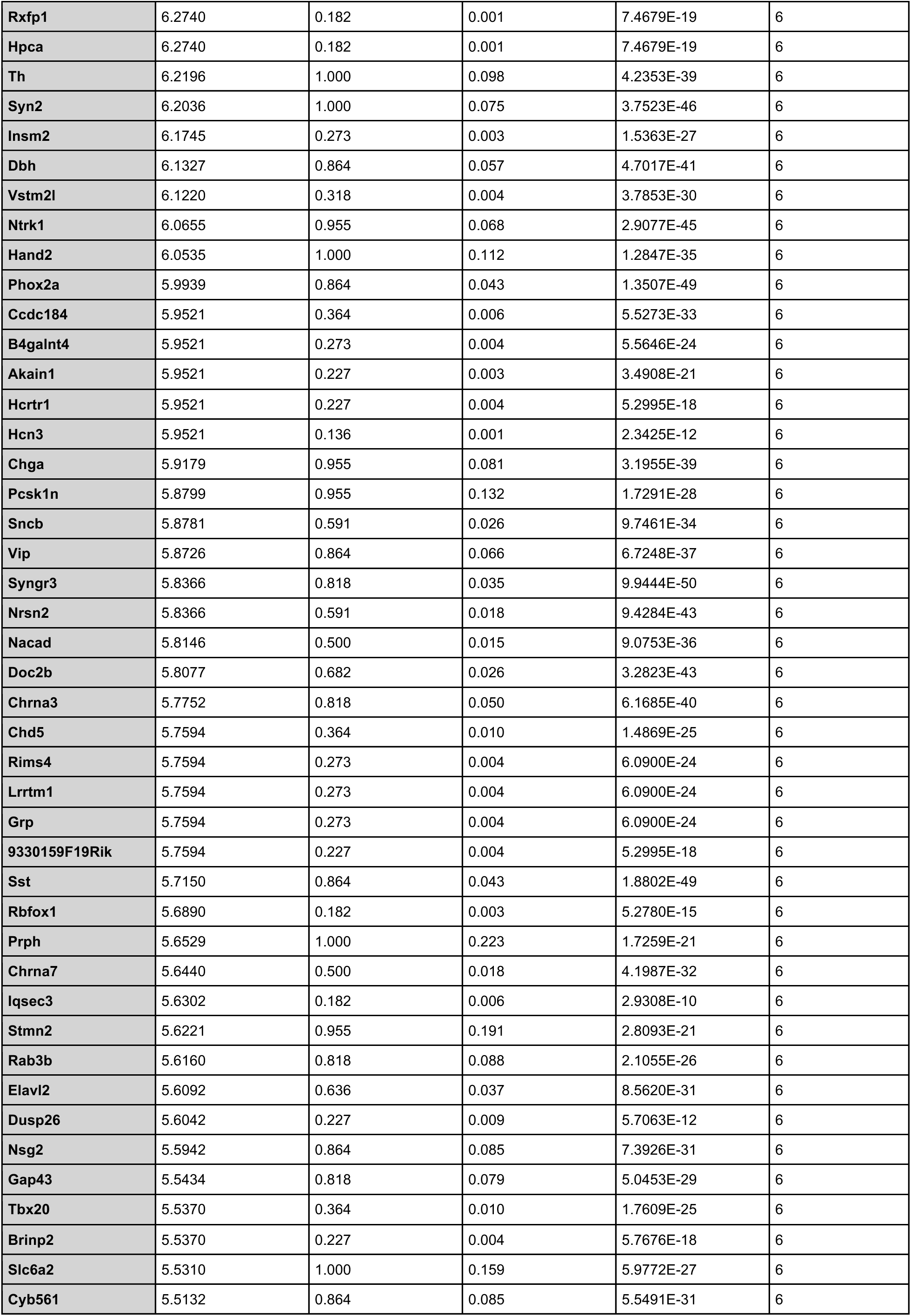
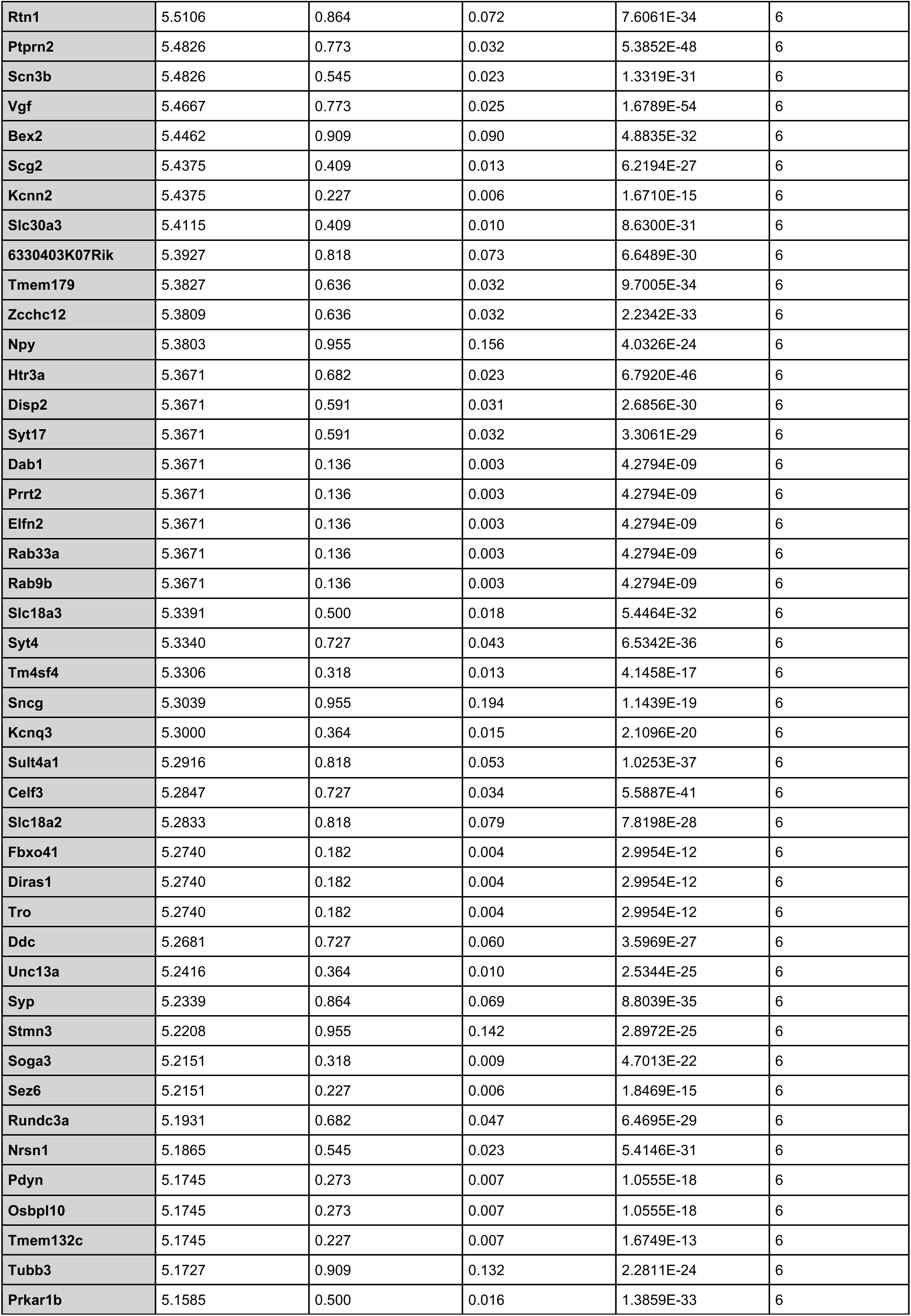
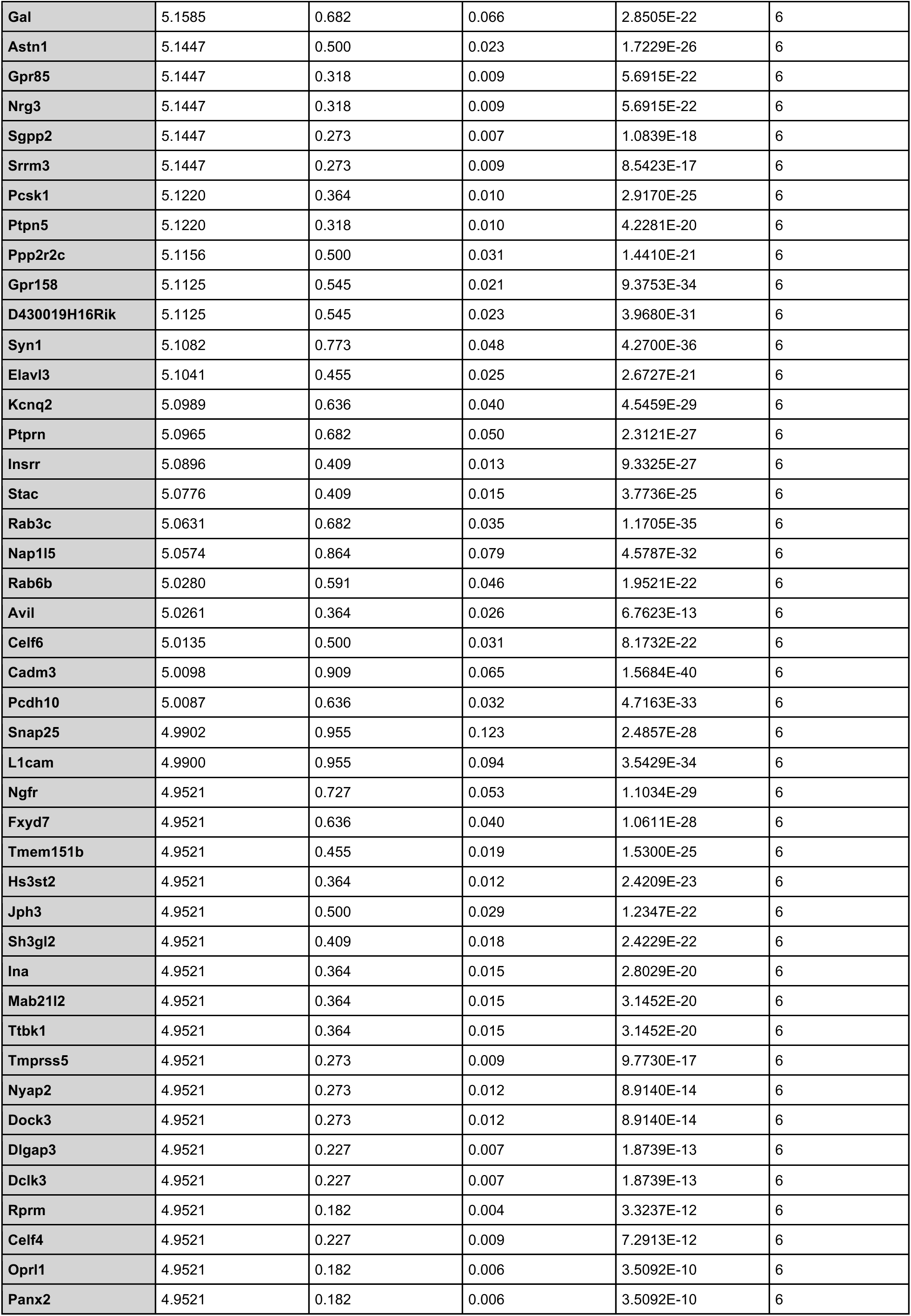
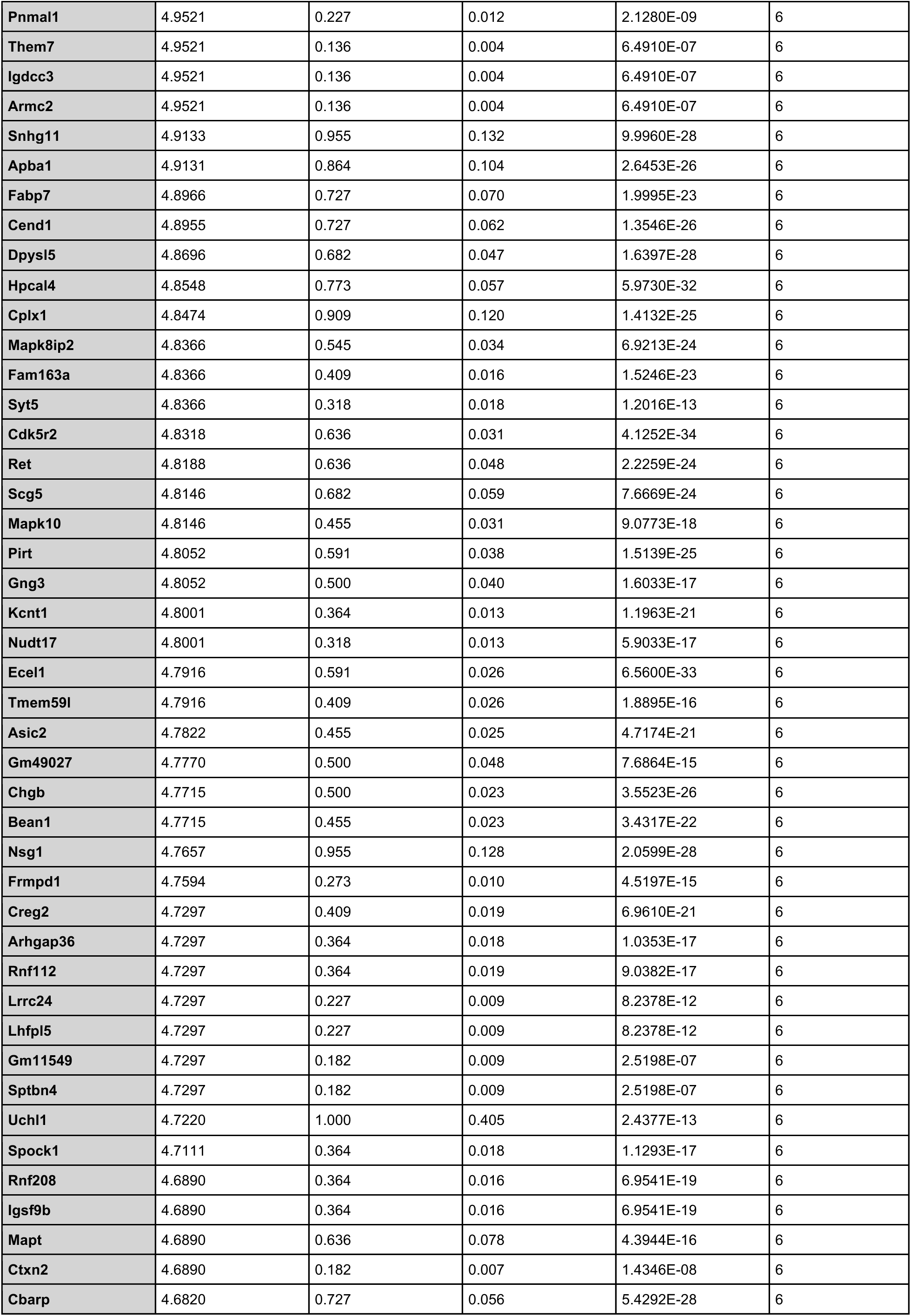
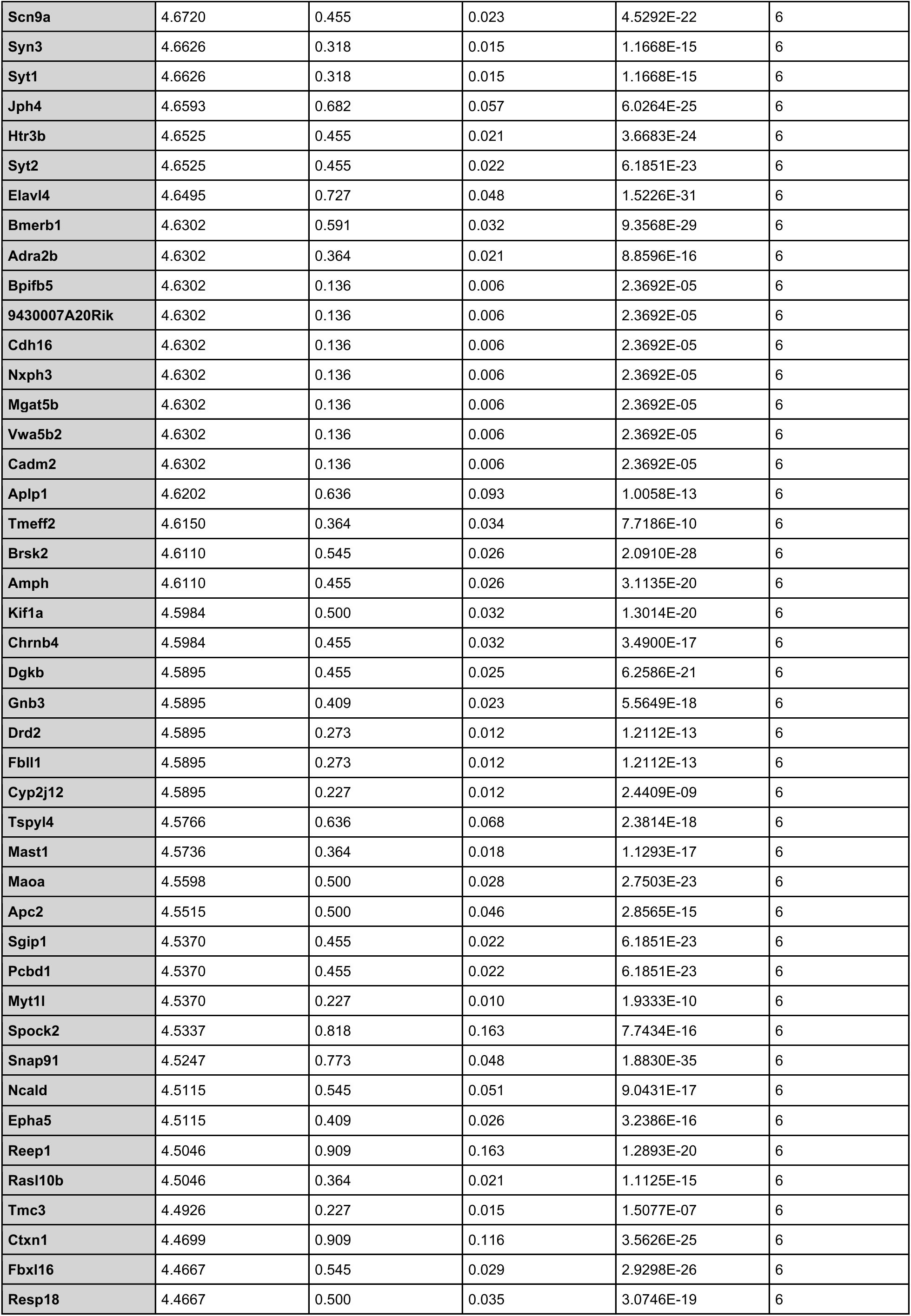
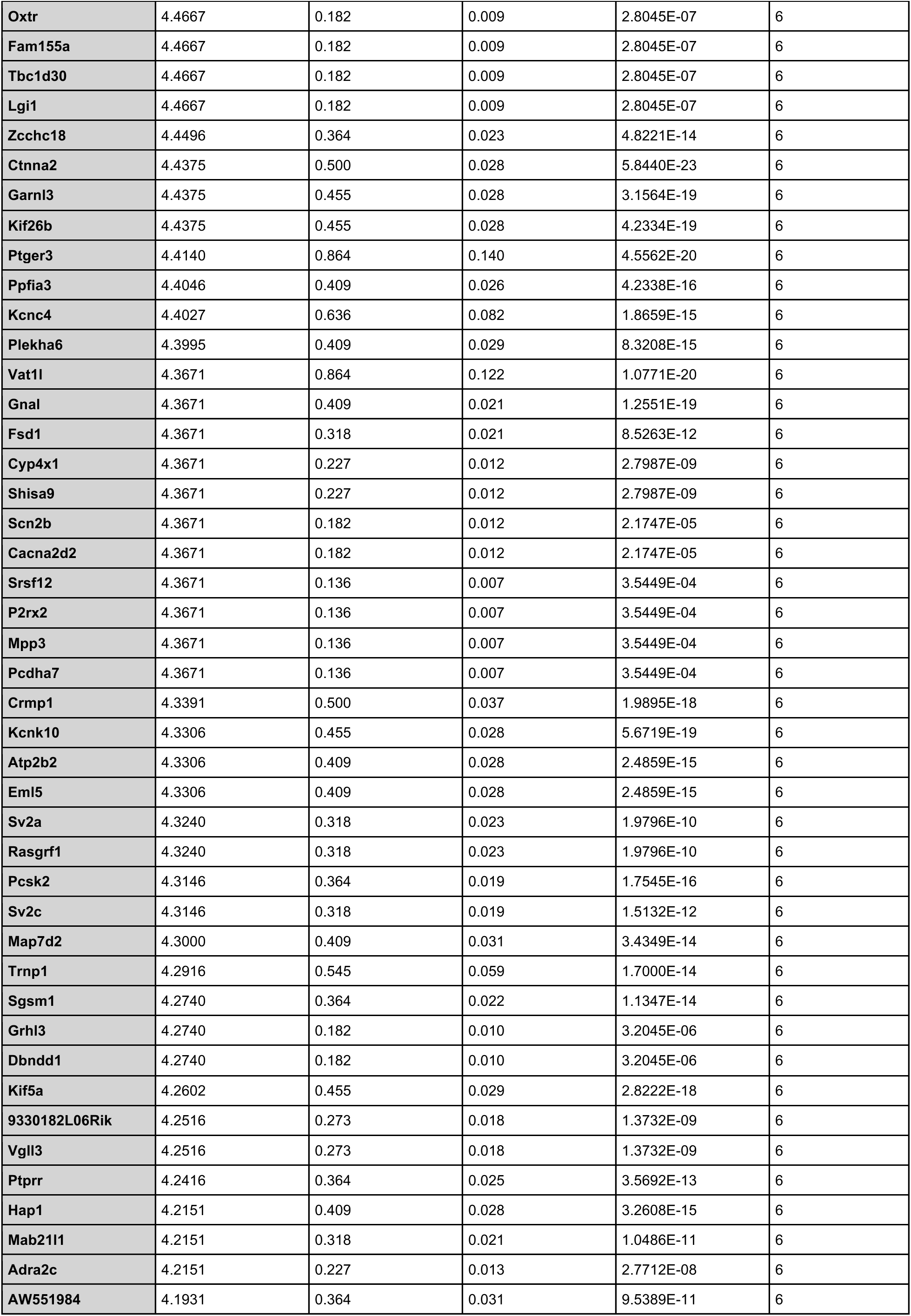
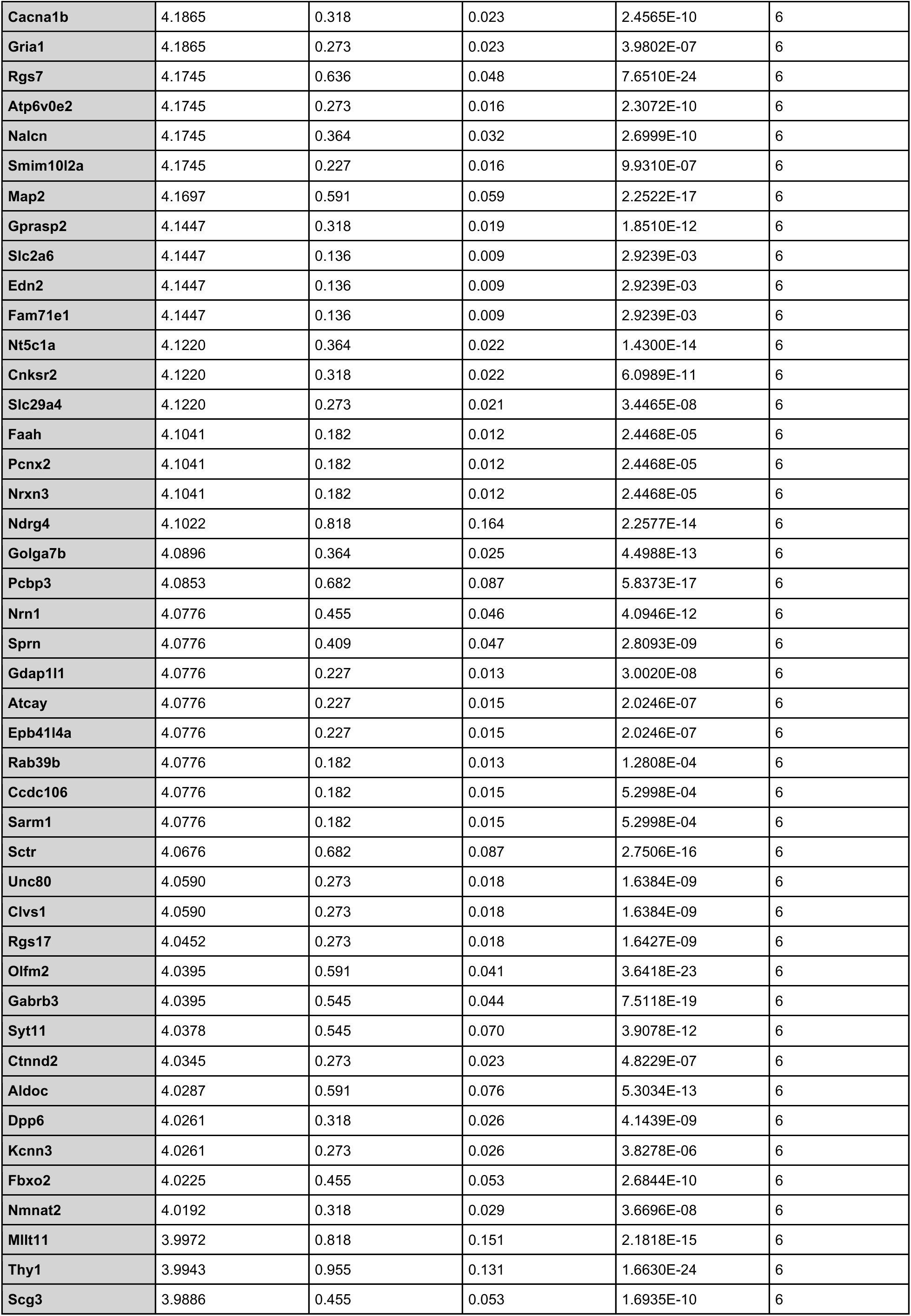
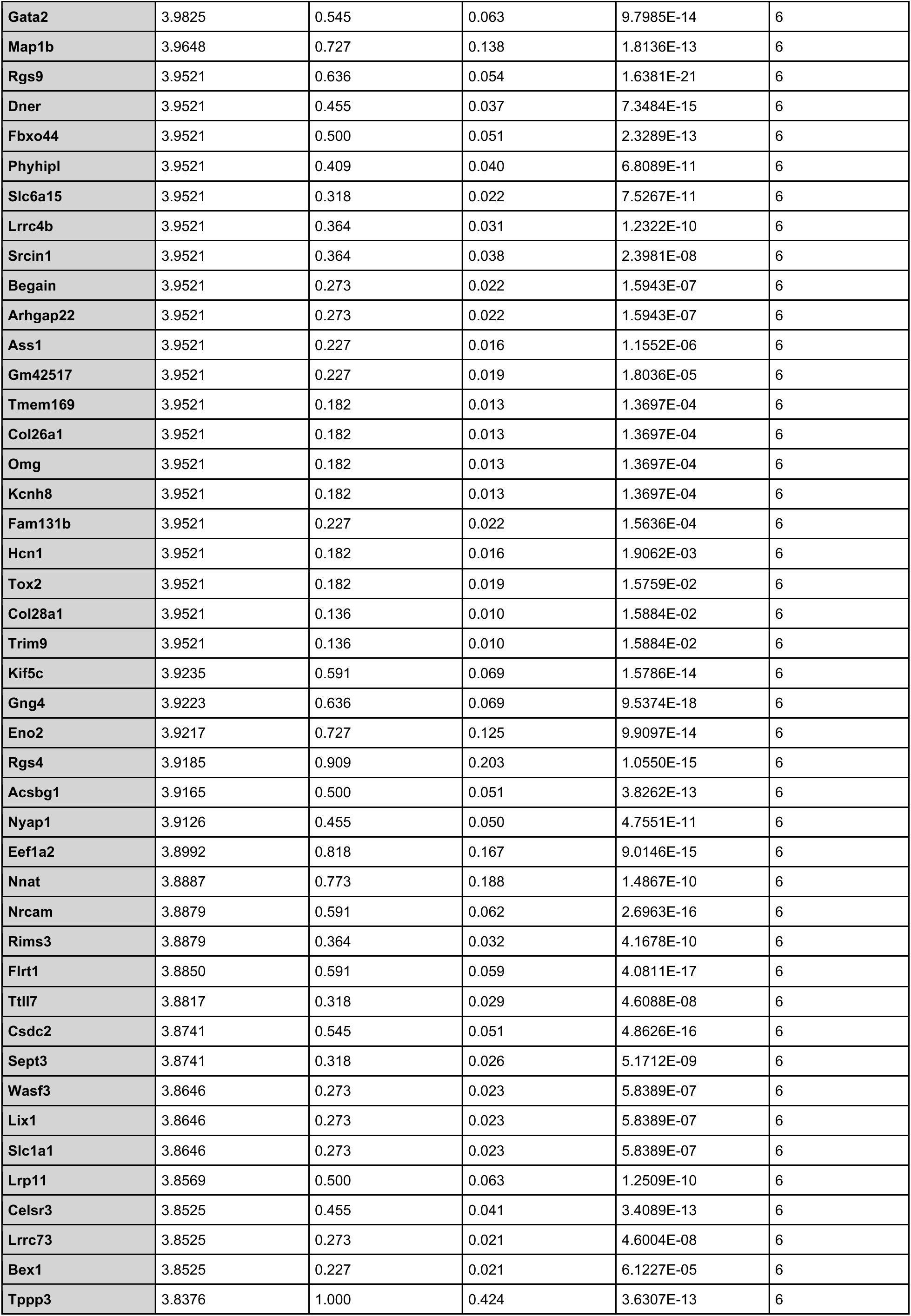
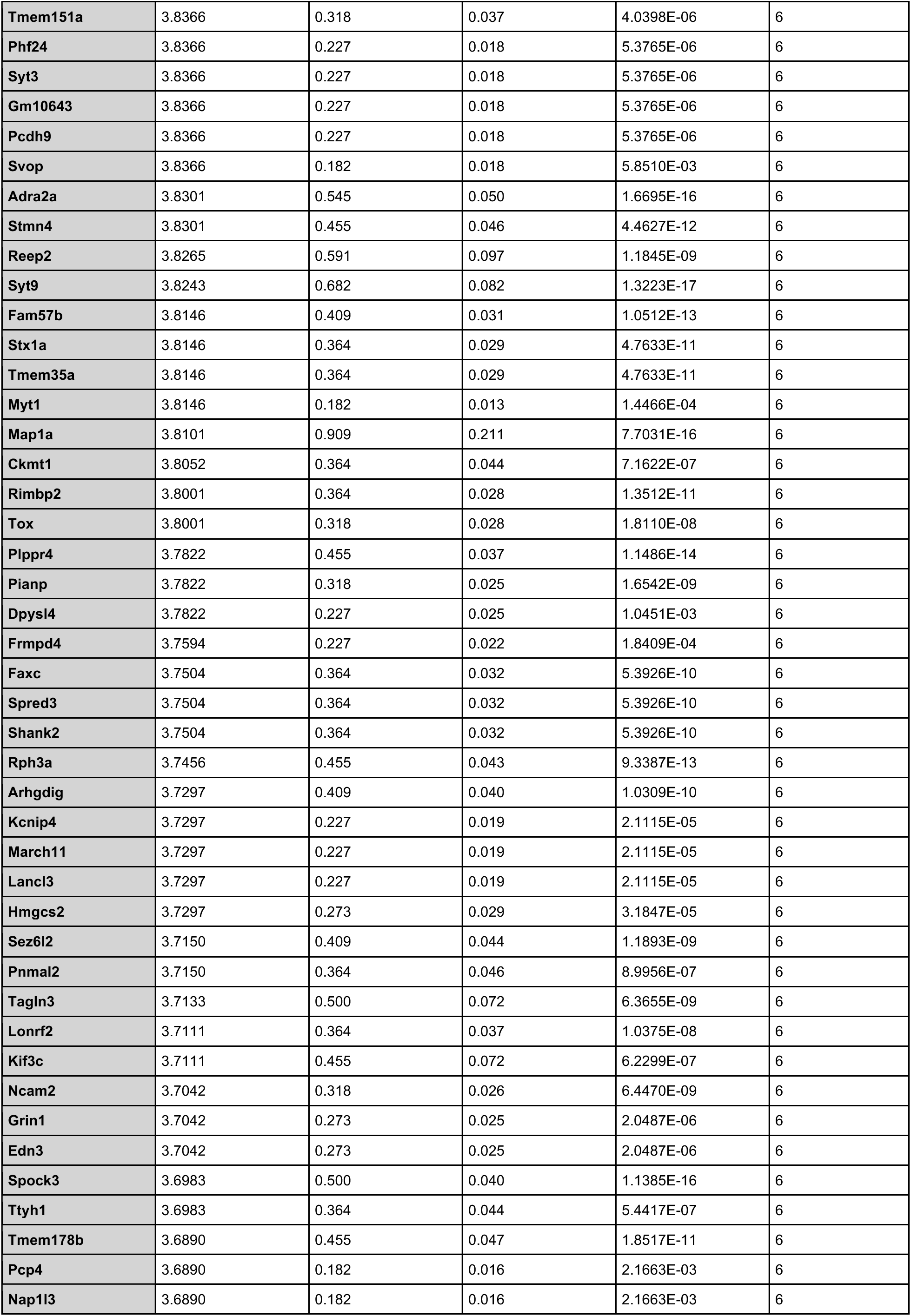
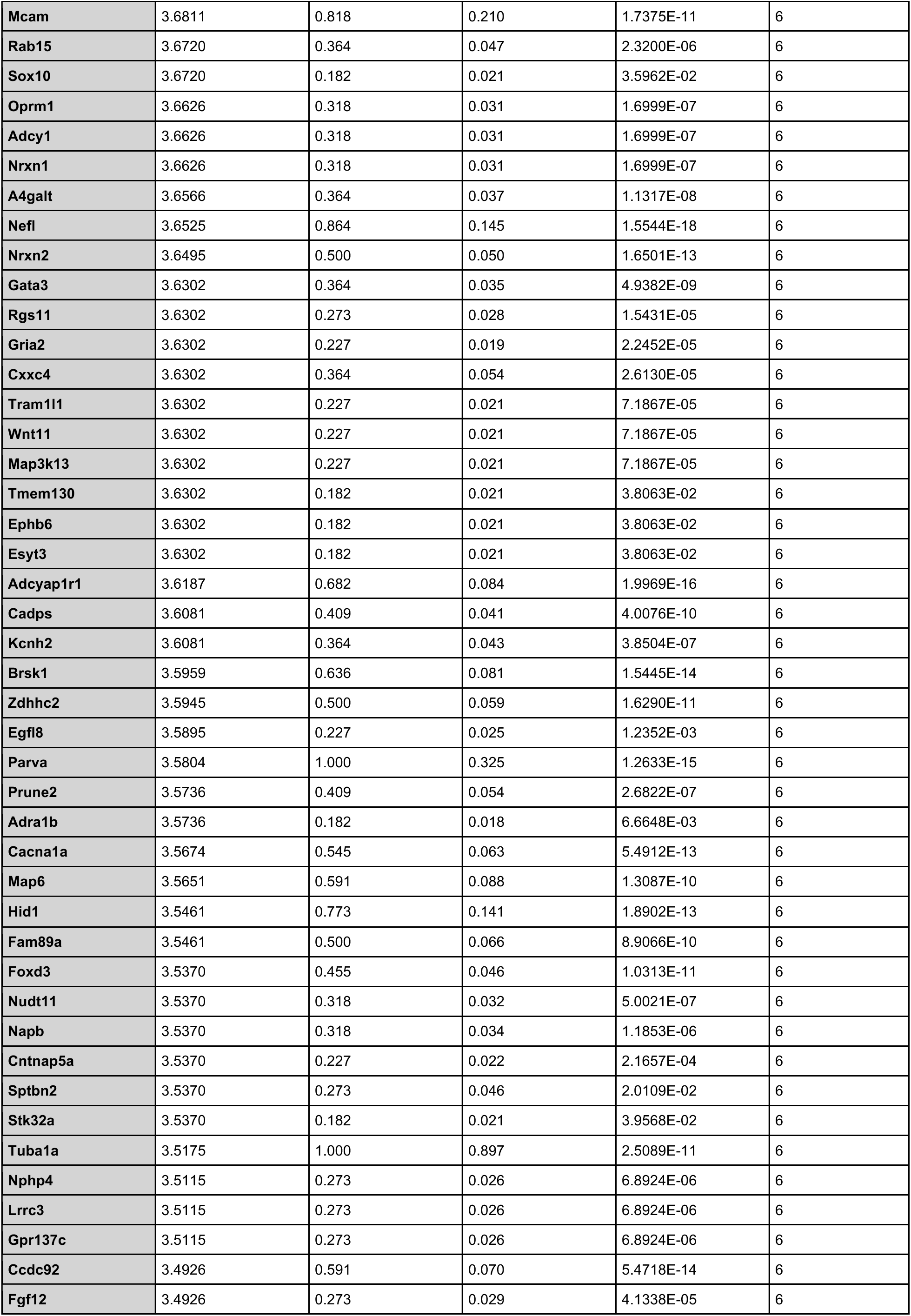
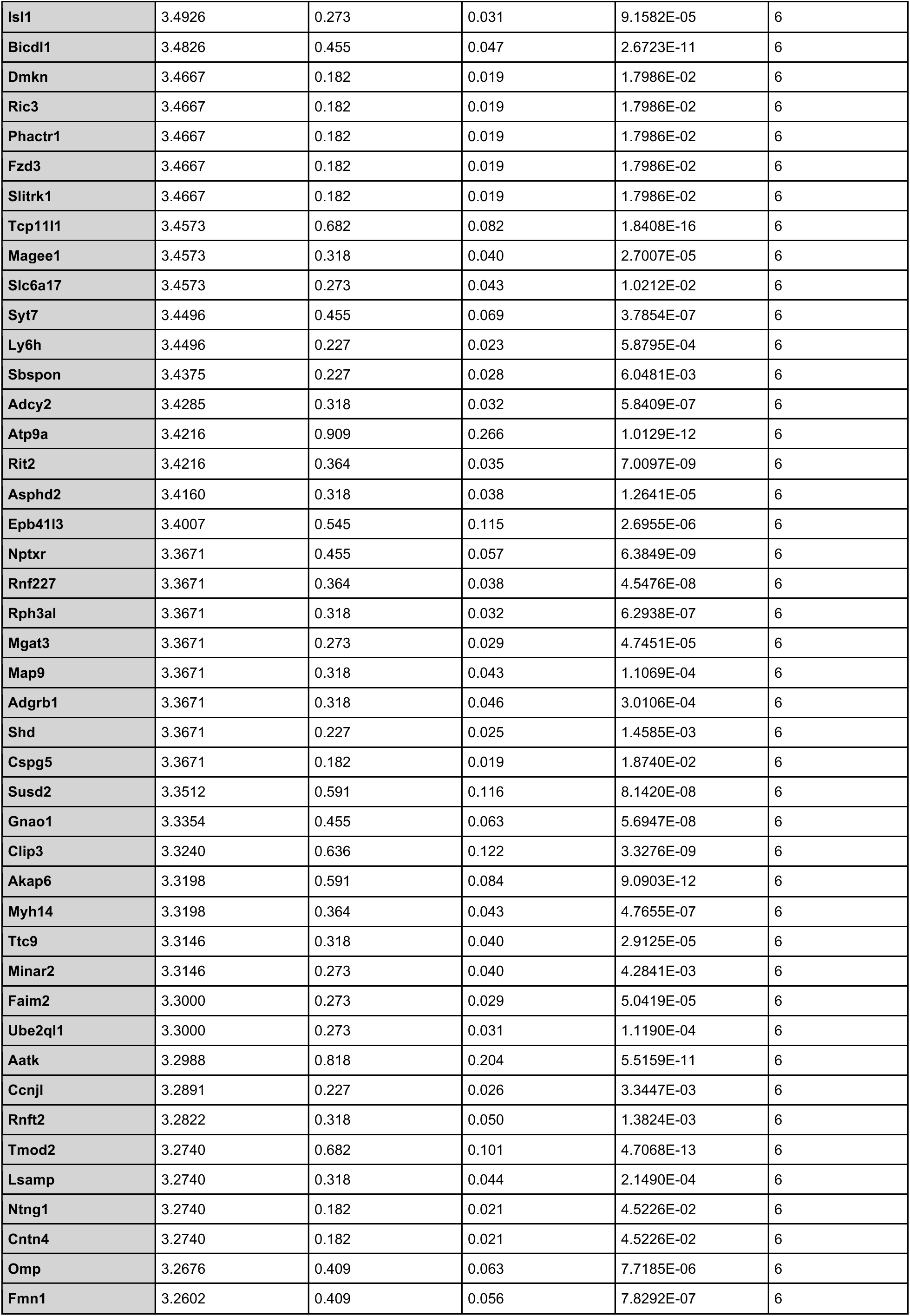
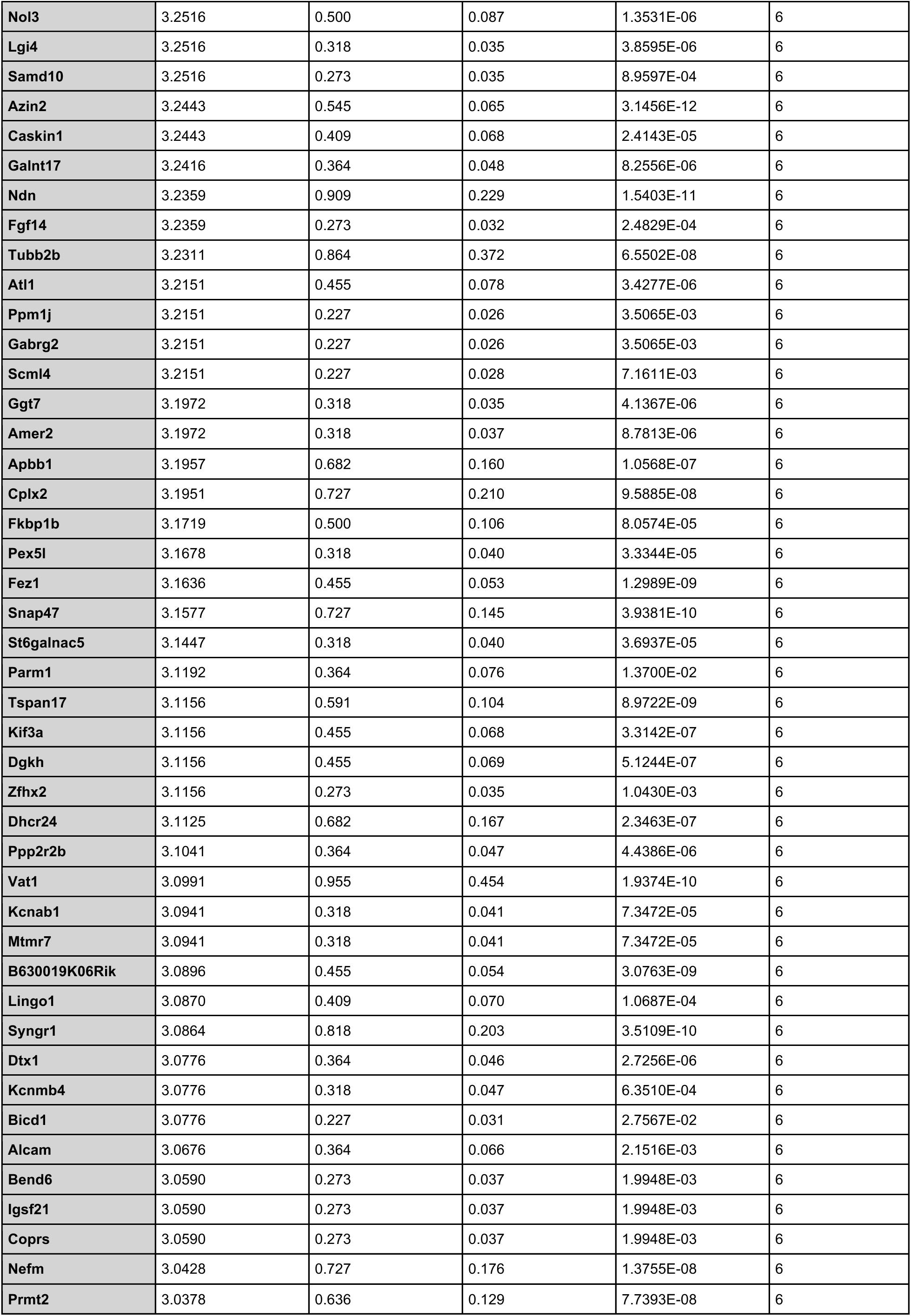
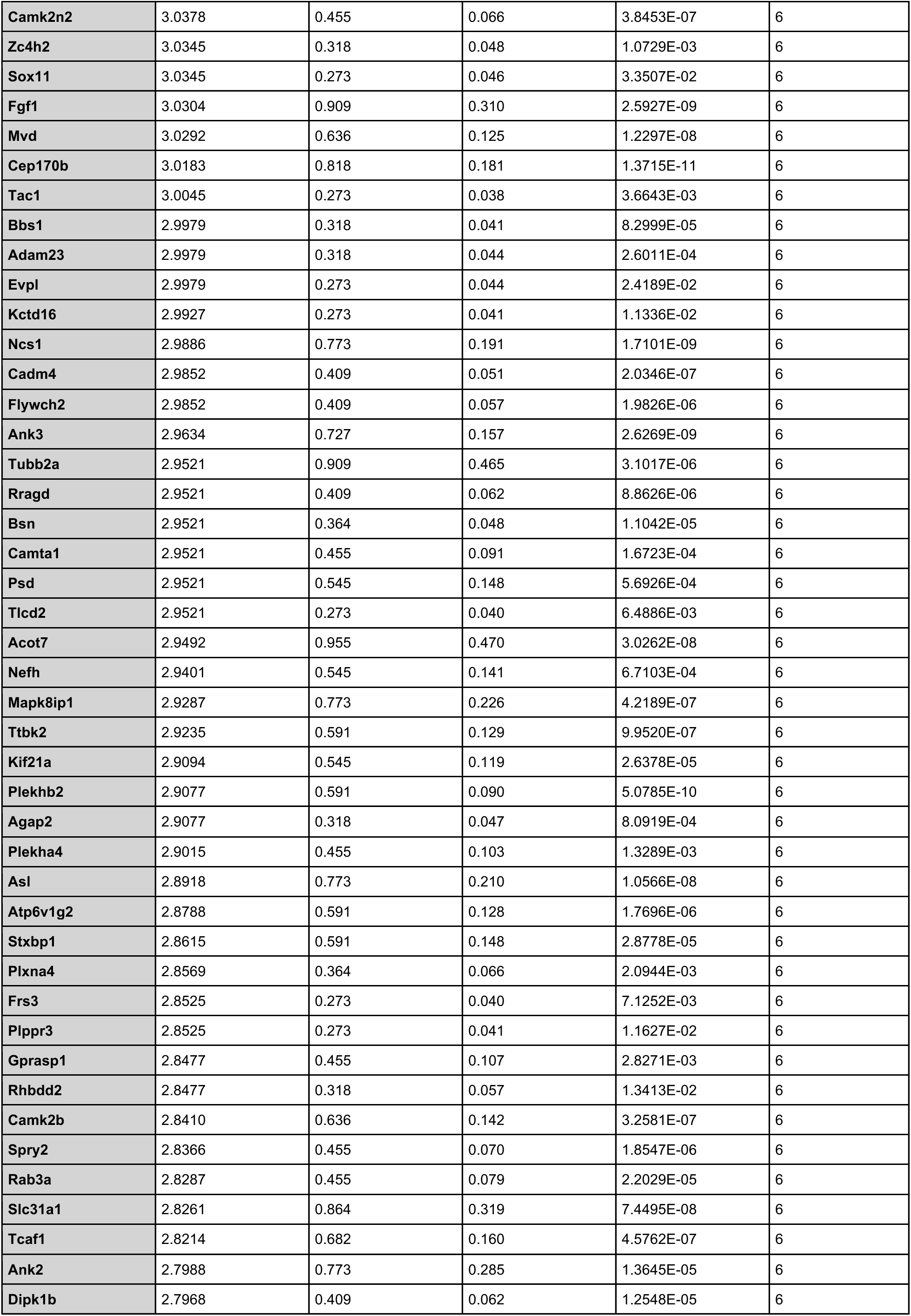
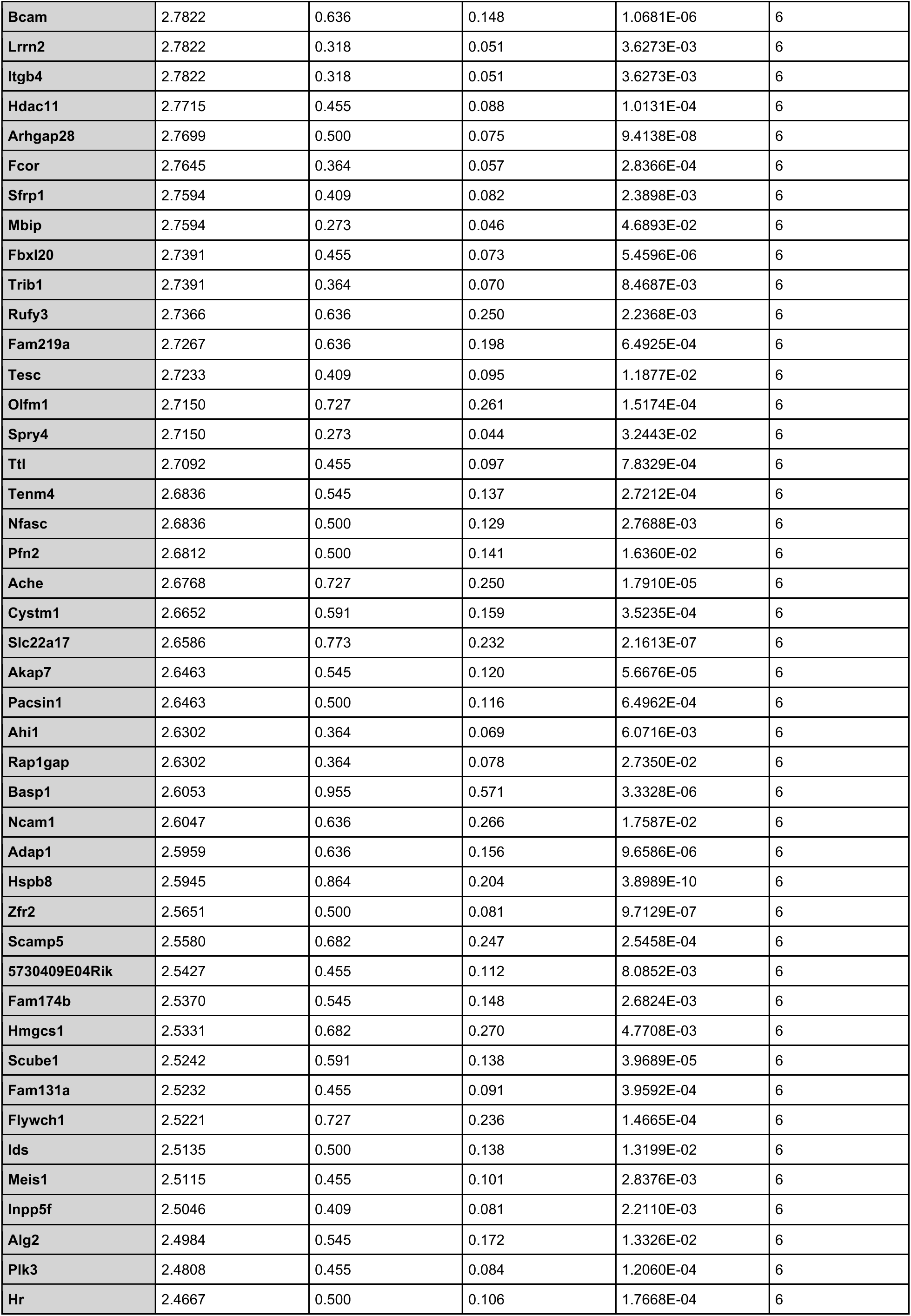
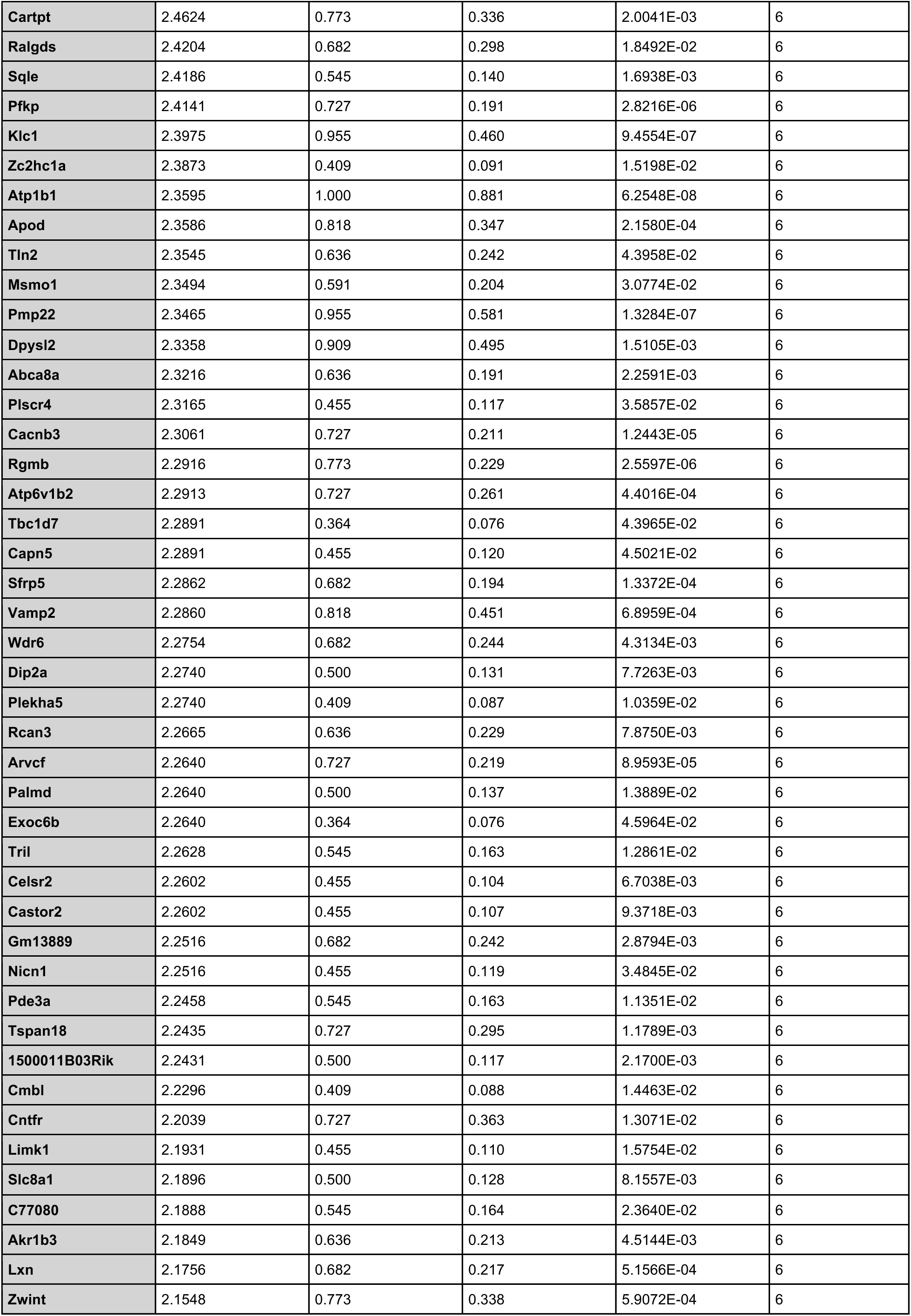
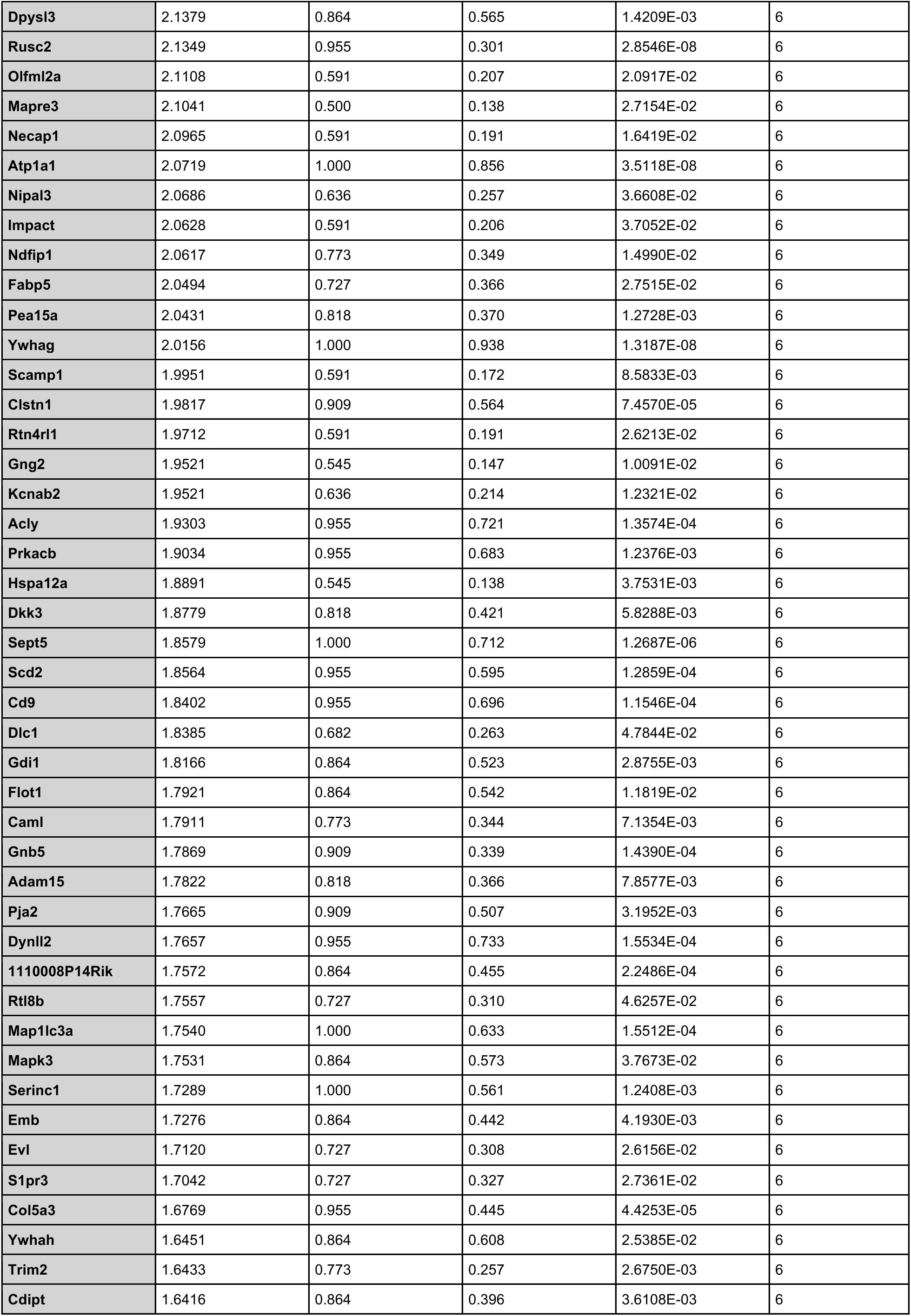
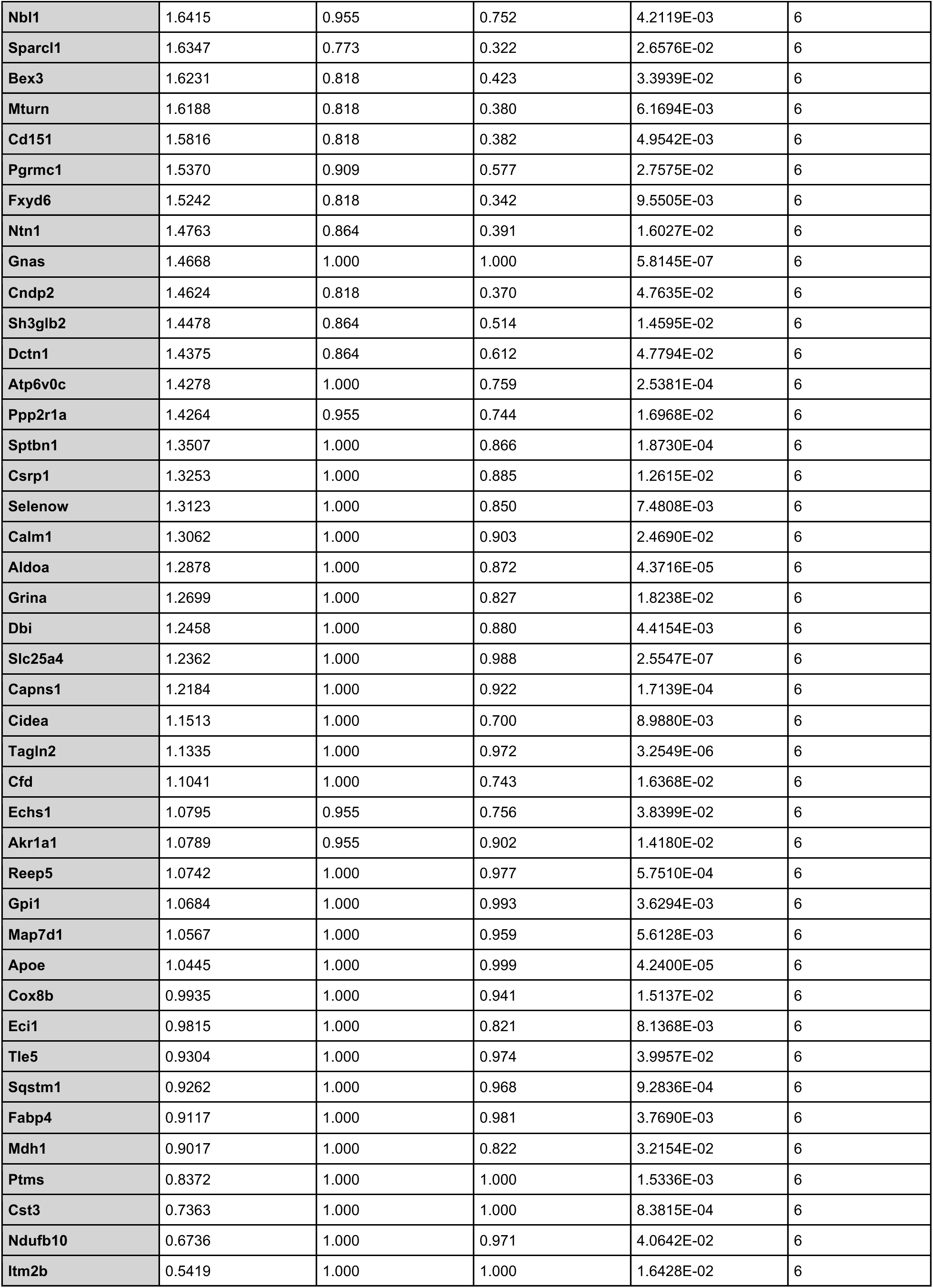
Unsupervised clustering analysis of spatial analysis over intervertebral disc regions. Markers were identified for each cluster using the SCT-normalized data. Values in the Log Fold Change column represent the natural log fold change (log-normalized) of gene expression in the target cluster relative to all other spots. The pct.1 and pct.2 columns indicate the proportion of spots expressing the gene within the target cluster and across all other clusters, respectively. Marker genes shown were restricted to those with a minimum detection frequency of 25% within the cluster to ensure biological representativeness. Statistical significance was defined as a Bonferroni-adjusted *p*-value < 0.05.

**Table 2.**
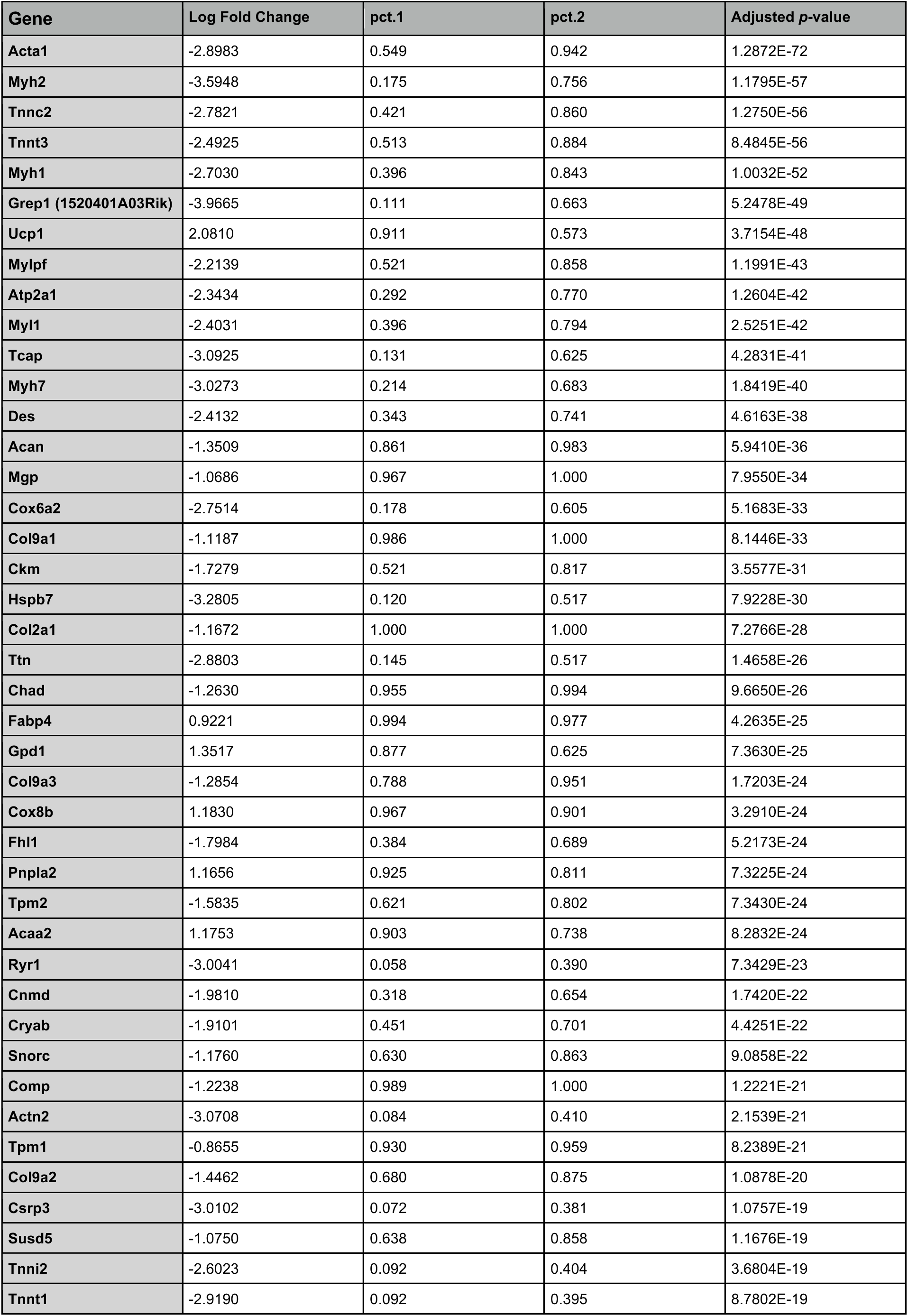
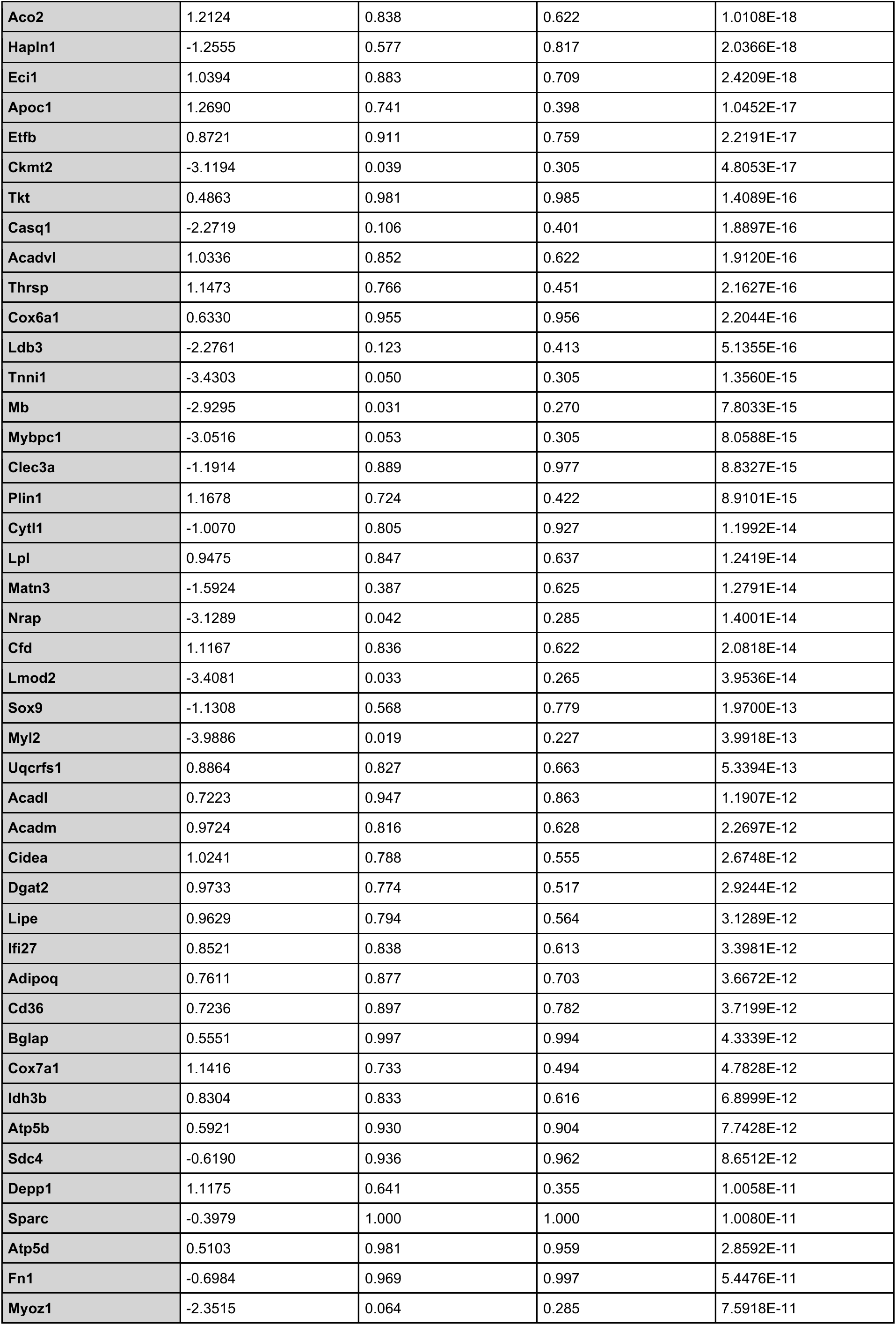
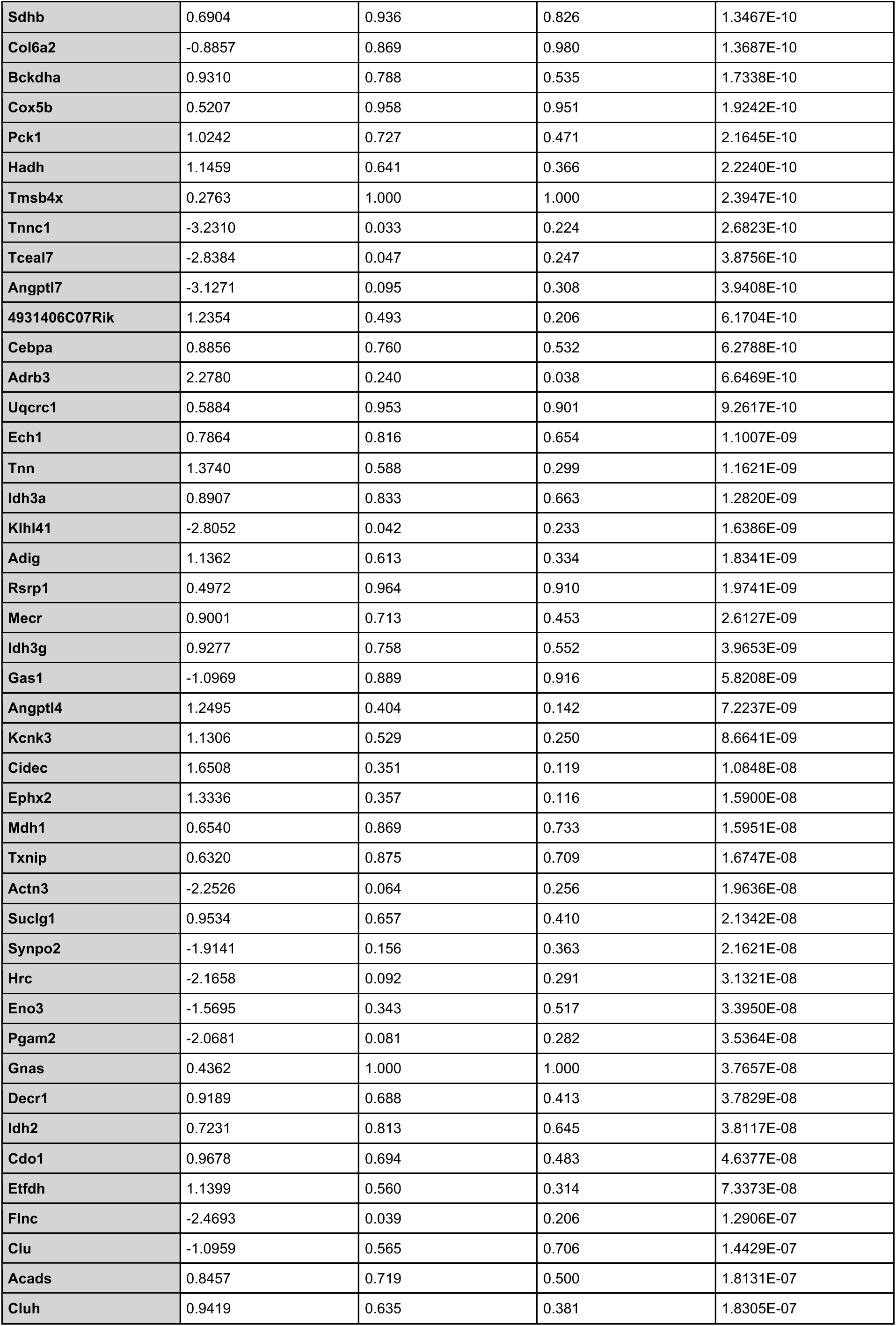
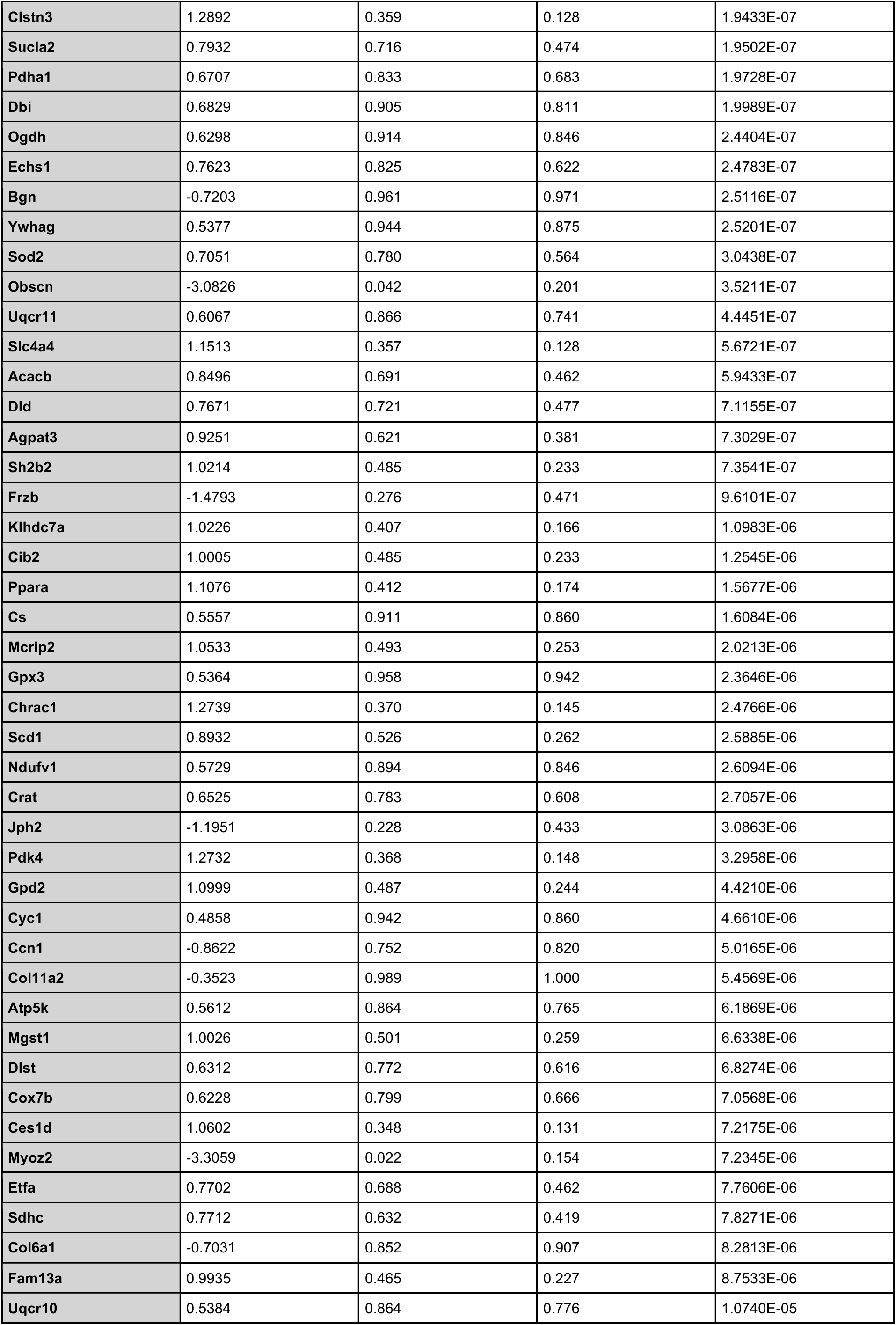
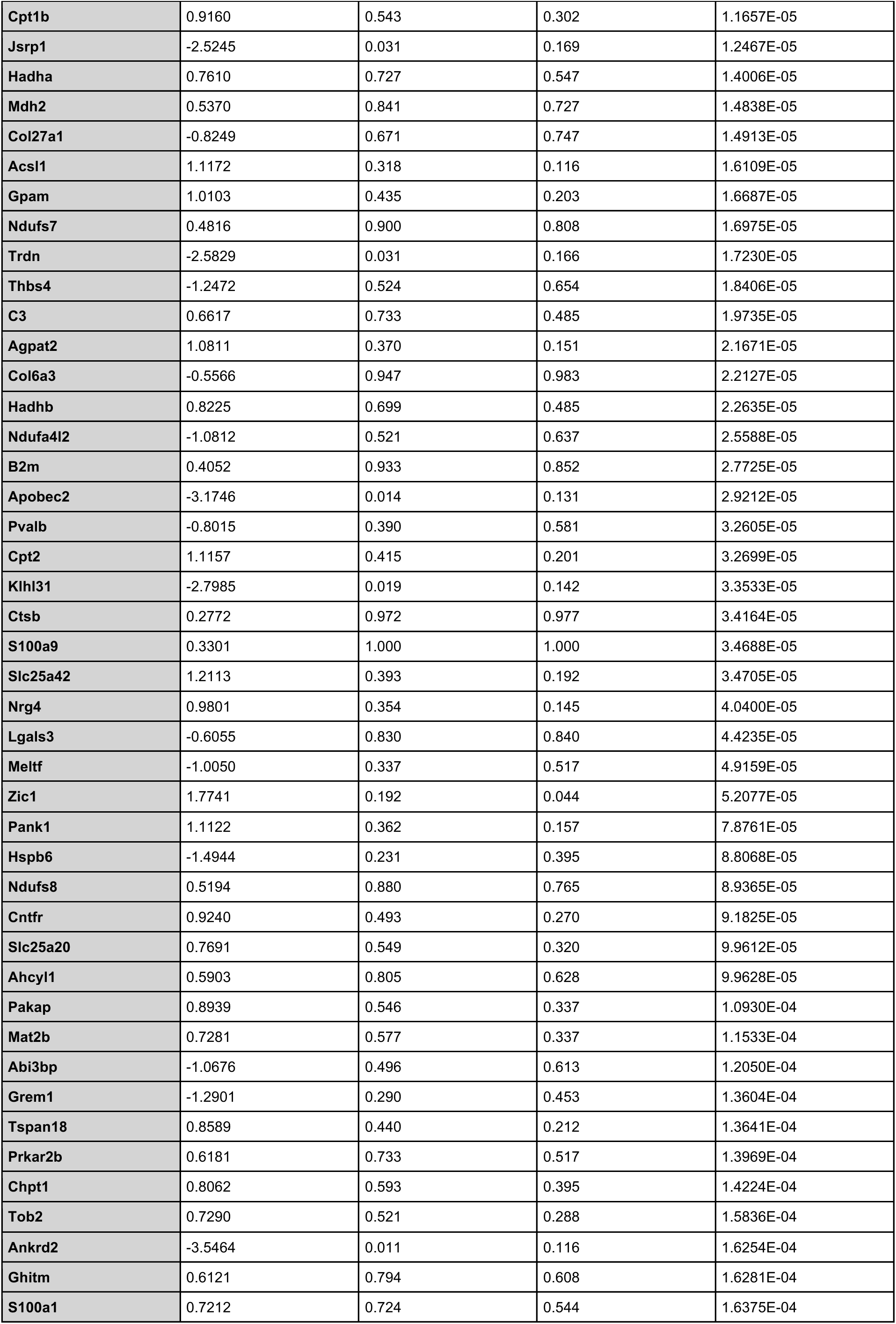
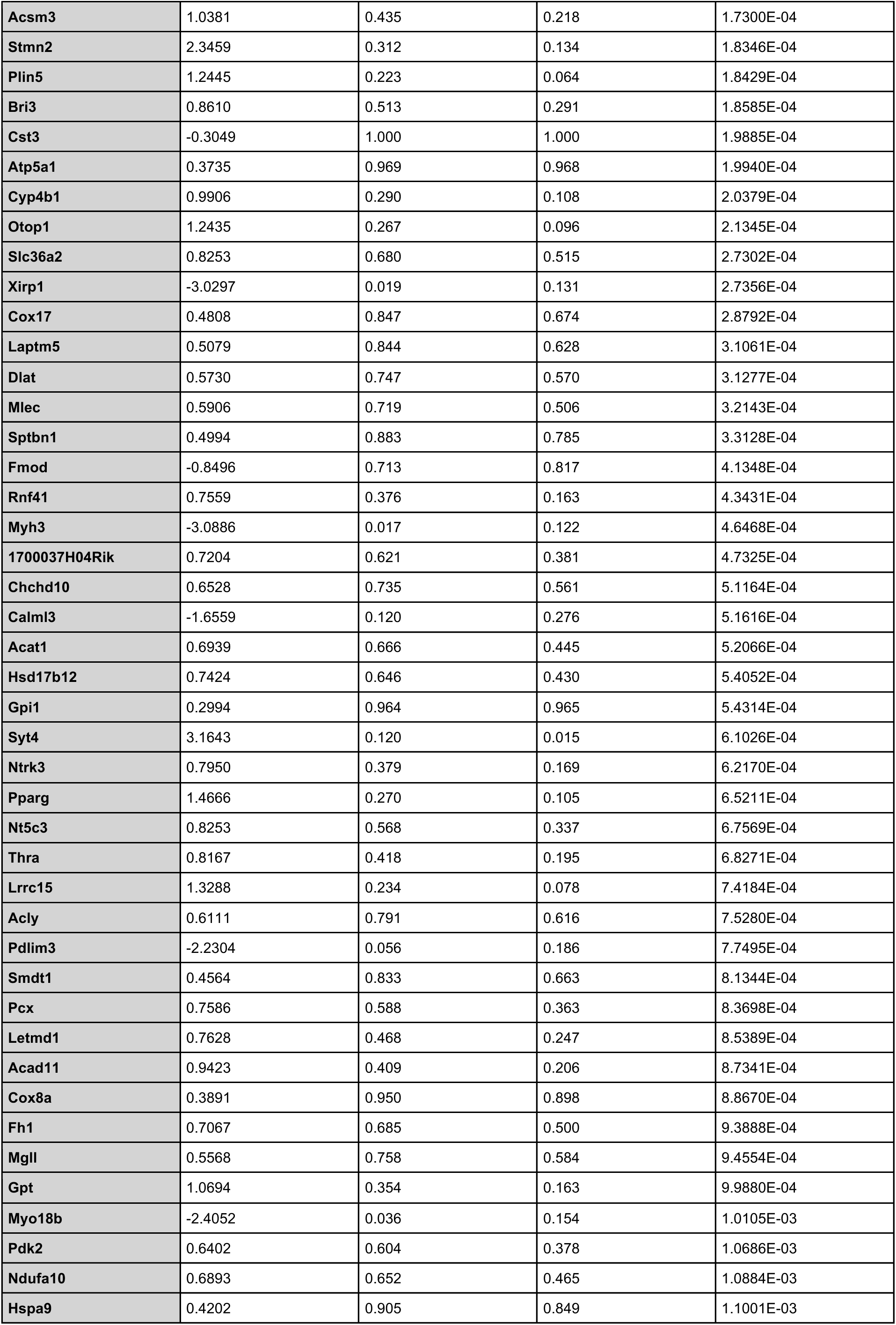
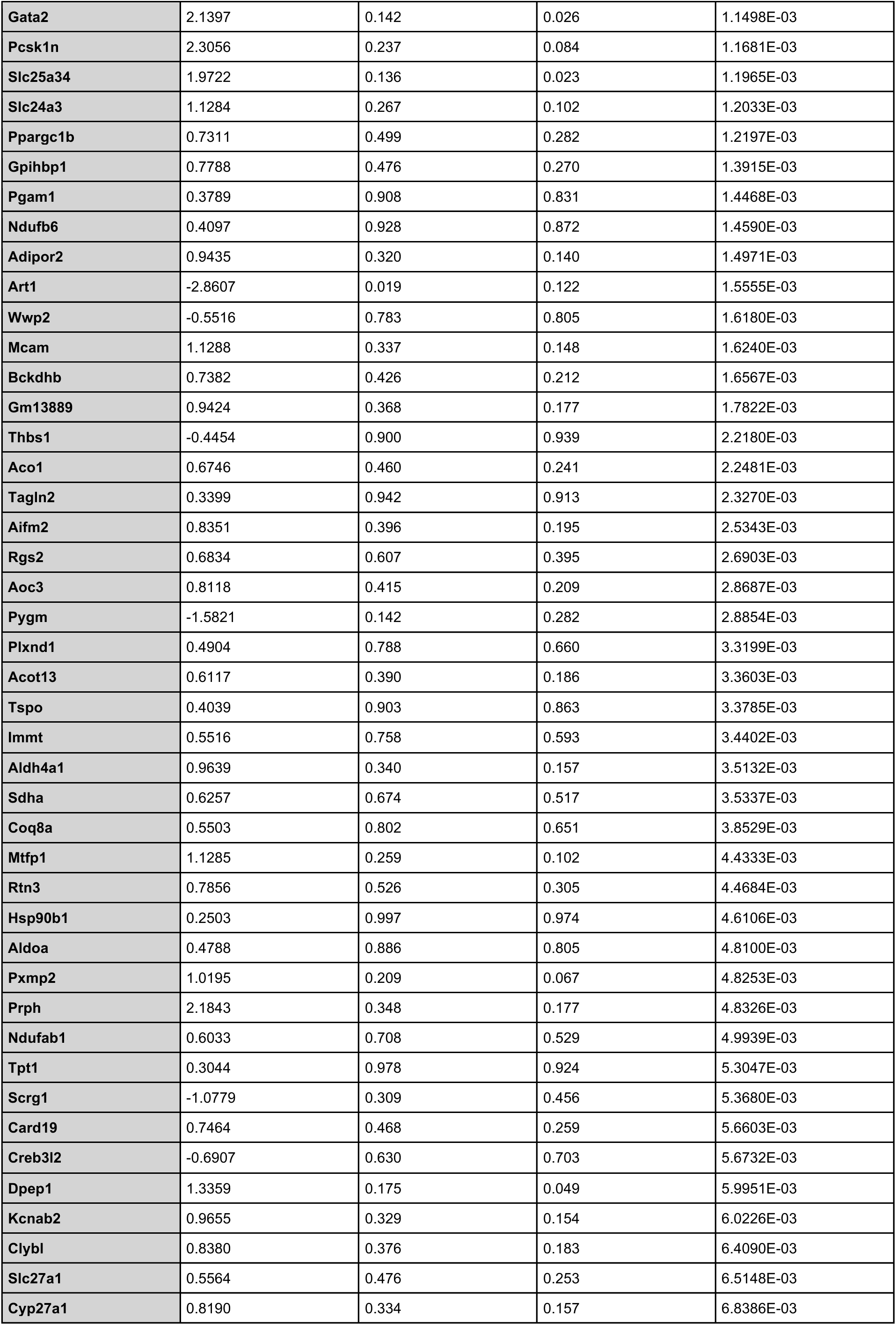
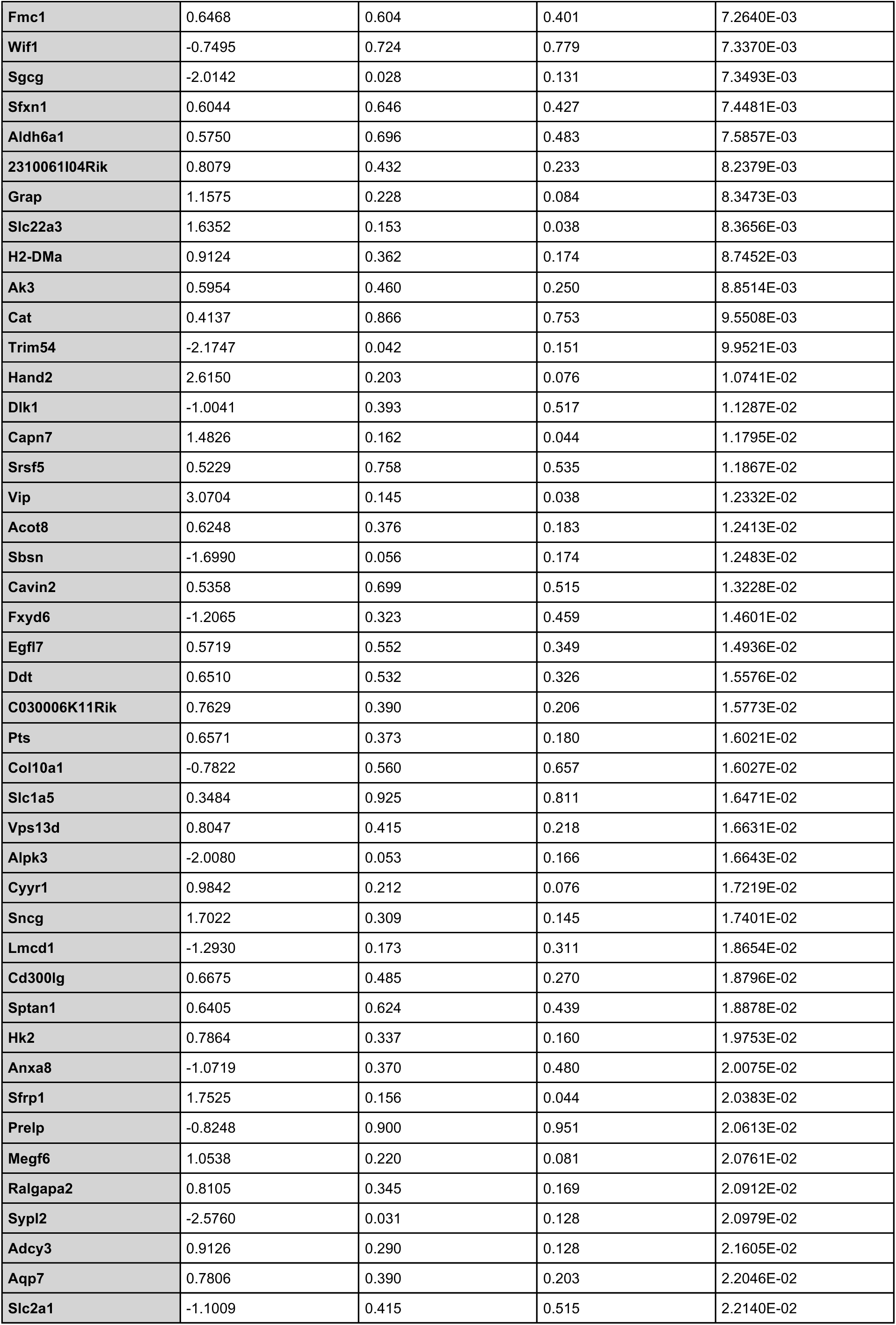
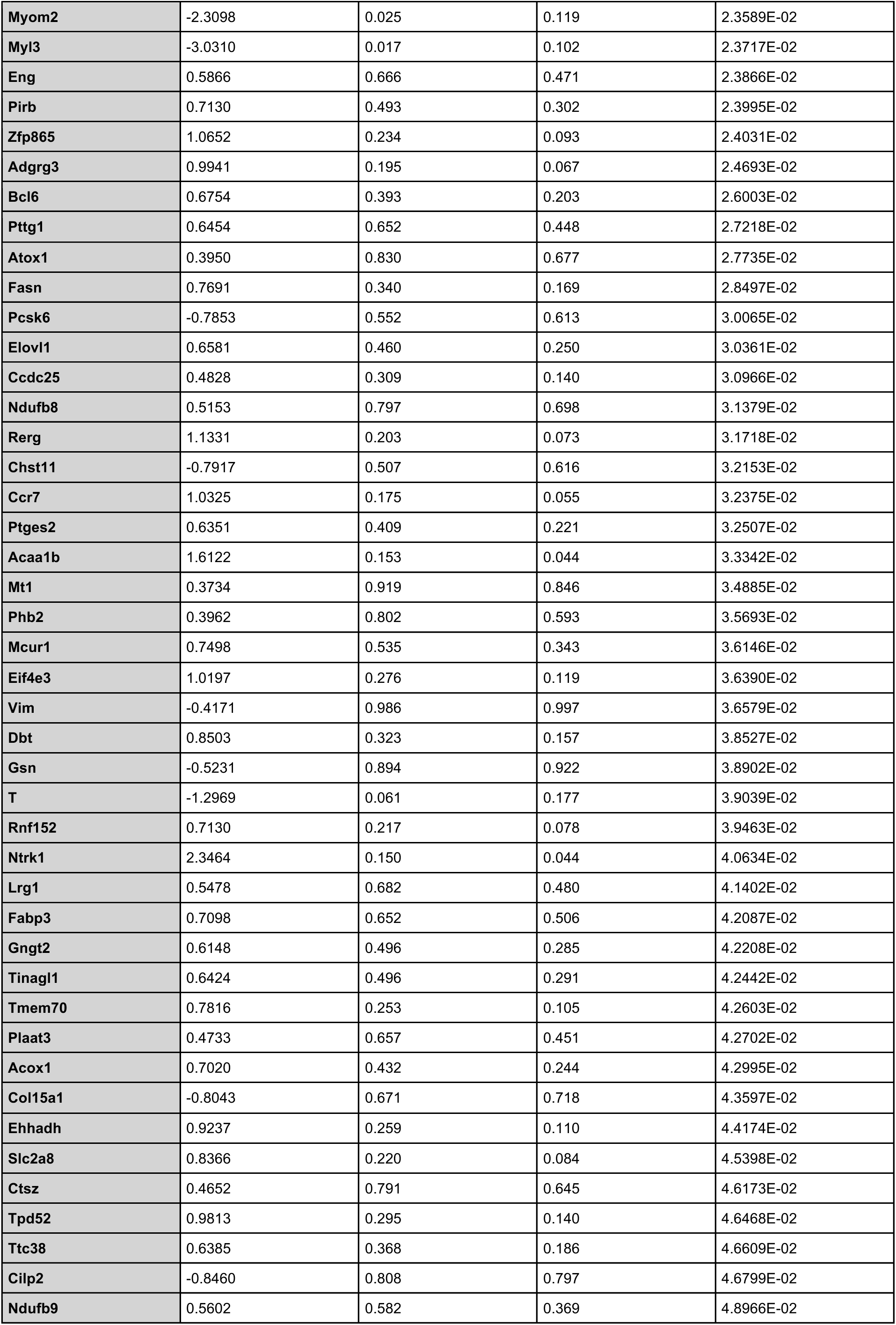
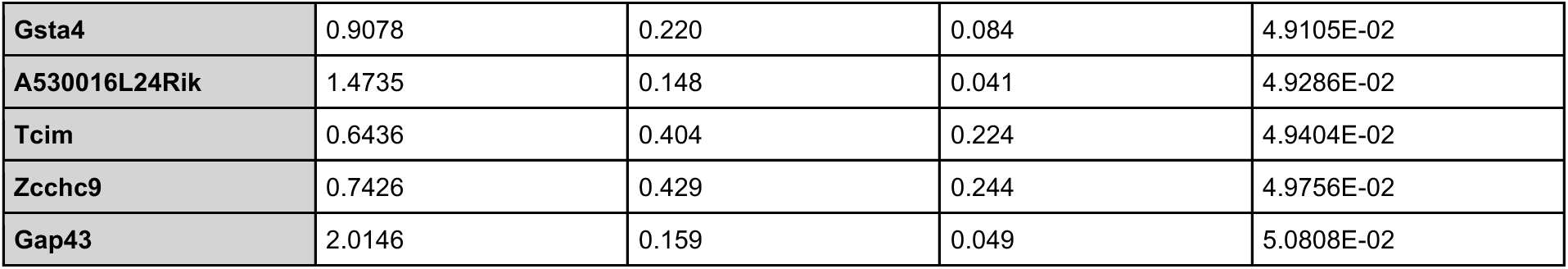
Differential gene expression of refined spatial analysis over intervertebral disc regions excluding high-muscle expressing capture spots. Differential expression was performed using the non-parametric Wilcoxon Rank Sum test. For condition-specific comparisons (*Adgrg6*-cKO vs. wildtype), data were subset to remove capture spots enriched for muscle-related transcripts and normalized using standard log-normalization. Statistical significance was defined as a Bonferroni-adjusted *p*-value < 0.05. Only genes detected in at least 10% of spots in either comparison group with a minimum natural log fold change of 0.25 were included.

### *Adgrg6-cKO* Mutant Mice Display a Significant Reduction of ECM Genes in the Inner Annulus Fibrosus

To validate and spatially resolve the transcriptomic changes identified by spatial transcriptomics, we performed RNA *in situ* hybridization for a panel of ECM and cartilage-related genes found to be downregulated in *Adgrg6-cKO* mice. Genes fell into three broad categories based on the magnitude and spatial character of their expression changes. A first set — including *Comp*, *Chad*, and *Col27a1* — showed no obvious change in pattern or expression level between wild-type and *Adgrg6-cKO* sections (Fig. 2a–d’, 3e–f’). A second set — *Acan*, *Sox9*, and *Creb3l2* — showed markedly reduced but detectable expression in *Adgrg6-cKO* mice relative to wild-type controls (Fig. 3a–d’, g–h’). A third set of genes — *Grep1*, *Frzb*, *Fmod*, *Col6a2*, and *Col14a1* — was profoundly decreased or nearly undetectable in *Adgrg6-cKO* sections despite robust expression in wild-type tissue (Fig. 2e–n’).

**Figure 2.**
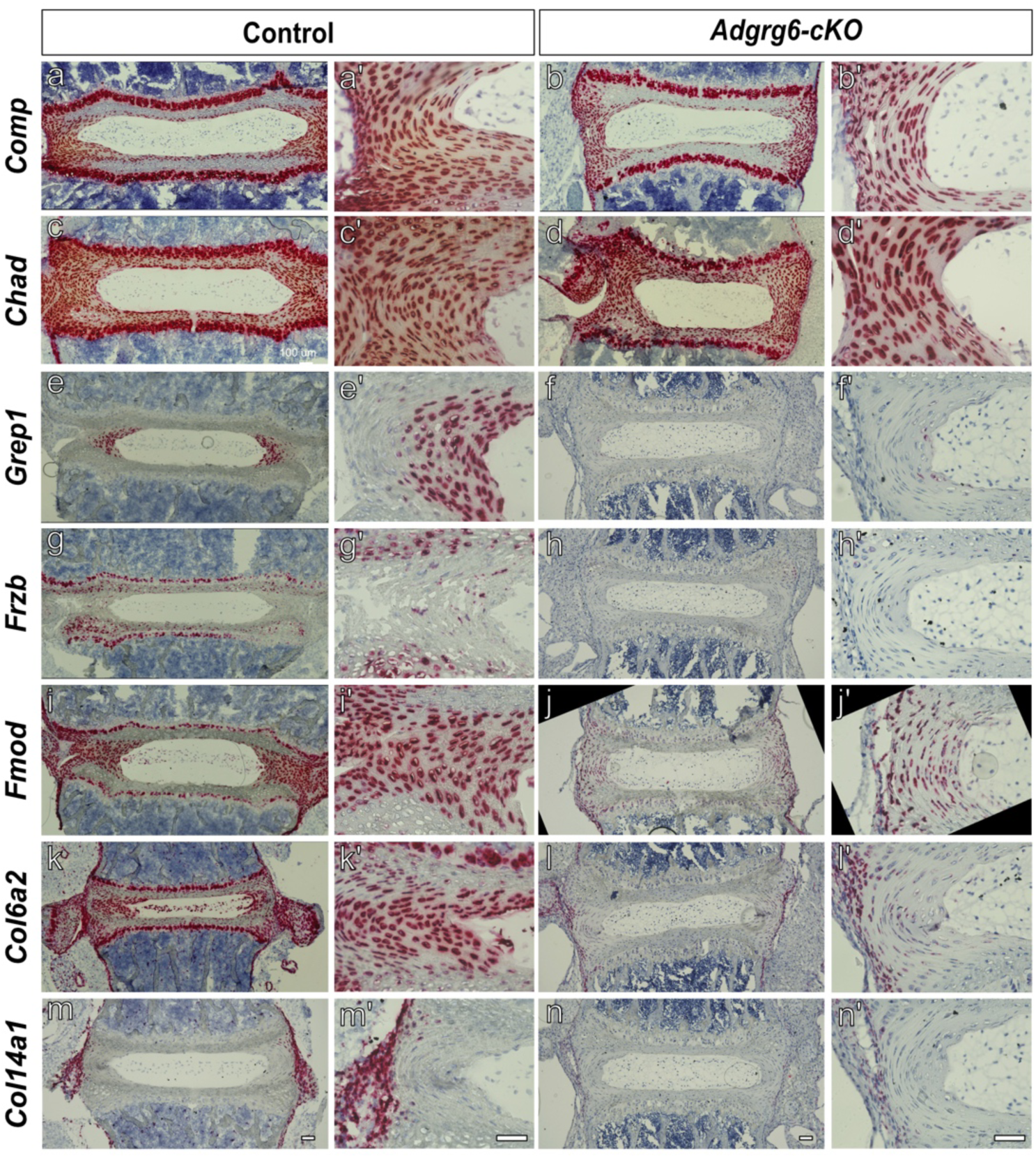
Genotype-specific expression of key ECM-regulatory genes in the intervertebral disc. RNA *in situ* hybridization (RNAscope) was used to validate the expression of extracellular matrix-associated genes identified as significantly decreased in the spatial transcriptomic analysis. The following genes were examined in the intervertebral disc (IVD): *Comp*, *Chad*, *Grep1*, *Frzb*, *Fmod*, *Col6a2*, and *Col14a1* (a-n). Representative images demonstrate gene-specific punctate signals within the intervertebral disc (a-n) and higher-magnification views of the annulus fibrosus (a’-n’). Columns correspond to individual genes as follows: *Comp* (a-b’), *Chad* (c-d’), *Grep1* (e-f’), *Frzb* (g-h’), *Fmod* (i-j’), *Col6a2* (k-l’), and *Col14a1* (m-n’) assessed in controls (a, c, e, g, i, k, m) and *Adgrg6-cKO* mutant (b, d, f, h, j, l, n) mice. Images are representative of *n ≥ 2* biologically independent samples per genotype. Scale bars = 100 µm in (a-n) and 50 µm (a’-n’).

**Figure 3.**
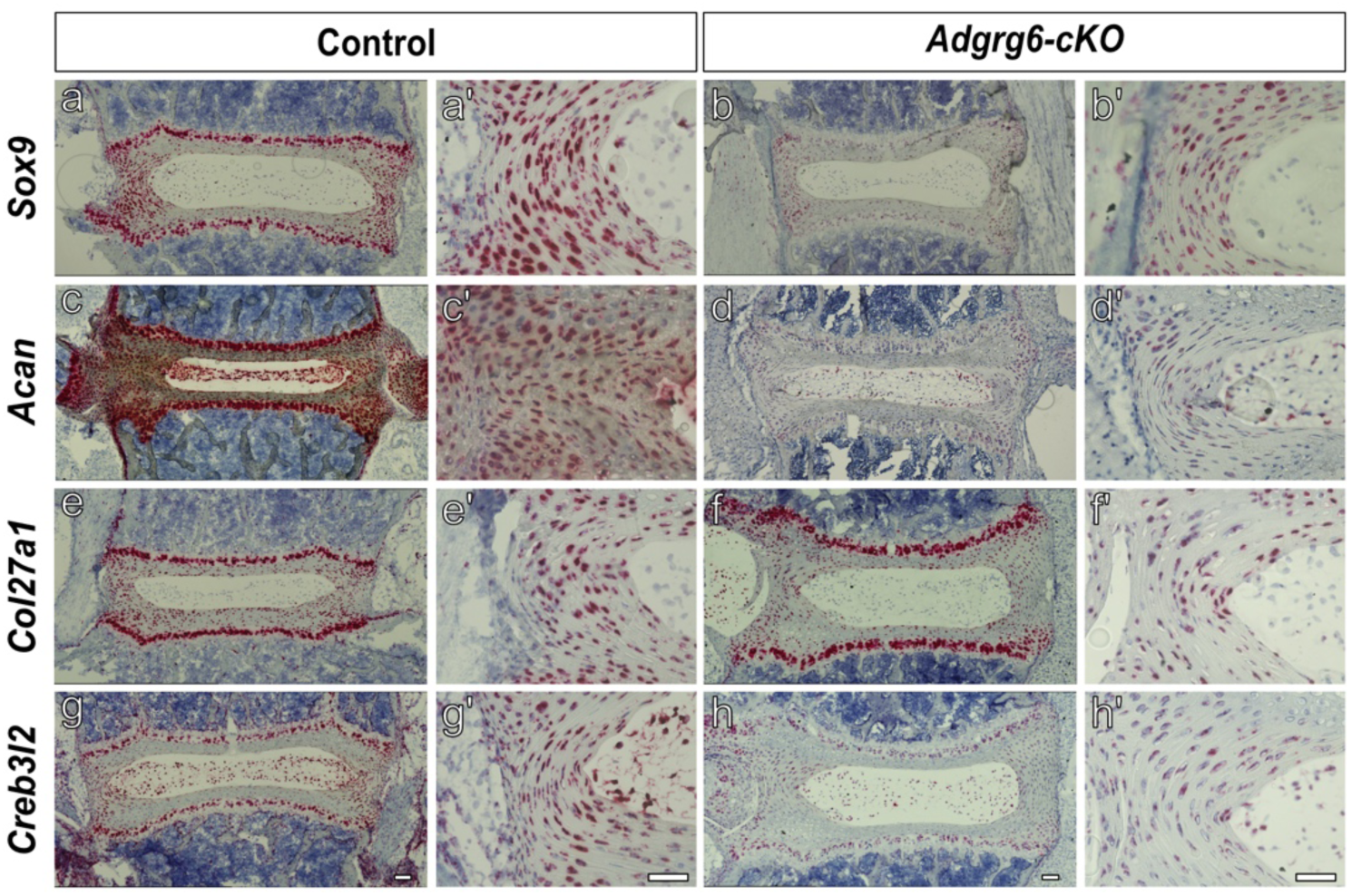
Expression of *Sox9* and *Sox9*-related genes in the intervertebral disc across genotypes. RNA *in situ* hybridization (RNAscope) was performed to validate spatial transcriptomic findings and examine the expression of select *Sox9* and *Sox9*-related genes within the intervertebral disc. Expression of *Sox9* (a-b’), *Acan* (c-d’), *Col27a1* (e-f’), and *Creb3l2* (g-h’) were evaluated. Representative images demonstrate gene-specific punctate signals within the IVD (a-h), along with corresponding higher-magnification views of the annulus fibrosus (a’-h’) in wild-type controls (a, c, e, g) and *Adgrg6-cKO* (b, d, f, h). Images are representative of *n* ≥ 2 biologically independent samples per genotype. Scale bars = 100 µm in (a-h) and 50 µm (a’-h’).

Among this third set, *Grep1* (also known as *1520401A03Rik* in mouse) showed the most pronounced downregulation in the spatial transcriptomics dataset (average logFC = −3.96) and a striking spatial specificity by *in situ* analysis confined to the inner annulus fibrosus (AF) in wild-type mice (Fig. 2e’), which was almost entirely absent in *Adgrg6-cKO* mutants (Fig. 2f’). *Frzb/Sfrp3*, a secreted Wnt antagonist, was strongly expressed in the cartilaginous endplate with minor expression in the AF of wild-type spines and was severely decreased in *Adgrg6-cKO* mice (Fig. 2g-h’). *Col6a2* and *Fmod* were broadly expressed throughout the AF in wild-type sections, with expression severely reduced in mutants (Fig. 2i-l’). *Col14a1* was strongly expressed in a thin layer of paraspinal (periosteum-like) tissue surrounding the length of the spine (Fig. 2m,m’; Supp. Fig. 5) and this expression was largely absent in *Adgrg6-cKO* mice (Fig. 2n, n’, Supp. Fig. 5). Together, these spatial expression patterns reveal that *Adgrg6* loss affects transcriptional programs across multiple discrete tissue compartments within and surrounding the IVD — including the inner AF, cartilaginous endplate, and paraspinal connective tissue — rather than reflecting a uniform suppression of ECM gene expression.

Notably, several of the ECM genes downregulated in spatial transcriptomics analysis of *Adgrg6-cKO* mice, including *Acan*, *Matn3*, *Col9a3*, and *Creb3l2*, have been identified as direct SOX9 targets by ChIP-seq in mouse chondrocytes ^36^ (Supp. Fig. 3). Consistent with this, *in situ* hybridization confirms that *Sox9*, *Acan*, and *Creb3l2* display overlapping expression in the endplate and AF (Fig. 3a-d’, g-h’) and are also reduced in *Adgrg6-cKO* mutants. *Col27a1*, despite being identified as a gene downregulated in *Adgrg6-cKO* mouse IVD by spatial transcriptomics, was not confirmed as reduced by *in situ* analysis (Fig. 3e-f’), which may reflect a subtle change below the sensitivity of the *in situ* hybridization approach. Collectively, these findings confirm that *Adgrg6* is required to sustain *Sox9* expression in the IVD and further suggest that SOX9 in turn drives transcription of a broad set of ECM targets with distinct spatial distributions across the AF and paraspinal tissues.

### Sox9 Occupies Accessible Chromatin Regulatory Elements Within the *Adgrg6* Locus

Our data above demonstrate that *Adgrg6* loss reduces *Sox9* expression in the IVD, while prior work has shown that postnatal *Sox9* deletion significantly reduces *Adgrg6* expression in the same tissue ^30^. Together with evidence that *Adgrg6*-dependent CREB signaling is required for maintenance of SOX9 protein expression ^21^, these observations support a self-regulatory feedforward loop to sustain gene expression in the IVD. However, whether SOX9 directly occupies regulatory elements at the *Adgrg6* locus has not been established.

To test this, we asked whether SOX9 binds accessible chromatin regions within the *Adgrg6* locus in IVD-resident cells. We used fluorescence-activated cell sorting (FACS) to isolate SOX9-EGFP⁺ cells from the IVDs of P7 *Sox9ires^EGFP/+^* embryos (Fig. 4a–b) and profiled chromatin accessibility genome-wide by ATAC-Seq^37^. To identify direct SOX9 occupancy at accessible sites, we performed SOX9 and IgG control Cut&Run-Seq on the same cell population ^38^. ATAC-Seq identified multiple accessible regions within both the *Col2a1* locus which is a well-established SOX9 transcriptional target ^39^ used here as a positive control and the *Adgrg6* locus (Fig. 4c–d). Cut&Run-Seq confirmed SOX9 occupancy at several of these ATAC peaks, with the strongest enrichment at the promoter and 5′ coding region of *Col2a1* and at a peak within the second intron of *Adgrg6*, a region with high vertebrate evolutionary conservation (Fig. 4d). These findings demonstrate that SOX9 directly binds putative regulatory elements at the *Adgrg6* locus in IVD-resident cells, establishing a transcriptionally direct arm of the *Adgrg6–Sox9* feedforward loop and providing a molecular mechanism through which SOX9 sustains *Adgrg6* expression to promote postnatal spinal homeostasis.

**Figure 4.**
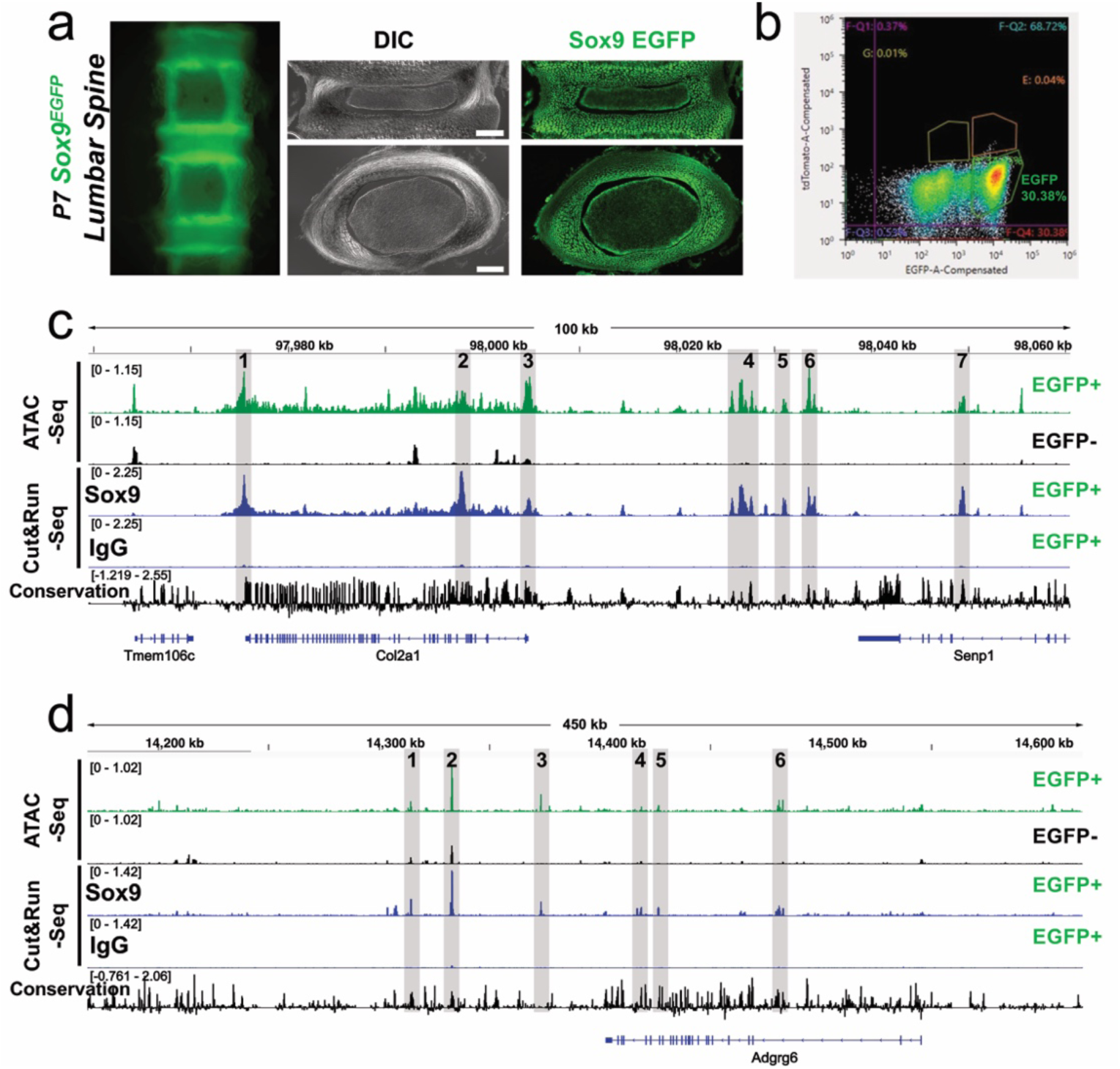
Sox9 directly binds to putative regulatory elements surrounding the *Adgrg6* locus. Direct fluorescence of the lumbar spine/IVD region isolated from postnatal day 7 (P7) *Sox9^EGFP^* mouse (a) (Left). DIC image or direct fluorescence of cryosectioned lumbar intervertebral disc isolated from P7 *Sox9^EGFP^* mouse (a) (Right). The top panel is a coronal cryosection of the IVD; the bottom panel is a transverse cryosection of the IVD (a). Scale bar equals 200 microns. FACS profile of *Sox9-*EGFP+ cells isolated from the developing spine/IVD of P7 *Sox9^EGFP^* mouse (b). Analysis of chromatin regulation *Sox9-*EGFP+ cells isolated from spine/IVD of P7 *Sox9^EGFP^* mouse shown as Integrated Genome Views (IGV) of ATAC-Seq, Sox9-Cut&Run-Seq, or IgG-Cut&Run-Seq (c, d), demonstrating SOX9 binding to several differentially accessible regulatory regions surrounding the *Col2a1* (c) and *Adgrg6* loci (d). Regions of overlap between ATAC-Seq peaks and SOX9 enriched-Cut&Run peaks are indicated (gray shadows), with evolutionary conservation in 60 vertebrate genomes displayed.

### Genetic Interactions Between *Adgrg6* and *Sox9* Increase Scoliosis Penetrance and Severity in Mice

SOX9 dosage sensitivity is well established, with partial reductions in SOX9 levels producing measurable differences in target gene expression and phenotypic outcomes ^40^. Given the feedforward co-regulatory relationship between *Adgrg6* and *Sox9* demonstrated above, and the independent association of both loci with human AIS ^8,21,24,31–33^, we hypothesized that reducing *Sox9* dosage in the spine would exacerbate scoliosis onset and severity in *Adgrg6-cKO* mice. To test this, we utilized a hypomorphic *Sox9* allele encoding an in-frame Asp272 deletion within the Sox9 TAM domain (hereafter *Sox9^del^*). Homozygous *Sox9^del^* mutants are viable and display mild skeletal phenotypes reminiscent of acampomelic dysplasia — including bilateral loss of floating ribs and late-onset scoliosis (>5 months) — without craniofacial malformations or respiratory distress^33^. We generated *Col2Cre; Adgrg6^f/+^;Sox9^del/+^* and *Adgrg6 ^f/+^*;*Sox9^del/+^* breeders and performed longitudinal microCT imaging and analysis of spine morphology at P40 and P120 across all genotypes. Body weights remained comparable across groups at both time points, indicating no general health differences that could confound interpretation of skeletal phenotypes (Supp. Fig. 6).

Genetic interactions between *Adgrg6-cKO* and *Sox9^del^* alleles were evident at both timepoints. Wild-type controls displayed straight spines with normal thoracic architecture at P40 and P120 (Fig. 5a, a’). Consistent with prior work, homozygous *Sox9^del^* mutants exhibited bilateral loss of the T13 floating ribs (90%; n=11) (Fig. 5b, b’, asterisk; Supp. Fig. 7a), but did not develop scoliosis at either timepoint (0%; n=11; Fig. 5e, f). *Adgrg6-cKO* single mutants showed progressive single thoracic scoliosis by P120 (20%; n=10) (Fig. 5c, c’, f; Table 3). In contrast, *Adgrg6-cKO*; *Sox9^del/del^* double mutants demonstrated a marked increase in scoliosis penetrance as early as P40 (54.5%; n=11; Fig. 5d, e; Table 3), which further increased by P120 (63.6%; n=11), with 9% of double mutants displaying double thoracic curves (Fig. 5d’, f; Table 3). A gene-dosage effect was also observed in compound *Adgrg6-cKO*; *Sox9^del/+^* heterozygous mutants, which exhibited increased scoliosis incidence (30%; n=11) and severity (20%; n=11) relative to *Adgrg6-cKO* alone (Supp. Fig. 7d), demonstrating that a single copy of the *Sox9* microdeletion allele is sufficient to modify scoliosis susceptibility. Notably, transheterozygous *Col2Cre; Adgrg6^f/+^*; *Sox9^del/del^* mice did not develop scoliosis by P120 (0%; n=12), implying that heterozygous loss of *Adgrg6* is not sufficient to sensitize the spine to reduced *Sox9* gene dosage. Fisher’s exact tests with Monte Carlo simulation (B = 5000) for pairwise severity counts at P120 confirmed significant genotype-dependent differences, with *Adgrg6-cKO*; *Sox9^del/del^* double mutants differing significantly from all other experimental genotypes and wild-type controls (Bonferroni-adjusted p < 0.01; Supp. Fig. 7d). Collectively, these data demonstrate that dose-dependent reduction in *Sox9* function is sufficient to enhance both the penetrance and severity of scoliosis in the *Adgrg6-cKO* background, establishing combined *Adgrg6–Sox9* insufficiency as a tractable model of polygenic scoliosis susceptibility.

**Figure 5.**
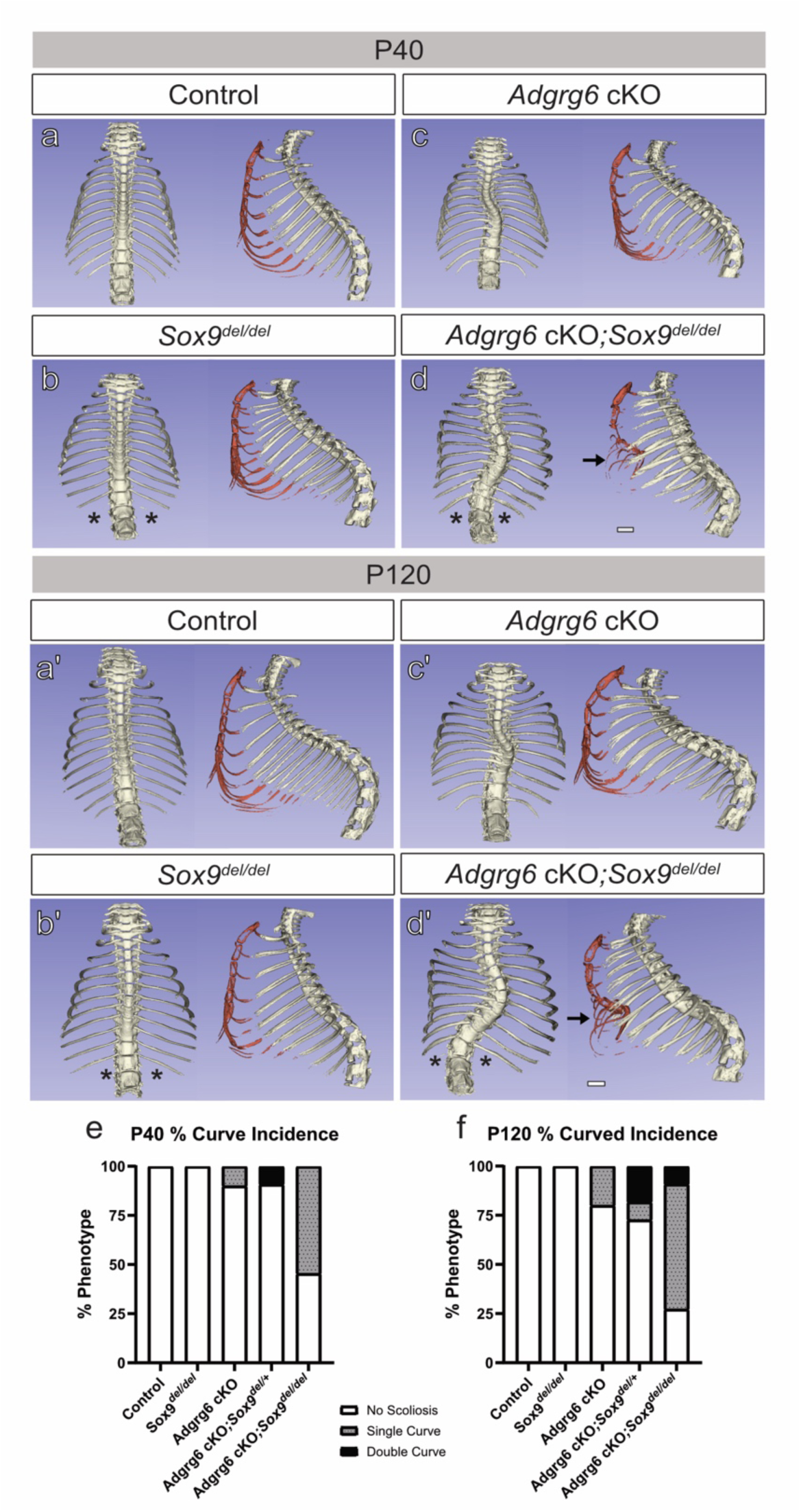
Genetics interaction between *Adgrg6* and *Sox9* mutations contribute to increased scoliosis penetrance and severity. Reconstructed microCT images from control (a-a’), *Sox9^del/del^* (b-b’), *Adgrg6-cKO* (c-c’), and *Adgrg6-cKO ;Sox9^del/del^* (d-d’) reveal skeletal defects at postnatal day (P)40 and (P)120. Wildtype mice show straight spines and natural barreling of the chest cavity at both time points (n=10) (a, a’). *Sox9^del/del^* mice present with loss of T13 rib pair (asterisk, 90%) and no scoliosis at either time points (0%, n= 11) (b, b’). *Adgrg6-cKO* mice show single thoracic scoliosis in 20% of animals at P120 (n=10) (c, c’). *Adgrg6-cKO ;Sox9^del/del^* mice presented with pectus excavatum (black arrow, 45.5%), complete loss of T13 rib pair (asterisk, 100%), and increased scoliosis incidence and severity as evidenced by no scoliosis (27.3%), single thoracic curvatures (63.6%), double thoracic curvatures (9.1%), and scoliosis concurrent with pectus excavatum in 36.4% of mice at P120 (n=11) (d, d’). *Adgrg6-cKO*, *Adgrg6-cKO ;Sox9^del/+^*, and *Adgrg6-cKO ;Sox9^del/del^* mice showed age-dependent progression in scoliosis penetrance and severity (e-f). Scale bar = 2 mm.

**Table 3.**
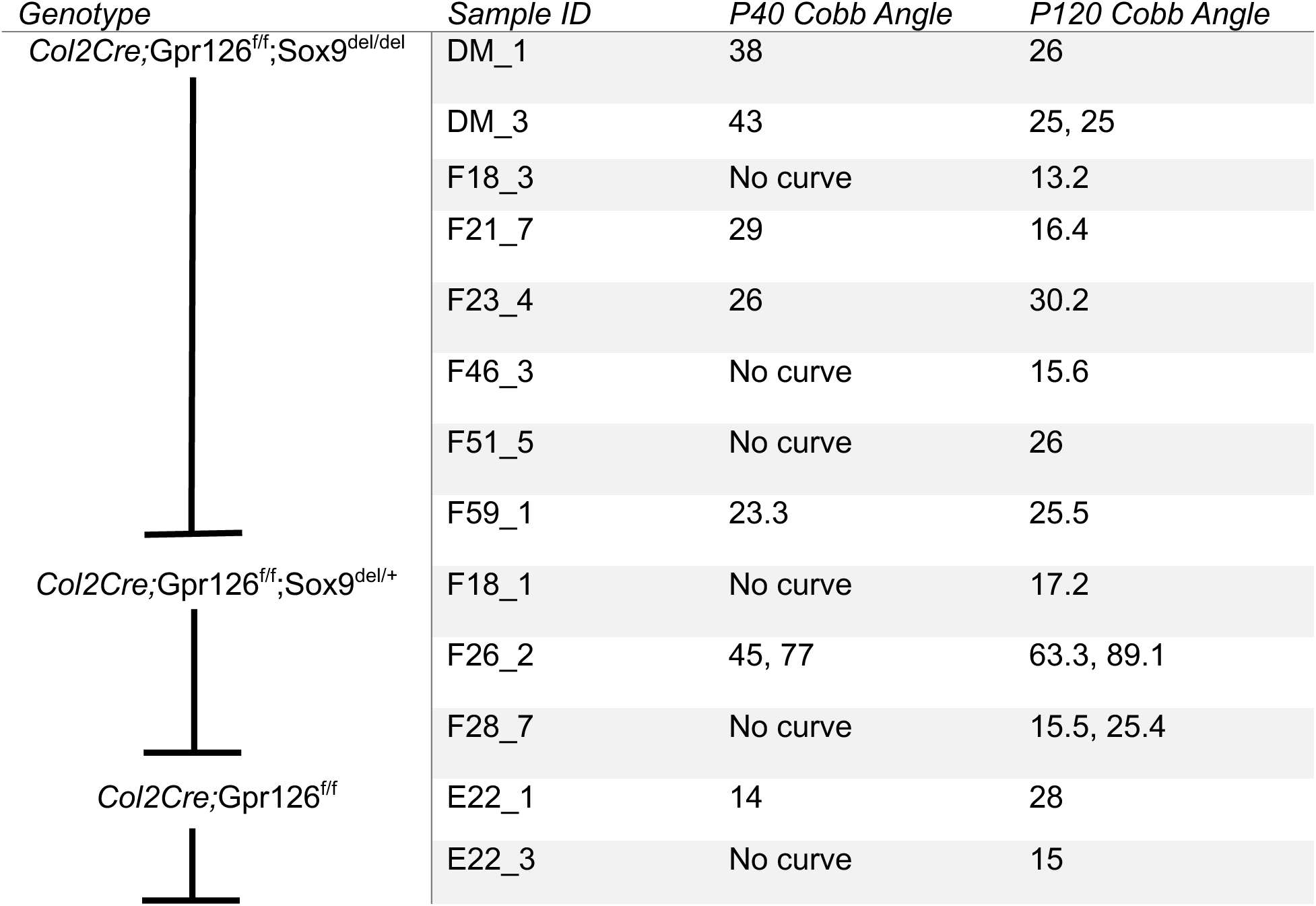
Longitudinal Cobb angle measurements in mice presenting with thoracic scoliosis. Cobb angles were quantified for all experimental genotypes in the dataset that had scoliosis that exhibited a measurable thoracic curvature (>10 degrees). “No curve” denotes a curve less than 10 degrees or the absence of noticeable scoliosis. At P40, *Col2Cre;Adgrg6^f/f^;Sox9^del/del^* presented with five single curves and three *no curves*. By P120, previously straight spines developed a single curve, resulting in a total of seven single curves and one double curve. Double thoracic curvatures are denoted by two Cobb angles (e.g. 45, 77).

### *Adgrg6-Sox9* Genetic Interactions Increase Pectus Excavatum Penetrance in Mice

Pectus excavatum (PE) is a congenital chest wall deformity characterized by inward depression of the sternum and ribs ^41^. We previously reported PE in Adgrg6-cKO mutant mice, occurring either independently or concurrently with AIS-like scoliosis ^28^. In the present cohort, PE was observed in *Sox9^del^* mutants and all double mutant combinations, but was absent in *Adgrg6*-*cKO* single mutants (Supp Fig. 7b, c). *Sox9^del^*homozygotes displayed low PE penetrance at P120 (18.2%; n=11) without concurrent scoliosis (Supp. Fig. 7b, c).

A clear gene-dosage effect on PE penetrance was evident across double mutant genotypes. Transheterozygous *Col2Cre; Adgrg6^f/+^; Sox9^del/+^* mice exhibited similarly low PE penetrance to *Sox9^del^* homozygotes (16.7%; n=12; Supp. Fig. 7b). *Adgrg6-cKO; Sox9^del/+^* compound heterozygotes showed a modest increase in PE incidence (30%; n=11), while *Adgrg6-cKO; Sox9^del/del^* double mutants displayed the highest PE penetrance (54.5%; n=11; Supp. Fig. 7b) and the greatest co-occurrence of PE and scoliosis (45.4%; n=11; Fig. 5d, d’, arrow; Supp. Fig. 7b, c). Fisher’s exact test for pairwise comparisons of PE incidence at P120 confirmed that only *Adgrg6-cKO; Sox9^del/del^* double mutants differed significantly from all other experimental genotypes and controls (p < 0.05; Supp. Fig. 7b). Concurrent incidence of PE and scoliosis at P120 showed no significant genotype-dependent differences (Supp. Fig. 7c), suggesting that these two phenotypes arise through at least partially independent pathogenic mechanisms. Taken together, these data indicate that the sternum and chest wall may be more sensitive to graded reductions in *Adgrg6* and *Sox9* function than the IVD and annulus fibrosus and further underscore the dose-dependent nature of the *Adgrg6–Sox9* genetic interaction in maintaining axial skeletal integrity.

### Loss of *Sox9* exacerbates the incidence of growth plate defects and ectopic cartilage regions in vertebrae

To evaluate if structural morphological changes in spinal tissues contributed to this increased susceptibility of scoliosis, P20 spines were analyzed using Alcian Blue Hematoxylin Eosin Orange-G staining (Supp Fig. 8a-d). We observed an increase of atypical growth plate extensions and ectopic cartilaginous regions in the vertebrae of *Adgrg6-cKO, Sox9^del^*, and *Adgrg6-cKO; Sox9^del/del^* double mutant spines, that were never observed in wild-type controls (Supp Fig. 8). To quantify the incidence and severity of this phenotype, we calculated the percentage of affected IVDs per spine and number of growth plate extensions per affected IVD (Supp. Fig. 8e, f). Consistent with prior reports of conditional *Adgrg6* deletion in IVD-associated tissues ^42^, *Adgrg6-cKO* mice exhibited growth plate defects with incomplete penetrance (33%; n = 3; Supp. Fig. 8c, e). In contrast, *Sox9^del^* spines showed a fully penetrant appearance of growth plate extensions with increased severity compared to *Adgrg6-cKO* (100%, n = 5; Supp. Fig. 8b, e, f). *Adgrg6-cKO; Sox9^del/del^* spines exhibited an increased severity of growth plate extensions and ectopic cartilage regions in the vertebrae compared to *Adgrg6-cKO* (85%, n = 6; Supp. Fig. 8d-f). Together, these findings suggest that reduction in *Sox9* expression is a major regulator that drives the growth plate defects. Moreover, these data suggest that defects in bone mineralization may be involved in the increased incidence and severity of AIS phenotypes in *Adgrg6-cKO; Sox9^del/del^* double mutant mice.

### Increased scoliosis incidence and severity are associated with decreased bone mineral density in thoracic vertebrae

Although conditional deletion of *Adgrg6* in osteoprogenitor cells does not cause scoliosis ^21^, *Adgrg6-cKO; Sox9^del/del^* double mutant mice showed severe growth plate extensions (Supp. Fig. 8), prompting us to examine whether combined *Adgrg6–Sox9* insufficiency alters vertebral microarchitecture. Bone density-mapped reconstructions with Otsu-based threshold analysis ^43^ of thoracic spines at P40 revealed a general reduction in bone mineralization in the high-density range (0.286–0.551 g/cm³) within the region of the scoliotic curve in double mutants compared to controls (Fig. 6a, d). To further characterize this, we segmented the T4 vertebra (Fig. 6a, d, arrow), as this region typically corresponds to the apex of thoracic scoliotic curves. Quantitative microCT analysis of T4 across all genotypes revealed significant reductions in both bone volume and bone mineral density (BMD) in *Adgrg6-cKO; Sox9^del/del^* mice compared to controls (p < 0.05; Fig. 6g, h). Otsu analysis further confirmed a significant reduction in high-density bone volume fraction in double mutants (p < 0.05; Fig. 6i), consistent with the density-mapped reconstructions of T4 vertebrae (Fig. 6c, f).

**Figure 6.**
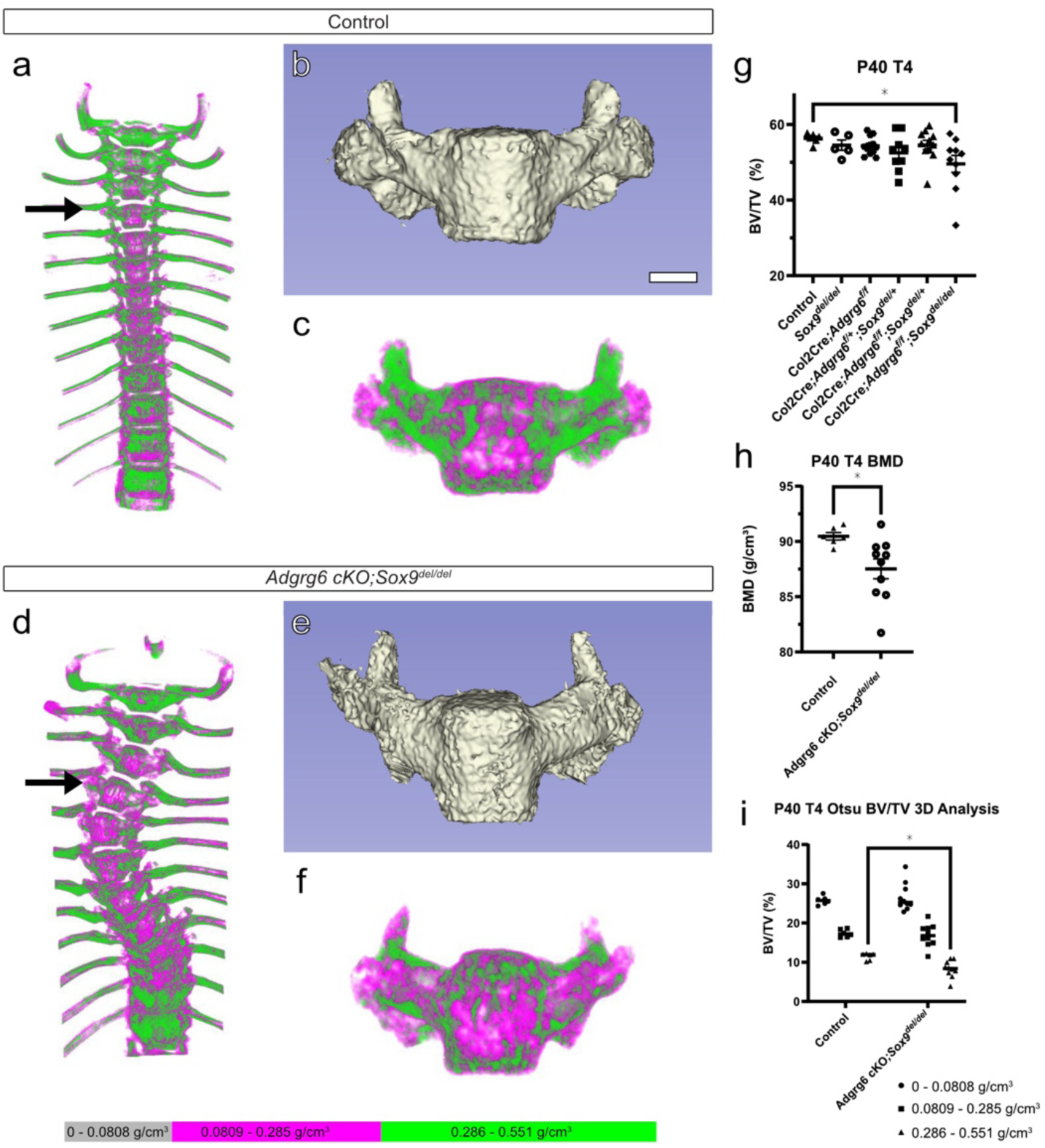
*Adgrg6-cKO ;Sox9^del/del^* double mutant mice exhibit microarchitectural alterations in a thoracic vertebra at P40. MicroCT analysis of thoracic vertebral bone architecture at P40. Density-mapped reconstructions of the thoracic spine reveal differences in overall mineral distribution between straight control and scoliotic *Adgrg6-cKO ;Sox9^del/del^* spines (a, d). Arrows indicate the T4 vertebra analyzed in subsequent panels. Three-dimensional segmentations of T4 illustrate overall vertebral morphology in control and double mutant mice (b, e). Corresponding density-mapped reconstructions of representative T4 vertebrae demonstrate altered mineral distribution throughout the vertebrae in double mutant mice (c, f). One way ANOVA and t-test analysis revealed a significant difference in bone volume and BMD in control and *Adgrg6-cKO ;Sox9^del/del^* T4 vertebrae (*p* < 0.05) (g, h). Two way ANOVA Otsu-based threshold 3D bone volume stratifying bone into low-(0–0.0808 g/cm³; grey), mid-(0.0809–0.285 g/cm³; magenta), and high-density (0.286–0.551 g/cm³; green) fractions revealed a significant loss of high-density distribution in double mutant mice (*p* < 0.05) (i). Each dot represents an individual mouse vertebrae, with mean ± SEM overlaid. Scale bar = 0.5 mm (b,c, e, f).

To determine whether these bone changes are specific to spinal regions susceptible to scoliosis, we assessed L5 vertebral microarchitecture in the same cohort at P40 (Supp. Fig. 9). MicroCT analysis of L5 revealed no significant differences in overall bone morphology or BMD across genotypes (Supp. Fig. 9g, h), consistent with representative cross-sectional images showing grossly normal inner bone architecture (Supp. Fig. 9a’–f’). However, Otsu analysis revealed a significant reduction in low-density bone volume in *Adgrg6-cKO; Sox9^del/del^* double mutants relative to controls (Supp. Fig. 9i), showing that subtle compositional differences are present in lumbar vertebrae which are not directly part of scoliotic regions. Collectively, these findings indicate that while mild alterations in bone composition extend throughout the spine, the most pronounced microarchitectural defects are concentrated in thoracic vertebrae, likely reflecting the combined effects of *Adgrg6–Sox9* insufficiency and the region-specific mechanical demands imposed by the onset and progression of scoliosis.

## DISCUSSION

Adolescent idiopathic scoliosis is best understood as a complex polygenic condition in which multiple susceptibility alleles of individually modest effect combine to cross a phenotypic threshold during the period of rapid pubertal growth. GWAS have reproducibly identified non-coding variants near *ADGRG6* and *SOX9* as AIS risk loci in human populations, yet the functional relationship between these two genes in the postnatal spine has remained undefined. Here, we integrate spatial transcriptomics, chromatin accessibility profiling, SOX9 chromatin occupancy, and multi-allelic mouse genetics to demonstrate that *Adgrg6* and *Sox9* operate in a feedforward regulatory circuit in which each gene has a self-reinforcing role in co-regulation to maintain a broad program of ECM gene expression in the IVD and paraspinal connective tissues. Critically, we show that combinatorial reduction of *Adgrg6* and *Sox9* function, which individually produce incompletely penetrant scoliosis phenotypes, dramatically increases both the incidence and severity of AIS-like pathology in mice. These findings provide a mechanistic framework for understanding how co-variation at the *ADGRG6* and *SOX9* loci may interact to modulate scoliosis susceptibility in human populations.

The central molecular finding of this study is the identification of a feedforward loop between *Adgrg6* and *Sox9* in IVD and paraspinal tissues. Prior work established that *Adgrg6*-dependent CREB signaling is required for maintenance of SOX9 protein expression in the IVD ^21^, and that postnatal *Sox9* deletion significantly reduces *Adgrg6* expression in the same tissue ^30^. The present study extends this bidirectional relationship by demonstrating, first, that *Adgrg6* loss reduces *Sox9* transcript levels and putative SOX9-target gene expression in the IVD, and second, that SOX9 directly occupies several more accessible and conserved chromatin regions surrounding the *Adgrg6* locus, including a region of the second intron of this gene in IVD-resident cells. Together, these observations define a self-reinforcing transcriptional circuit in which *Adgrg6* signaling sustains SOX9 protein levels, while SOX9 in turn directly promotes *Adgrg6* transcription by directly binding to conserved regulatory elements. Feedforward architectures of this kind confer robustness to transient perturbations under normal conditions but render a system acutely sensitive to sustained or combinatorial disruption, which is consistent with the dose-dependent exacerbation of scoliosis we observe in *Adgrg6-cKO; Sox9^del/del^* compound mutants. We propose because *Adgrg6* can act as a force-sensing mechanoreceptor ^25^ that this circuit represents a core regulatory node for postnatal IVD homeostasis in response to stress and strain of the spine, and that partial disruption of either arm through non-coding variants affecting regulatory element activity in the IVD and paraspinal tissues or through coding variants affecting protein function may be sufficient to shift an individual toward AIS susceptibility when combined with additional genetic or environmental risk factors.

The spatial transcriptomic and *in situ* hybridization data reveal that *Adgrg6* loss does not produce a uniform suppression of ECM gene expression in the IVD but rather affects discrete subsets of targets with distinct spatial distributions and with apparent sensitivity thresholds, which is a modality well-established in response to alterations in SOX9 dosage and its regulatory role on target gene expression ^40^. Using RNA *in situ* hybridization we showed that many genes such as *Comp*, *Chad*, and *Col27a1* had no detectable change in expression, while *Acan*, *Sox9*, and *Creb3l2* were reduced but the patterning of expression was maintained, and a third set — *Grep1*, *Frzb*, *Fmod*, *Col6a2*, and *Col14a1* — were profoundly depleted. This graded response likely reflects differences in promoter architecture, SOX9 occupancy affinity, or dependence on co-activators whose activity is selectively disrupted downstream of *Adgrg6*. Among the most severely downregulated targets, *Grep1* is of particular interest. *Grep1* encodes a secreted ECM protein with homology to human Glycine-rich extracellular protein 1 and has been implicated in regulating GDF15 expression ^44^. Moreover, *Grep1* shows highly specific expression in the inner AF of wild-type mice, and near-complete absence in *Adgrg6-cKO* mutants, predictive of a spatially restricted role in maintaining inner AF identity that warrants further investigation. *Frzb (Sfrp3)*, a secreted Wnt antagonist expressed in the cartilaginous endplate, is similarly notable given emerging evidence for Wnt pathway dysregulation of cartilage homeostasis and correlated expression with disc degeneration ^45^. *Col14a1* expression in a periosteum-like layer surrounding the spine, and its marked reduction in *Adgrg6-cKO* mice, extends the transcriptional consequences of *Adgrg6* loss beyond the IVD proper to the paraspinal connective tissue envelope — a compartment increasingly recognized as relevant to spinal biomechanics and AIS pathogenesis ^21,23^. Collectively, the spatial heterogeneity of these expression changes underscores that *Adgrg6* coordinates transcriptional programs across multiple anatomically distinct tissue compartments, and that its loss disrupts IVD homeostasis in a spatially and quantitatively nuanced manner.

A key translational implication of this work is that the *ADGRG6* and *SOX9* GWAS loci, while typically analyzed as independent AIS risk factors, may in fact converge on a shared regulatory pathway whose output is sensitive to combinatorial genetic perturbation. We identified a putative enhancer in the second intron of mouse *Adgrg6*. This is compelling given that the strong *ADGRG6* AIS-associated variant (rs6570507) is also within the 2^nd^ intron of human *ADGRG6* ^24^. This suggests that these putative enhancer regions are instructive for spatially defined regulation of gene expression within the AF and paraspinal tissues. Future studies employing reporter assays or CRISPR-based mutagenesis of these putative regulatory elements in human cell lines or *in vivo* models will be needed to test this hypothesis directly. More broadly, our double mutant data provide a functional analog to the proposed polygenic burden model of AIS ^3^. Our genetic interactions demonstrated that neither heterozygous *Adgrg6* nor heterozygous *Sox9^del^* loss alone is sufficient to cause scoliosis, but their combination on the *Adgrg6-cKO* background crosses a phenotypic threshold in a dose-dependent manner. This mirrors the epidemiological observation that AIS risk is not determined by any single variant but by the aggregate burden of partially penetrant alleles across interacting pathways ^4,10^, and positions *Adgrg6–Sox9* compound insufficiency as a tractable experimental model for dissecting the genetic architecture of polygenic scoliosis susceptibility.

The microarchitectural bone phenotype observed in *Adgrg6-cKO; Sox9^del/del^* double mutants provides additional insight into the mechanisms by which *Adgrg6–Sox9* insufficiency promotes scoliosis progression. The partial reduction in BMD in double mutants, and its spatial restriction to thoracic rather than lumbar vertebrae, argues against a primary osteoblast defect, consistent with the prior demonstration that osteoprogenitor-specific *Adgrg6* deletion does not cause scoliosis ^21^. Instead, we favor a model in which failure of cartilage endplate maintenance, evidenced by growth plate defects and ectopic cartilage formation within the vertebrae body, secondarily compromises vertebral bone quality. Supporting this model, ablation of *Adgrg6 Adgrg6* in osteochondral progenitor cells causes delayed formation of the secondary ossification center and growth palate dysplasia in the long bone ^46^, which may alter bone mechanics in the vertebrae. Furthermore, *Adgrg6* loss in the cartilaginous endplate elevates STAT3 activation and MMP13 expression alongside reduced SOX9 expression, promoting growth plate herinations that resemble Schmorl’s nodes ^42^. SOX9 is a well-established regulator of cartilage endplate formation and IVD compartment identity ^30^, and its combined reduction with *Adgrg6* loss may further destabilize growth plater integrity below the threshold required for normal vertebral development. Such growth plate disruption could impose asymmetric mechanical loading across the thoracic spine, driving the aberrant bone remodeling and curve progression characteristic of AIS ^47^. The growth plate defects observed in *Sox9^del^* and double mutant spines at P20 support this interpretation and raise the possibility that endplate pathology is an early initiating event in the disease cascade, rather than a secondary consequence of established curvature.

In summary, this study demonstrates that *Adgrg6* and *Sox9* are co-regulated through a self-regulating feedforward transcriptional circuit, and that their combinatorial insufficiency is sufficient to model the polygenic architecture of AIS susceptibility in mice. By establishing a direct molecular link between two of the most replicated AIS GWAS loci, these findings move beyond statistical co-association to provide a mechanistic basis for understanding how non-coding variation at *ADGRG6* and *SOX9* might interact to modulate disease risk in human populations. More broadly, this work supports an emerging view of AIS as a condition arising from the convergence of subtle defects across interconnected regulatory circuits governing IVD homeostasis, connective tissue integrity, and spinal biomechanics — and suggests that the *Adgrg6–Sox9* feedforward loop represents one functionally validated node within this broader network. Identifying additional interacting loci within and beyond this circuit and determining how their combined perturbation maps onto the spectrum of AIS severity observed in human patients, remains an important goal for future investigation.

## Supplemental Figures

**Supplemental Figure 1.**
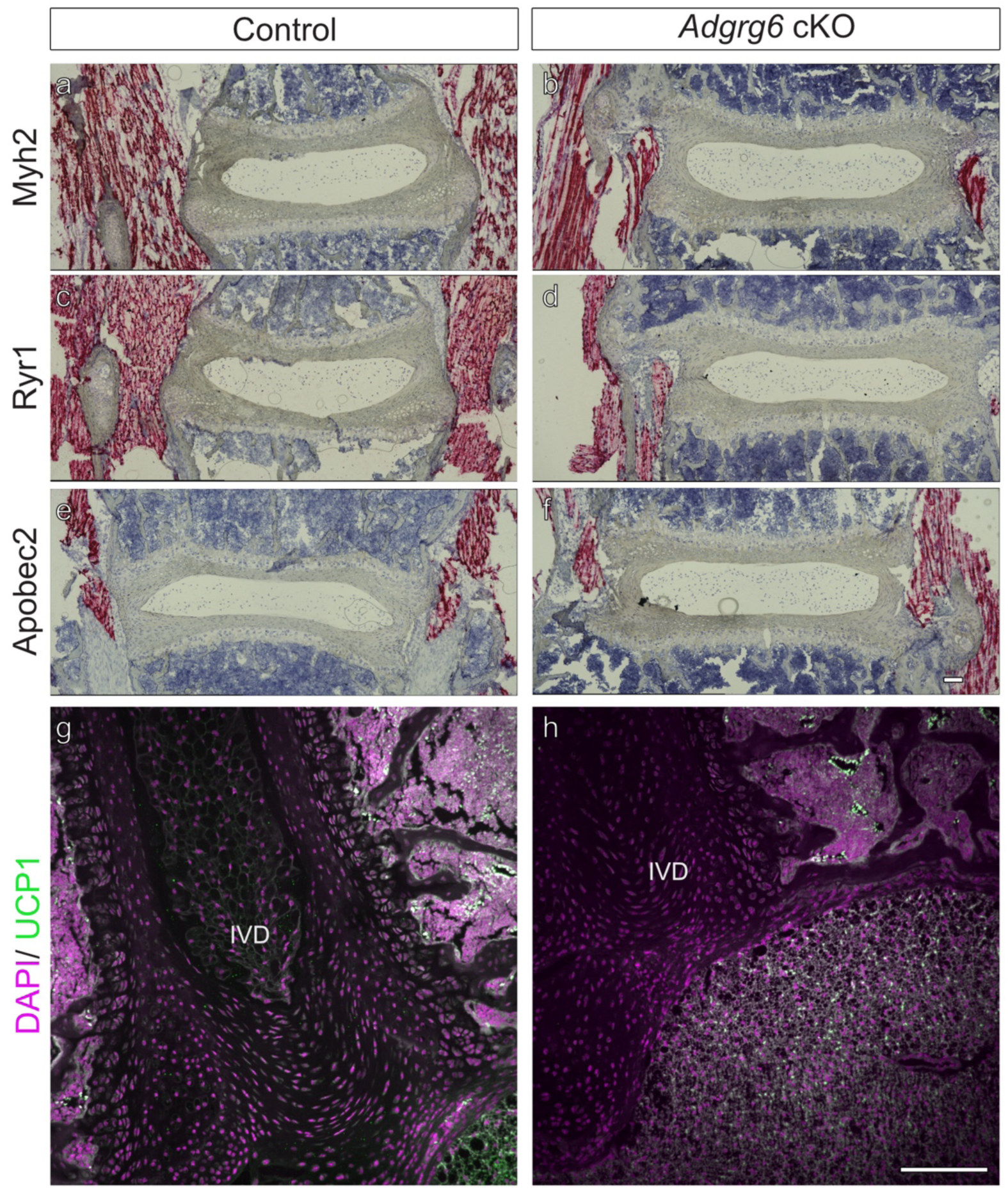
RNA *in situ* analysis of muscle and adipose-associated transcripts. Representative RNAScope images comparing control and *Adgrg6-cKO* spines probed for muscle genes *Myh2* (a-b), *Ryr1* (c-d), and *Apobec2* (e-f). Red puncta indicate transcript signal, which is robust in the paraspinal skeletal muscle adjacent to the vertebral column in both genotypes. No detectable expression of *Myh2* or *Ryr1* is observed within the IVD (a-d). In contrast, *Apobec2* shows very low-level signal in the annulus fibrosus. UCP1 expression (green) in the paraspinal adipose and muscle tissues in wild-type control (g) and *Adgrg6-cKO* mutant (h) mice, contrasted with nuclei staining to highlight the intervertebral disc (IVD) (DAPI/magenta). Scale bars = 100 um.

**Supplemental Figure 2.**
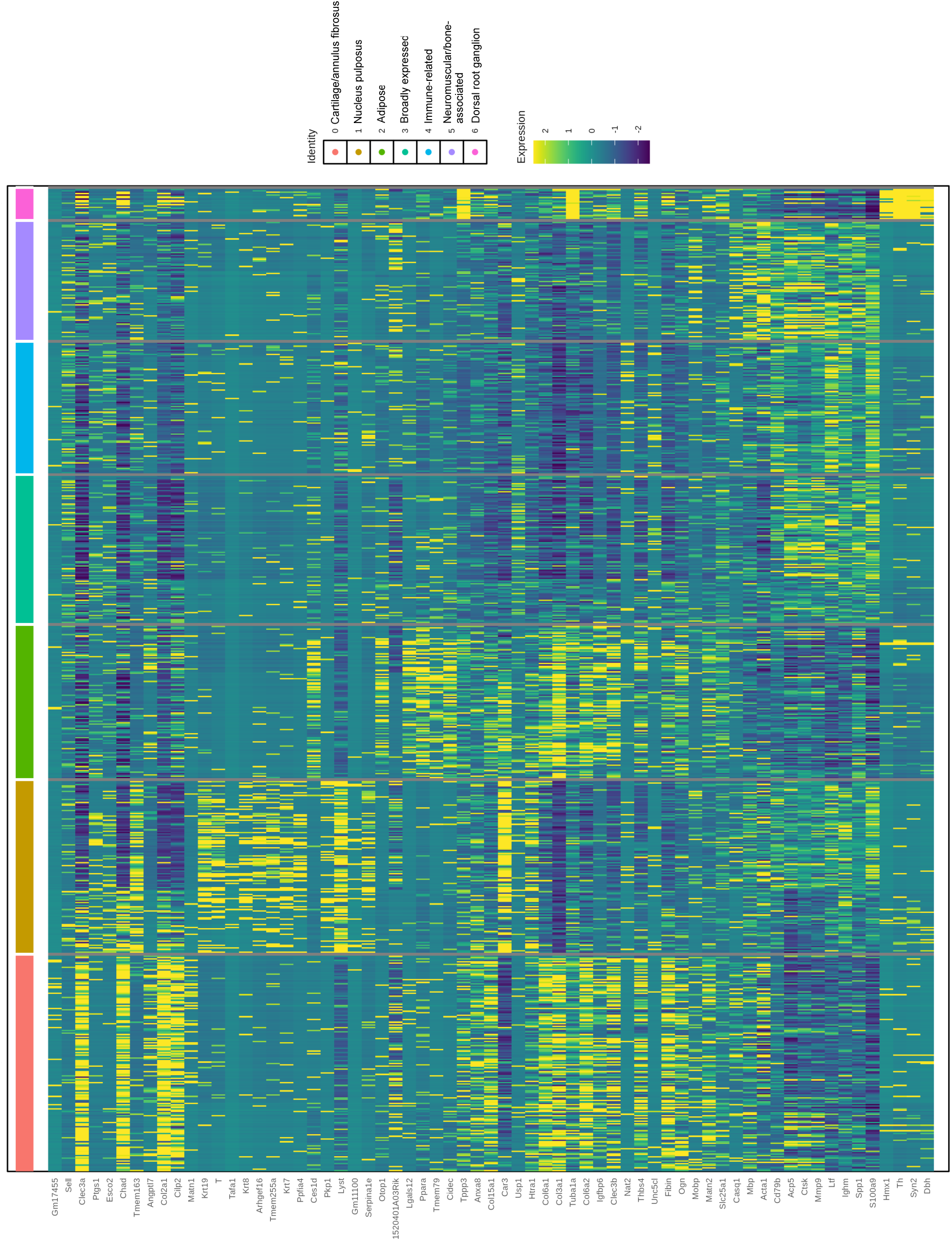
IVD specific gene cluster analysis and heatmap. Rows indicate the unique set of the top 15 expressed genes per cluster. Columns show cells grouped by clusters (color bar above). Values depict scaled normalized expression.

**Supplemental Figure 3.**
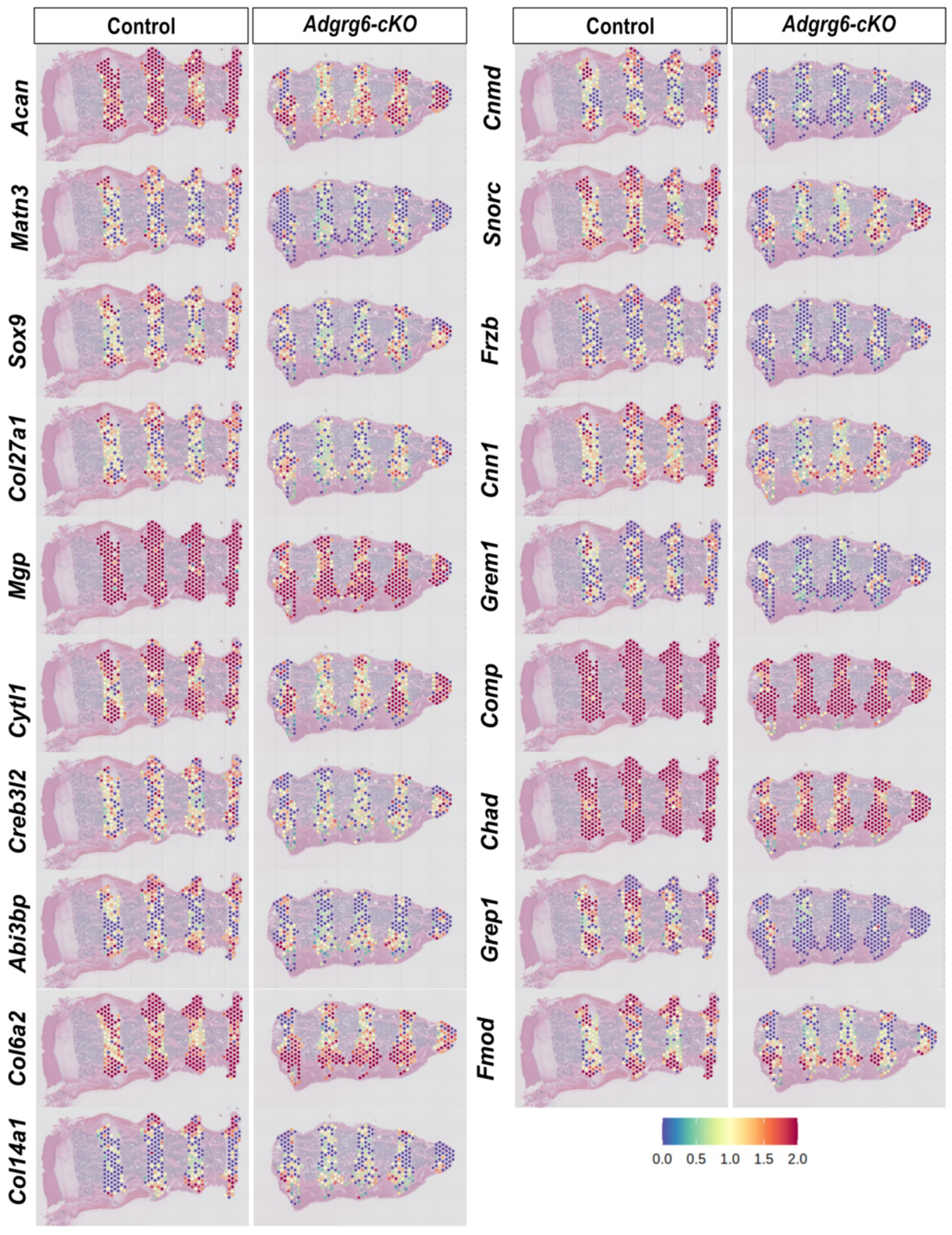
Intervertebral disc specific spatial gene expression altered in *Adgrg6-cKO* mice. Spatial feature plots show log-normalized cartilage development and extracellular matrix organization transcript abundance per spot. Color intensity corresponds to expression magnitude, with consistent scale across genes and samples.

**Supplemental Figure 4.**
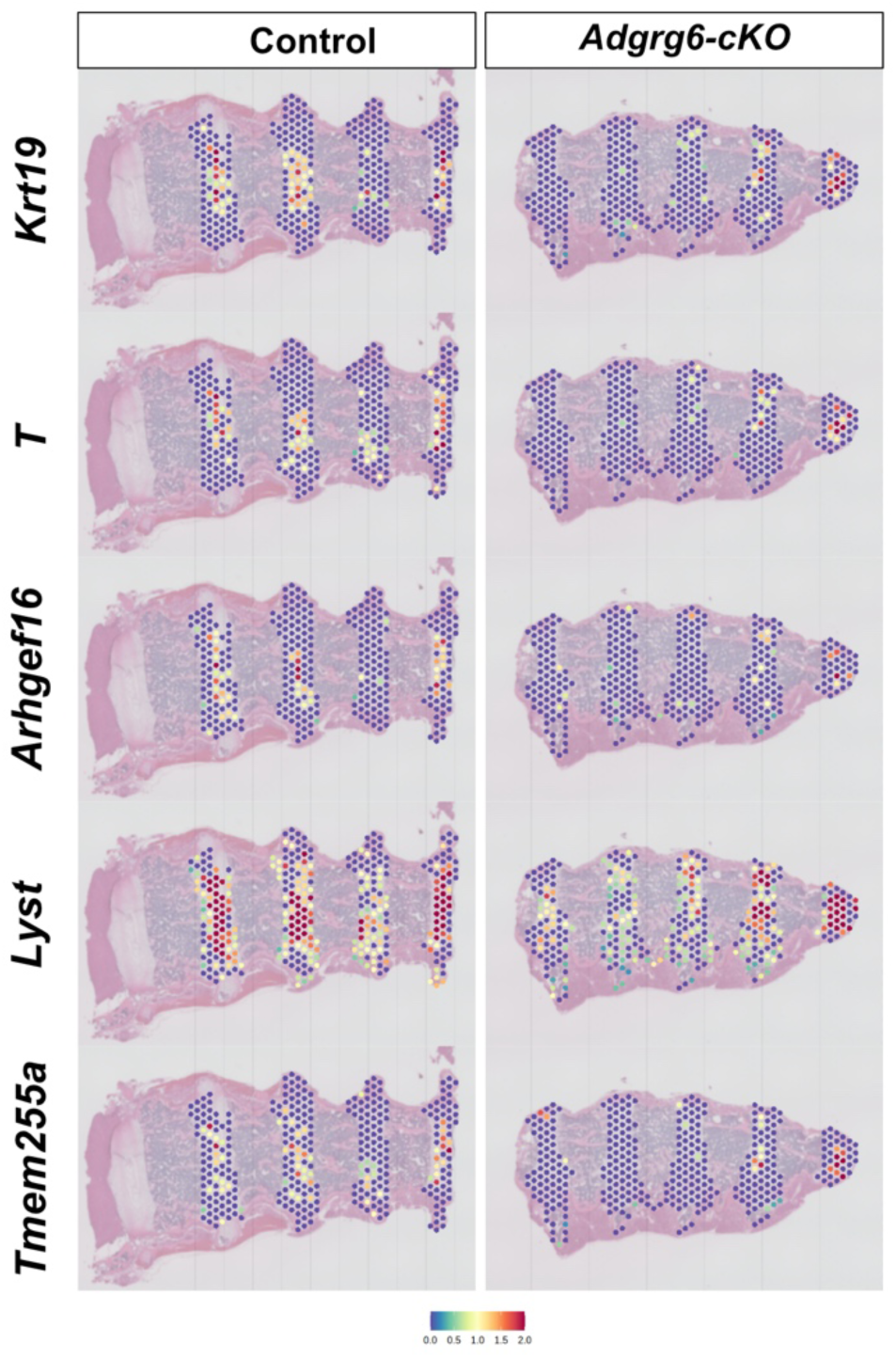
Nucleus pulposus specific spatial gene expression in wildtype and *Adgrg6-cKO* mice. Spatial feature plots show log-normalized nucleus pulposus transcript abundance per spot. Color intensity corresponds to expression magnitude, with consistent scale across genes and samples.

**Supplemental Figure 5.**
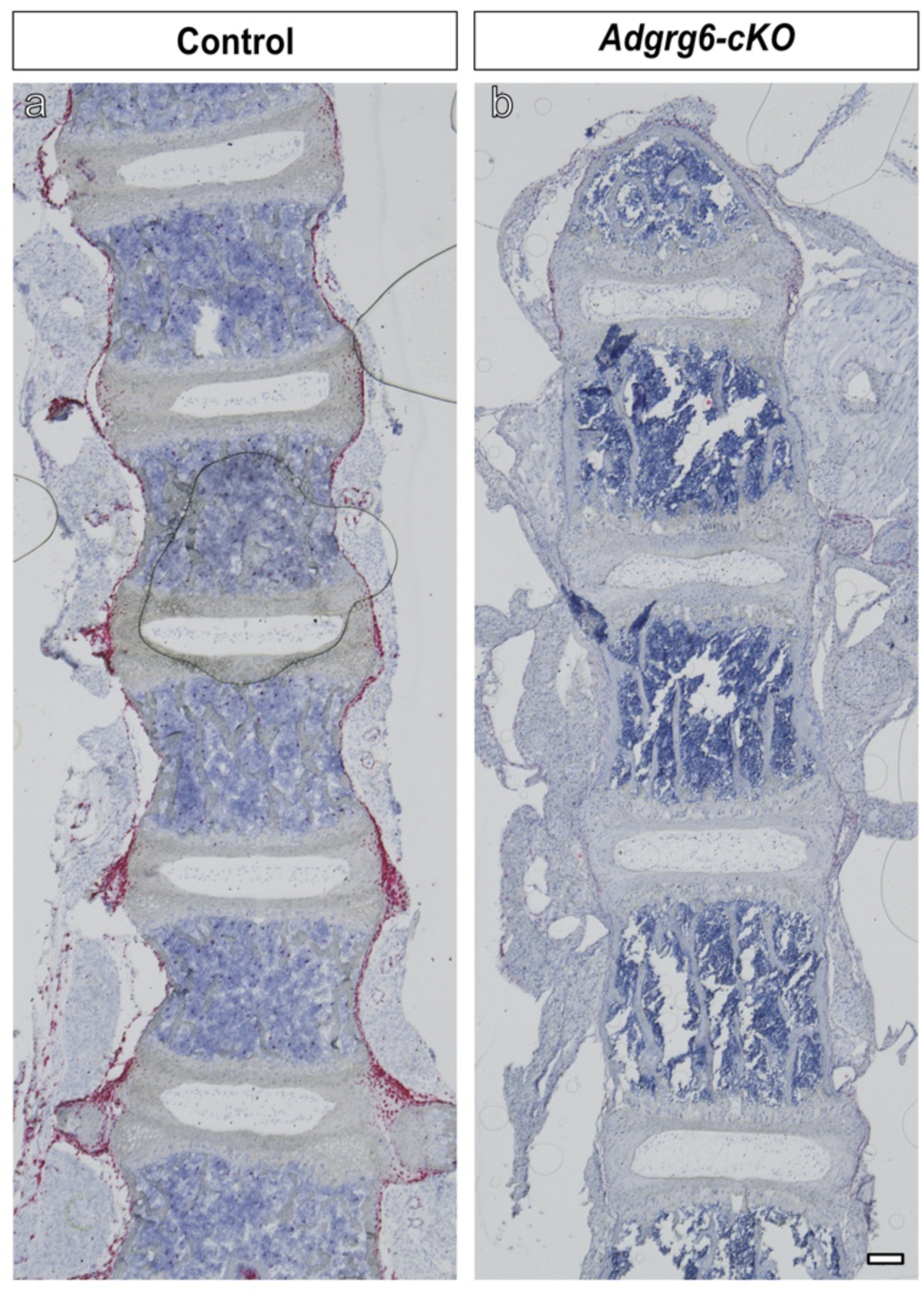
C*o*l14a1 is expressed in periosteum-like tissue adjacent to the spine. RNAScope *in situ* hybridization was performed to detect *Col14a1* expression in sagittal sections of the thoracic spine. In control mice, robust *Col14a1* expression is observed in periosteum-like tissue surrounding the vertebral column (a). In *Adgrg6-cKO* spines, *Col14a1* expression is markedly reduced and largely absent within these paraspinal tissues (b). Scale bar = 100 µm.

**Supplemental Figure 6.**
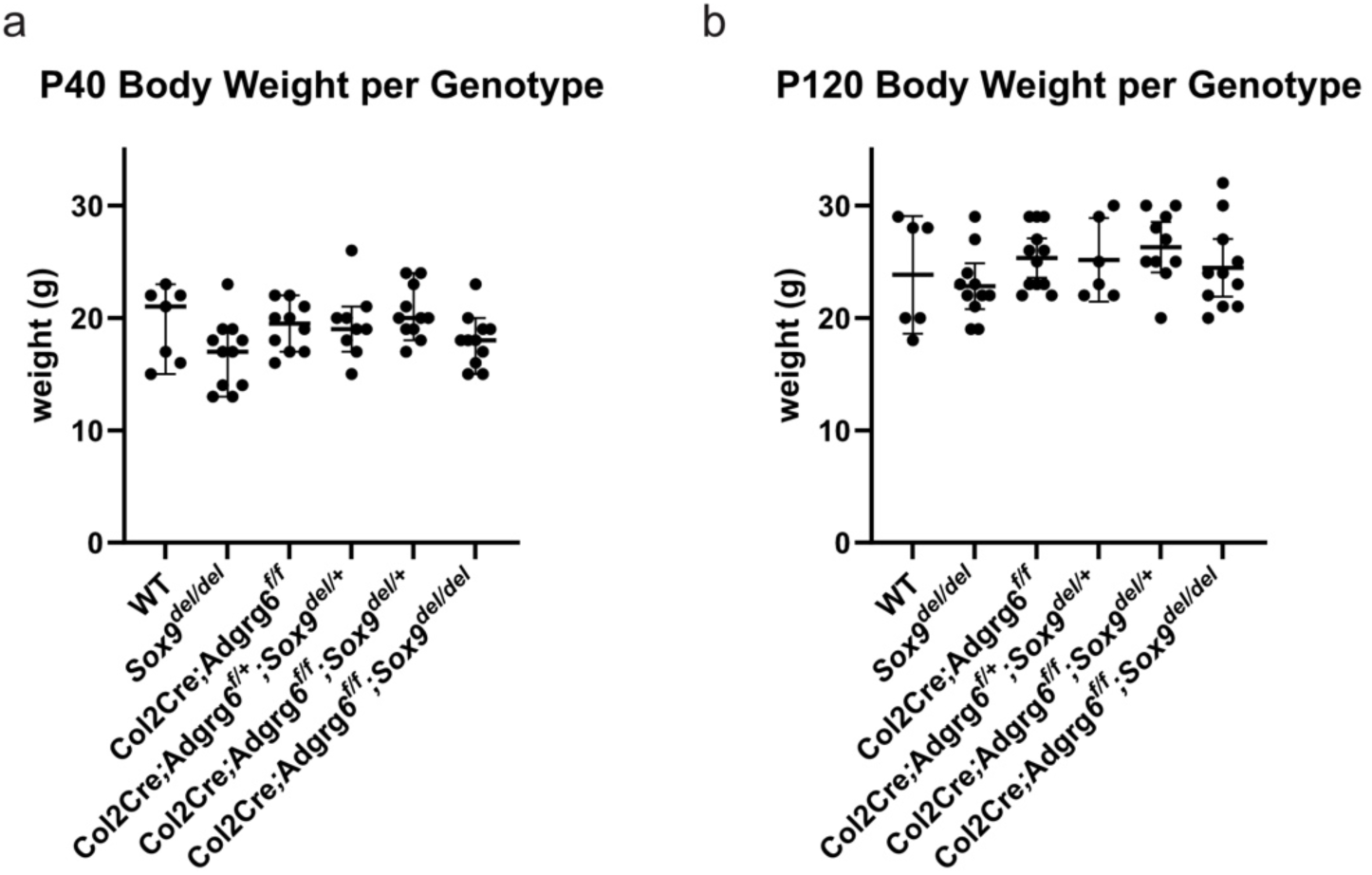
Longitudinal body weight measurements indicate normal growth patterns in experimental and control mice. Body weight measures of mutant and control mice scanned at P40 (a) and P120 (b) are not significantly different. Individual points are shown, with median ± 95% confidence intervals overlaid.

**Supplemental Figure 7.**
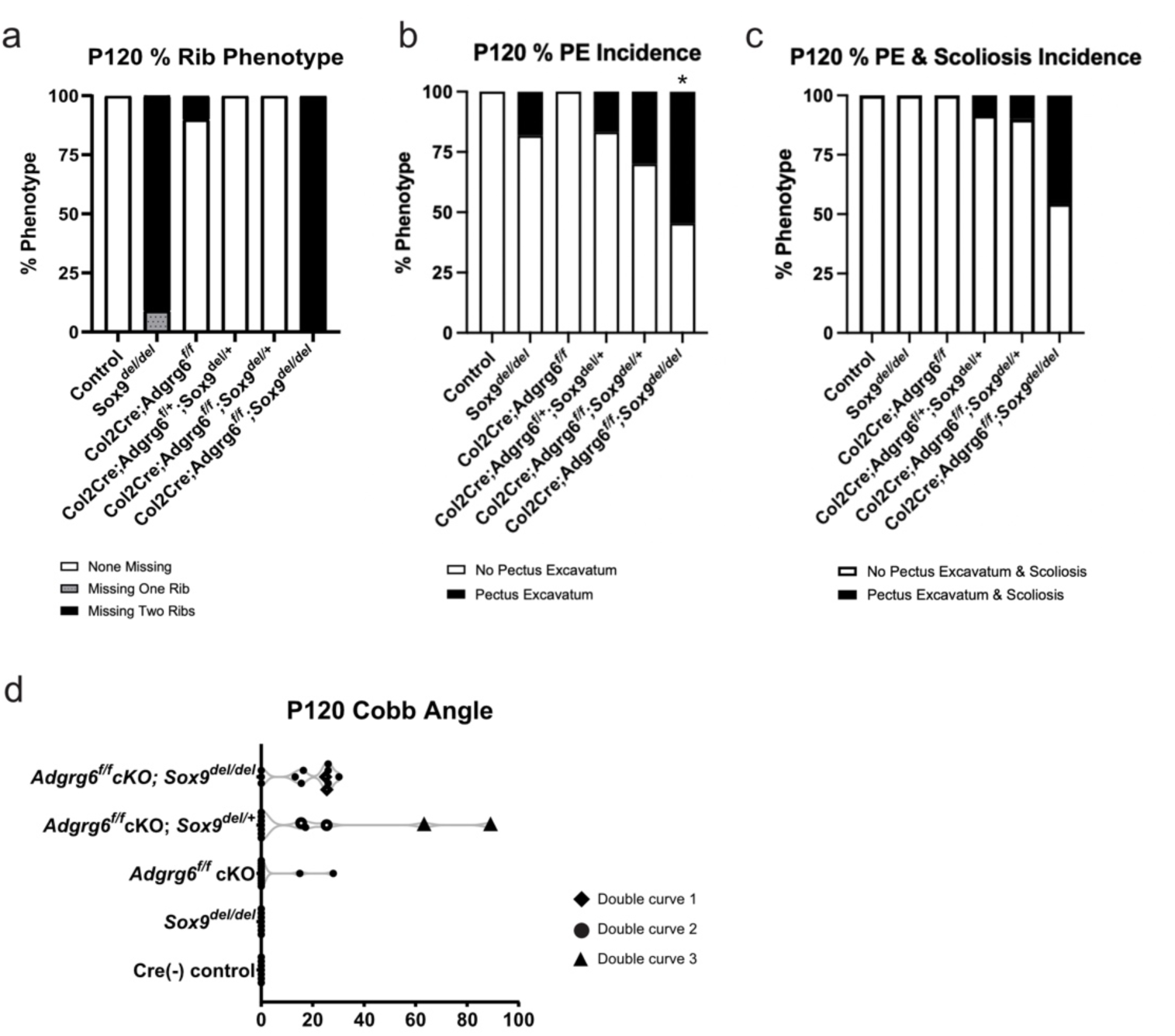
Genotype-dependent incidence of scoliosis, T13 rib loss, and pectus excavatum phenotypes at P120. Histogram of the percentage of mice exhibiting loss of the T13 rib pair phenotype at P120 across genotypes (a). Histogram of the incidence of pectus excavatum (PE) across genotypes (b). Fisher’s Exact Test for pairwise comparisons of PE incidence count at P120 showed significant genotype-dependent differences in *Adgrg6-cKO*; *Sox9^del/del^* double mutant mice when compared across genotype, * = p ≤ 0.05 (b). Histogram of the concurrent of PE and scoliosis at P120 (c), statistical analysis using Fisher’s Exact Test for pairwise comparisons showed no significant genotype-dependent differences for this relationship. Scoliosis severity distributions differed significantly by genotype, with *Adgrg6-*cKO; *Sox9^del/del^*double mutant mice significantly different from *Sox9^del^*, Col2Cre; *Adgrg6^f/+^*; *Sox9^del/+^*, and control mice (p ≤ 0.01)(d). Increased incidence and severity of thoracic scoliosis in *Adgrg6-*cKO; *Sox9^del/+^* and *Adgrg6-*cKO; *Sox9^del/del^* mutant mice quantification shows higher incidence of thoracic scoliosis and increased severity compared with controls (d).

**Supplemental Figure 8.**
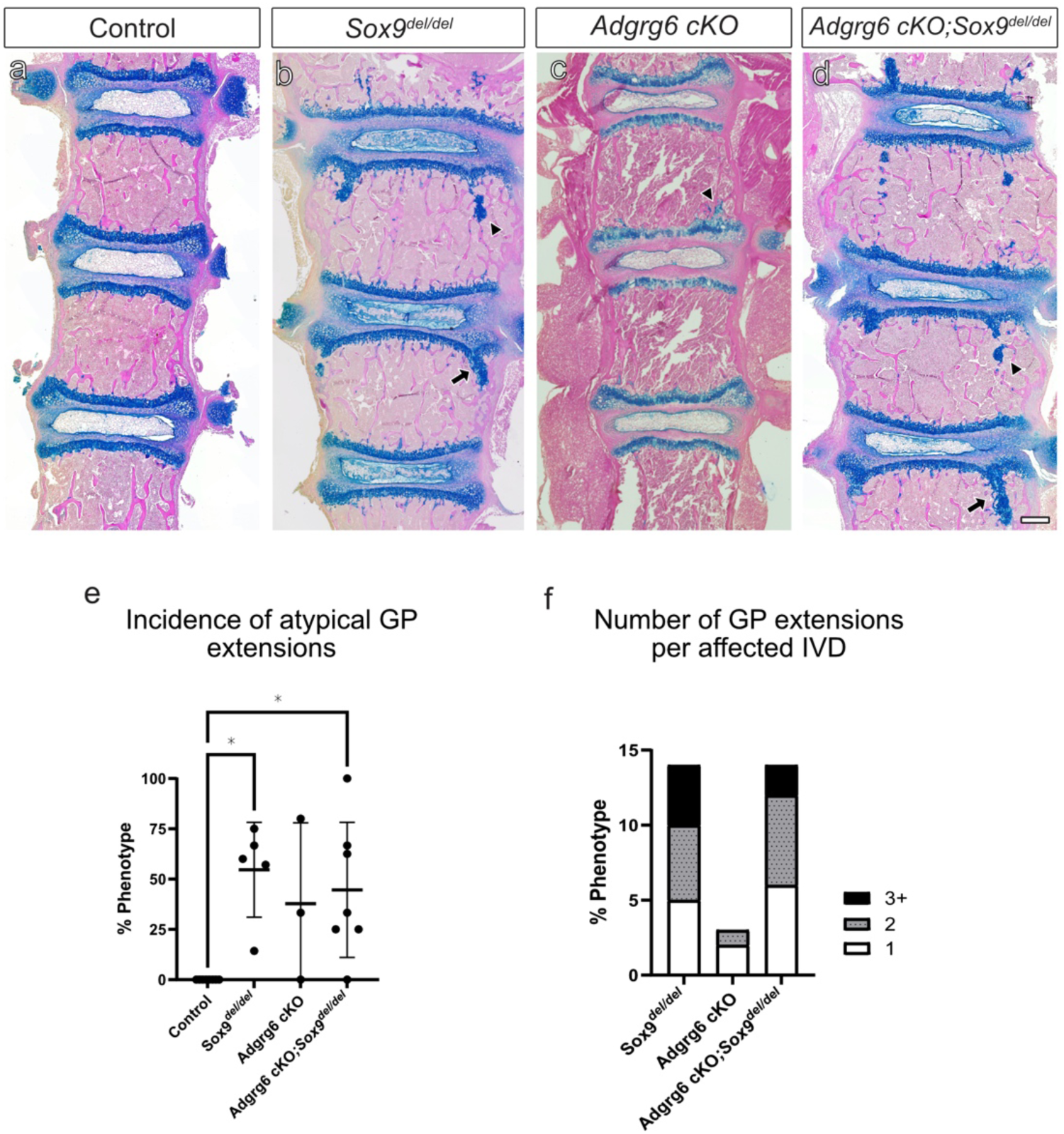
Histological analysis at (P)20 reveals growth plate defects and ectopic cartilage regions in vertebral bone of *Adgrg6-*cKO; *Sox9^del/del^* double mutant mice. Alcian Blue Hematoxylin Eosin Orange G stain reveals growth plate (GP) extensions (arrows) and ectopic cartilaginous regions in the vertebrae that are disconnected from the GP (arrowheads) in *Sox9^del^*, *Adgrg6-*cKO, and *Adgrg6-*cKO; *Sox9^del/del^* double mutant mice (b-d), that are not observed in wild-type mice (a). The incidence of these growth plate defects was quantified as the percentage of affected IVDs per spine, with each point representing an individual mouse with the median ± 95% confidence intervals overlaid (e). One way ANOVA identified genotype as a major source of variation (38% of total variance), with *Sox9^del^* and *Adgrg6-*cKO; *Sox9^del/del^* exhibiting significantly increased incidence compared to controls (p ≤ 0.05)(e). *Sox9^del^* and *Adgrg6-*cKO; *Sox9^del/del^* double mutant mice exhibited increased severity of GP defects, exhibit higher incidence of three or more defects per affected IVD (f). Scale bar = 500 um.

**Supplemental Figure 9.**
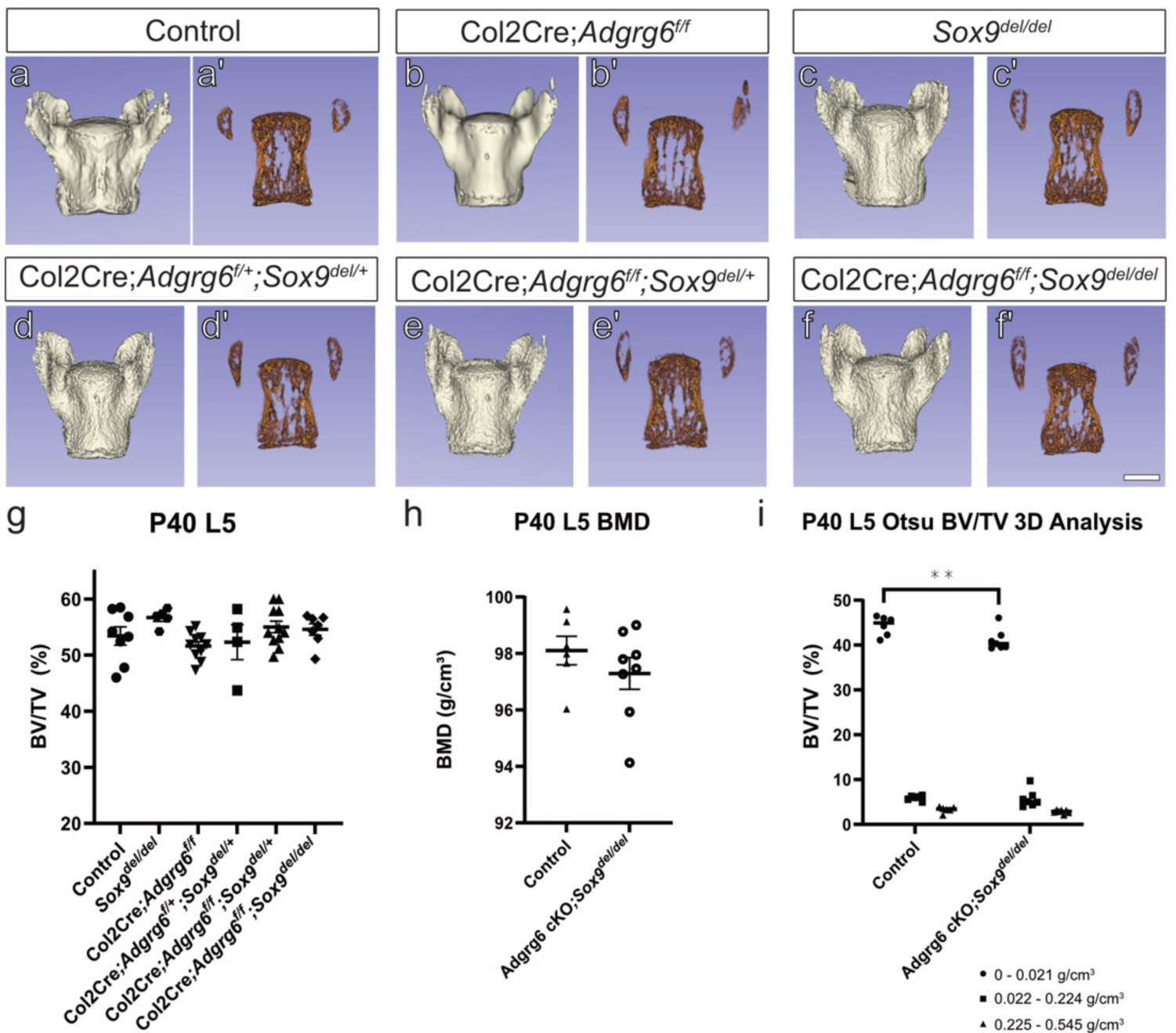
Lumbar (L)5 vertebral microarchitecture is largely preserved across genotypes at P40. MicroCT reconstructions of lumbar vertebrae (L)5 reveal no gross morphological differences across genotypes (a-f). Representative 10 µm midline slices similarly show no gross alterations of inner bone among genotypes (a’-f’). One way ANOVA and t-test analysis of bone microarchitecture revealed no significant changes in total bone volume (g) or bone mineral density (h). Two way ANOVA of OTSU-based threshold 3D bone volume analysis stratifying bone into low- (0–0.021 g/cm³), mid- (0.022 – 0.224 g/cm³), and high-density (0.225–0.545 g/cm³) fractions reveals a significant loss of low-density distribution in double mutant mice (*p* < 0.05) (i). Each dot represents an individual mouse, with mean ± SEM overlaid. Scale bar = 1 mm.

## METHODS AND MATERIALS

### Mouse strains

All animal studies were approved by the Institutional Animal Care and Use Committee at the University of Texas at Austin (AUP-2024-00222) and in accordance with national animal welfare guidelines. All mouse strains were previously described, including Col2a1-Cre ^48^, *Adgrg6^f/f^* (Taconic #TF0269), *Sox9^del^* ^33^. All mouse positioned in the scanner with the incisors secured in the nose cone and maintained on isoflurane at low flow during acquisition. Respiratory rate was continuously monitored throughout scanning using a tape marker on the dorsal thoracic region. *In vivo* microCT scans were acquired at 55 kV, 200 μA, 10 μm voxel size, 4K resolution, continuous mode, with 1 mm aluminum filter on Skycan 1276 (Bruker). Reconstructions were performed using the following parameters: smoothing = 5, ring artifact correction = 6, beam hardening = 10, and threshold = 0.065. Images were visualized and analyzed using CTVox, Dataviewer, CTAn Bruker software and 3D Slicer software (Version 5.6.1) including SlicerMorph ^49^. Cobb angles were measured in thoracic spine segments using 3D slicer. Curvature was quantified by measuring the angle from the apical vertebra to the most inferior tilted vertebra. P40 and P120 thoracic cavity morphometrics were visualized using 3D Slicer segmentation to colorize the thoracic column, including the ribs from the costal cartilage and sternum. Morphometric analysis of P40 T4 and L5 vertebrae were segmented in CTAn and sequentially straightened in DataViewer. Straightened datasets were re-opened in CtAn, and the vertebral midline was defined by identifying the most dorsal and most ventral slices containing the vertebral body. From the midline, 10 um slices were used to extracted to generate inner bone visualizations in 3D Slicer. All vertebral microCT datasets were processed using CtAn with standardized batch tasklists. Separate tasklists were used for whole vertebral body segmentation, BMD histogram analyses, and Otsu multi threshold-based analysis. Thresholds and ROI handling steps were held constant across all samples within each analysis type. Whole vertebral body processing was performed using a two-step thresholding approach. Tissue was first segmented using a grayscale threshold of 55–255. A bitwise operation was then performed to define the ROI (ROI = Copy Image), followed by image reload to apply the ROI mask. Bone was subsequently segmented using a threshold of 80–255, and standard 3D analysis was performed. BMD measures were obtained from histogram analysis in three-dimensional space within the defined volume of interest (VOI). Otsu-based thresholding analysis was performed as previously described ^43^. Spine and individual vertebral segmentations were subject to colorimetric threshold mapping to visualize the Otsu-based thresholding.

### Histological Analysis of Mice

For staining and immunohistochemistry protocols, P20 thoracic spines were histologically processed as previously described by Liu et al ^21^. Alcian Blue Hematoxylin Eosin Orange G staining was completed using standard protocols from Center of Musculoskeletal Research, University of Rochester. Immunohistochemistry analysis against UCP1 (1:200) (Thermofisher, #PA1-24894) was performed using a DAB chromogenic kit (Vector SK-4105). Slides were baked at 60°C, deparaffinized in xylene, and rehydrated through graded ethanol to water. Antigen retrieval was performed using proteinase K (Invitrogen #25530-015) for UCP1 and pepsin (Sigma P-7000) in HCl for SNORC. Endogenous peroxidase activity was quenched, and sections were blocked with normal serum prior to overnight incubation with primary antibodies at 4°C. The following day, the signal was developed using a DAB chromogenic detection kit. Slides were rinsed, dehydrated, and cover slipped for brightfield imaging.

### RNAScope Staining for Spine Sections

RNA *in situ* probes including Mm-Acan (439101), Mm-Apobec2 (482001), Mm-Chad (484881), Mm-Col14a1 (581941), Mm-Col27a1 (520001), Mm-Col6a2-C1 (1084001-C1), Mm-Comp (480711), Mm-Creb3l2-C1 (1079301-C1), Mm-Fmod (479421), Mm-Frzb (404861), Mm-Grep1-C1 (1755071-C1), Mm-Myh2 (401401), Mm-Ryr1-C1 (1120301-C1), and Mm-Sox9 (401051) were purchased from Advanced Cell Diagnostics (ACD). Thoracic spines were histologically processed following a protocol specific for skeletal tissue for RNA analysis^50^. P20 spine tissues were fixed in 4% paraformaldehyde (PFA) at room temperature for 24 hours, washed 3X 15 minutes in RNAase-free 1X PBS, decalcified in a Morse’s Solution for 24 hours at room temperature, and washed again 3X 15 minutes in RNAase-free 1X PBS. Samples were immediately processed in a 70%, 80%, 90%, 95%, 100%, and 100% ethanol series followed by two changes of xylene, and two changes of paraffin. Samples were paraffin-embedded and sectioned at a 4 μm thickness. RNA *in situ* analysis was performed using RNAScope 2.5 HD Detection Reagent – RED kit (ACD Bio) following the manufacturer’s instructions, with modifications optimized for skeletal tissues as previously described ^50^.

### Spatial Transcriptomics Cohort, Data, and Quality Control

Mouse thoracic spine sections from an Adgrg6 mutant and a wildtype control (n=1 per condition) were profiled using 10x Visium v1 spatial transcriptomics with our previously established workflow ^46^. To ensure disc centric signal and minimize off target high UMI contamination from adjacent tissues (e.g., muscle), intervertebral disc (IVD) regions were manually delineated in 10x Genomics Loupe Browser v8.0.0 Loupe Browser, and downstream analyses were restricted to these ROIs. Genomics Space Ranger (v3.0.0) was used to compute raw data, and the outputs were imported into Seurat (v.4.2.0) in R (v.4.2.1); spot level QC metrics (feature counts, UMI counts, mitochondrial percentage) were computed, and high quality IVD spots were retained using 200<nFeature_Spatial<8000 and %mt<5%.

### Mitigating Sampling Bias: Muscle-filtering Strategies in Spatial Clustering

Differential sampling of muscle adjacent to the IVD can inflate global signals and confound disc-intrinsic DE. We therefore applied a gene-based exclusion bias-mitigation strategy using Muscle-score filtering. A per-spot “muscle-ness” score was computed from canonical muscle genes, and spots with Muscle percentage >1% were excluded. The cleaned dataset was re-normalized and re-clustered to recalibrate variance. Joint analysis was performed on the cleaned dataset using SCTransform residuals. Principal component analysis (PCA) was computed, and the effective dimensionality was set to PCs≈9 based on elbow shape and cumulative variance criteria. Downstream analysis utilized Seurat (v.4.2.0) build-in functions. UMAP embeddings were generated on PC 1:9, followed by k-NN graph construction and Louvain clustering across a resolution grid 0 to 1. Cluster stability was inspected with clustree, and a final working resolution of r=0.5 was selected for interpretability and modularity. Spatial context was obtained by overlaying cluster assignments on H&E images with SpatialDimPlot. Cluster annotation used FindAllMarkers (Wilcoxon; thresholds log 2 FC>0.25, p adj<0.05) and curated IVD marker sets to assign biological identities. For DE, count layers were unified and analyzed on the Spatial assay with log normalization to yield interpretable log2 FC and p-values (Wilcoxon test; thresholds log 2 FC>0.25, p adj<0.05).

### Cluster Marker Identification

After muscle-score filtering and re-clustering, cluster identities were set. We conducted One-vs-all marker discovery used Seurat’s FindAllMarkers on the SCT assay. To emphasize cluster specificity, stringent markers were additionally defined as detected in more than 50% of spots in the target cluster and detected in fewer than 10% of spots outside that cluster. The top 20 up-regulated markers per cluster were ranked by fold change. Visualization included per-cluster volcano plots (one vs rest), and a heatmap of the top 15 significant markers per cluster.

### Assessing Disc Programs: Key Genes and Pathway Analysis

To assess the disc program switch, spatial and UMAP feature plots were generated for Sox9 and key ECM/cartilage genes using log-normalized data from the Spatial assay to visualize regional expression differences between conditions. Gene Ontology over-representation analysis (GO-ORA; Biological Process) was performed with clusterProfiler on “Down in mutant” gene sets, using Benjamini–Hochberg adjustment; focus terms included cartilage development (GO:0051216) and extracellular matrix organization (GO:0030198). Gene Set Enrichment Analysis (GSEA) was conducted with clusterProfiler’s GSEA function using a ranked list created from the muscle-filtered *Adgrg6-cKO* vs Control comparison, ranking all genes by average log fold change (ranging from the most upregulated to the most downregulated genes). Cluster-specific marker gene sets were defined via FindAllMarkers on the re-clustered dataset, filtered at avg_logFC > 0.25 and p_adj<0.05. Pathway-level enrichment for “cartilage development” and “extracellular matrix organization was visualized showing the enrichment score “mountain” curve and the ranked-list “barcode.”

### Fluorescence detection

Lumbar spine/IVD from P7 *Sox9^EGFP^* mouse was dissected and fixed in 4% paraformaldehyde (in PBS) at 4 °C overnight, washed with PBS, and incubated in 30% sucrose at 4 °C overnight. Tissues were embedded in OCT, and frozen sections were cut at 16 μm using a cryostat. Slides were washed in PBST (Phosphate Buffered Saline with 0.1% Tween 20) 3 X 15 minutes at room temperature; and mounted under a coverslip with Aqua-mount (13800; Lerner Labs). EGFP fluorescence in intact spine/IVD was directly imaged, using a dissection Leica fluorescent microscope. EGFP fluorescence in cryosections was directly imaged at the Core for Imaging Technology & Education at Harvard Medical School.

### Fluorescence-activated cell sorting from the spine/IVD of postnatal day 7 *Sox9^EGFP^* mouse

To determine whether Sox9 directly regulates the expression of *Adgrg6* by binding to its regulatory elements in the spine/IVD, we employed FACS to isolate Sox9-expressing cells (EGFP+) and Sox9 non-expressing cells (EGFP-) from P7 *Sox9^EGFP^* mice. The spine/IVD was dissected, and the tissues surrounding the vertebral body and intervertebral disc were removed under a fluorescent dissecting microscope. The dissected spine/IVD tissues were further digested in 2.5 % collagenase type II (Worthington, Cat#: LS004177) plus 0.5% collagenase type I (Worthington, Cat#: LS004196) at 37 ^O^C in a thermomixer (Eppendorf ThermoMixer C, 700 rpm) for 2.5 hours to 3 hours. The homogenate was filtered with a 35 μm cell strainer (Falcon, Cat#: 352235). The cells were pelleted by centrifugation for 5 min at 300 g, washed with PBS at least four times, and resuspended in FACS sorting buffer (PBS plus 1% fetal bovine serum; DRAQ7 dye or DAPI was added to the solution at 1/1000 dilution). DRAQ7 dye (Novus Biologicals Cat# NBP2-81126) is a DNA dye that fluoresces at far-red upon DNA binding, and DAPI (4’,6-diamidino-2-phenylindole) is a DNA dye that fluoresces at blue; both are only capable of penetrating dead cells with compromised membrane structure. The cells were passed through the cell strainer (Falcon, Cat#: 352235) immediately before putting them in the FACS. The digested single cells were sorted using a SONY SH800 cell sorter to separate different populations that express or don’t express EGFP. For cell sorting, all particles (events) passing over the laser were first evaluated for forward scatter (FSC) and back scatter (BSC) to eliminate aggregates and debris by gating for those cells that were in the center of the FSC/BSC graph. The doublets were also eliminated by gating for those cells in the linear range of forward scatter height (FSC-H) and forward scatter area (FSC-A). Subsequently, the dead cells were eliminated by gating for those that were negative for DRAQ7 DNA dye or DAPI. After the gates were set, the laser power, gain, and all other parameters of the cell sorter were kept the same throughout the experiment. Although the same laser parameters were used across experiments, the process of gating was assessed for every litter that was sorted to ensure the accuracy of the gates.

### ATAC-Seq and Cut&Run-Seq

After FACS, cells were washed twice in ice-cold PBS. 50,000 cells (per sample) were used to perform ATAC-Seq, and 500,000 cells (per sample) were used to perform Cut&Run-seq, following the detailed protocols described in our most recent work ^51^.

### Statistics

Statistical analysis and graphs were generated using R (v.4.2.1) and GraphPad Prism (version 10.2.1). Statistical significance was determined by a p-value of less than 0.05.

### Statistics Code Availability

All code used to generate spatial transcriptomics results has been deposited on Github (https://github.com/xuziyi0909/Mouse_Spine_Intervertebral_Disc_SpatialTranscriptomics).

## Acknowledgements

We thank Sylvie Beaudenon-Huibregtse, Jessica Podner, and Anna Battenhouse and the team at the Genomic Sequencing and Analysis Facility at the University of Texas at Austin, Center for Biomedical Research Support for helping with the Spatial Transcriptomics experiment (RRID#: SCR_021713). This work was supported by grants from NIH to A.B.L. (NIAMS: R01AR074385 and R01AR076562), to C.-H.Z. (NIAMS: R21AR081990), to R.S.G. (NIAMS: R01AR072009), and to Z.L. (NIAMS: R00AR077090 and R01AR083966).

## Author Contributions

VA designed, performed, and analyzed most of the experiments.

ZX analyzed spatial transcriptomic data.

CHZ and JK performed ATAC-seq and CUT&RunSeq

JI performed microCT segmentation and analysis under the supervision of VA.

KM performed histological staining, RNA *in situ* hybridization, and slide imaging under the supervision of VA.

AL provide guidance and supervision of CZ and JK

ZL performed spatial transcriptomics

RG provided guidance with the project and supervised VA, ZX, JI, and KM.

RG and VA designed most experiments, interpreted results, and alongside ZX wrote the manuscript.

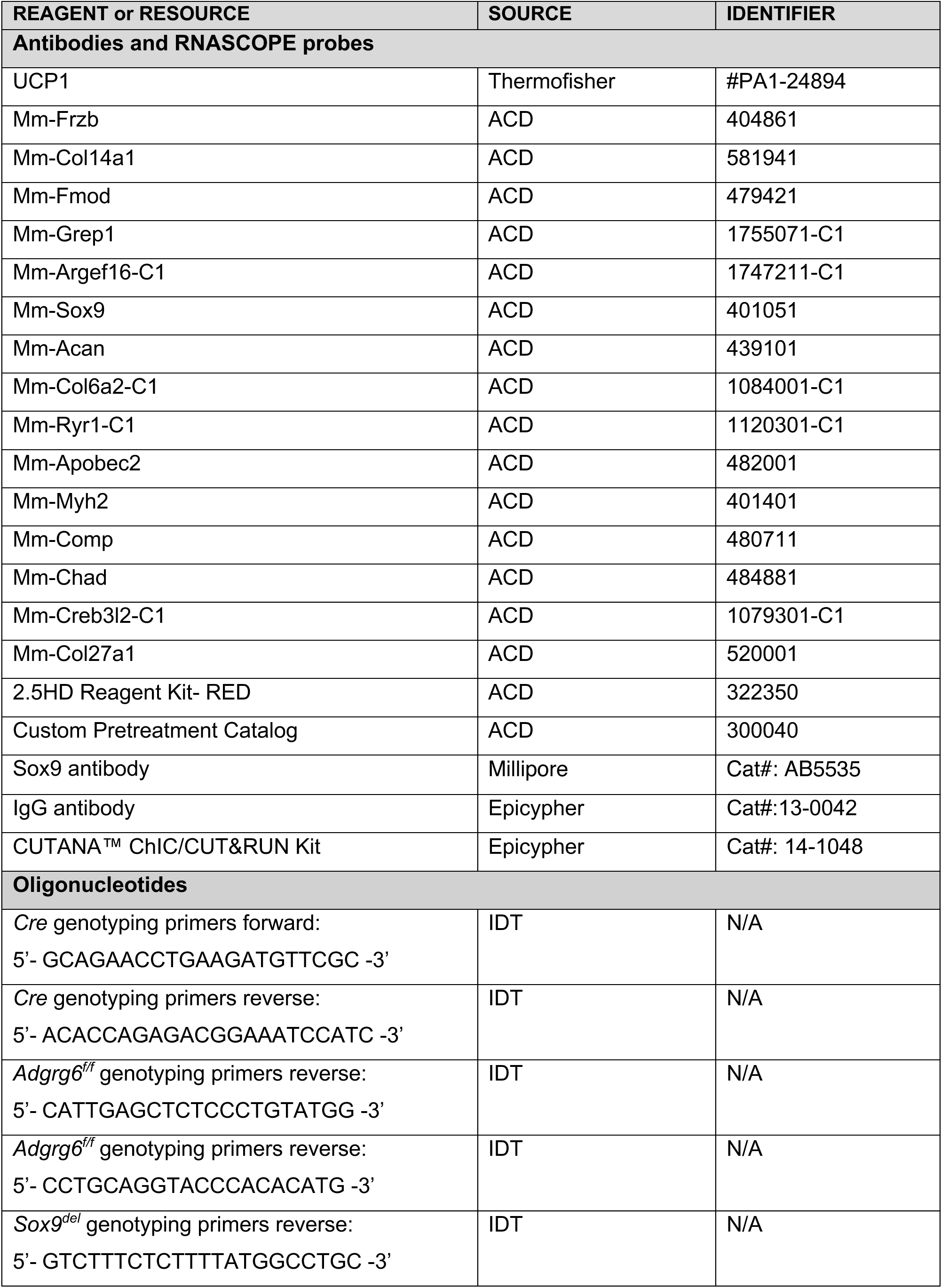
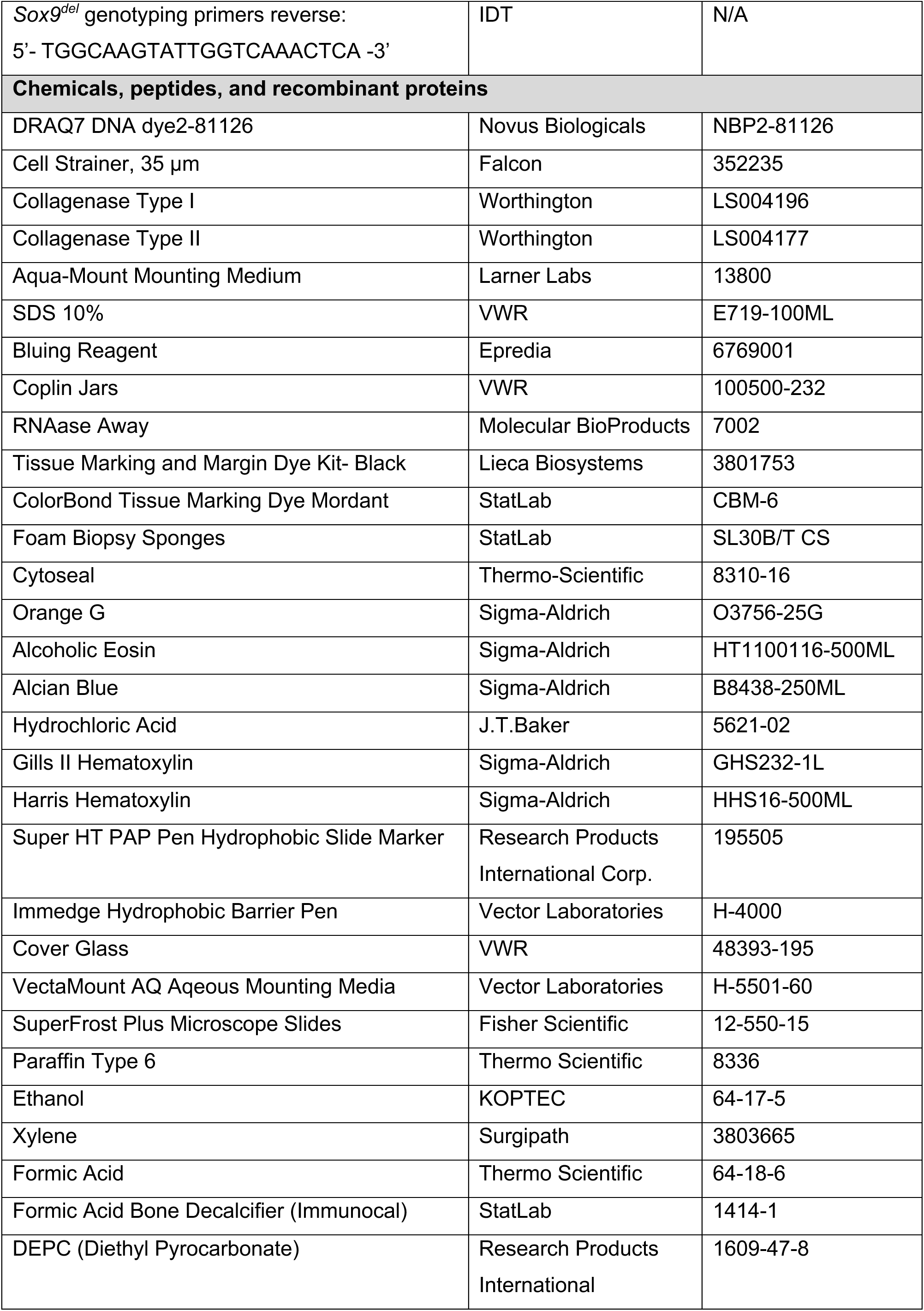
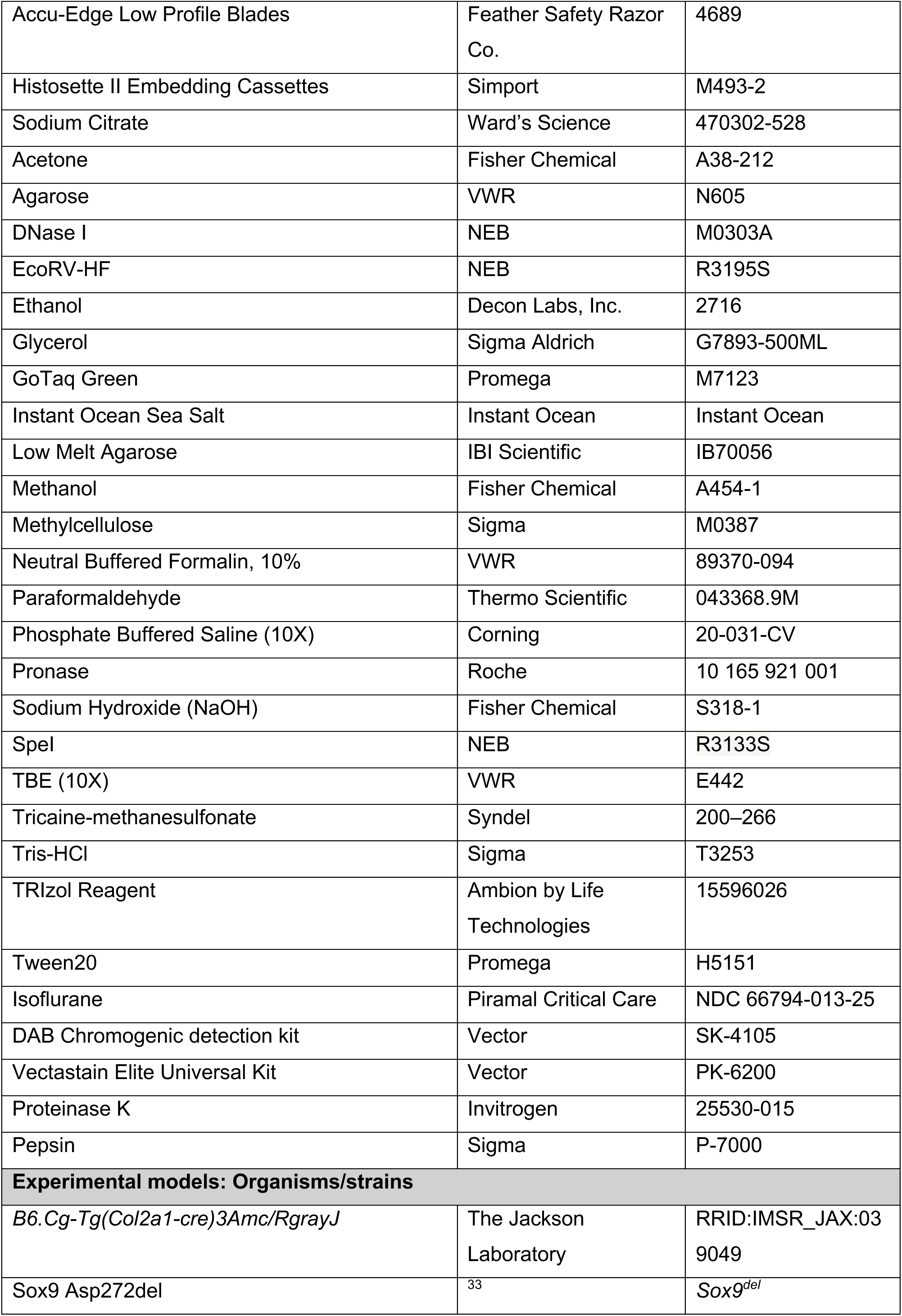
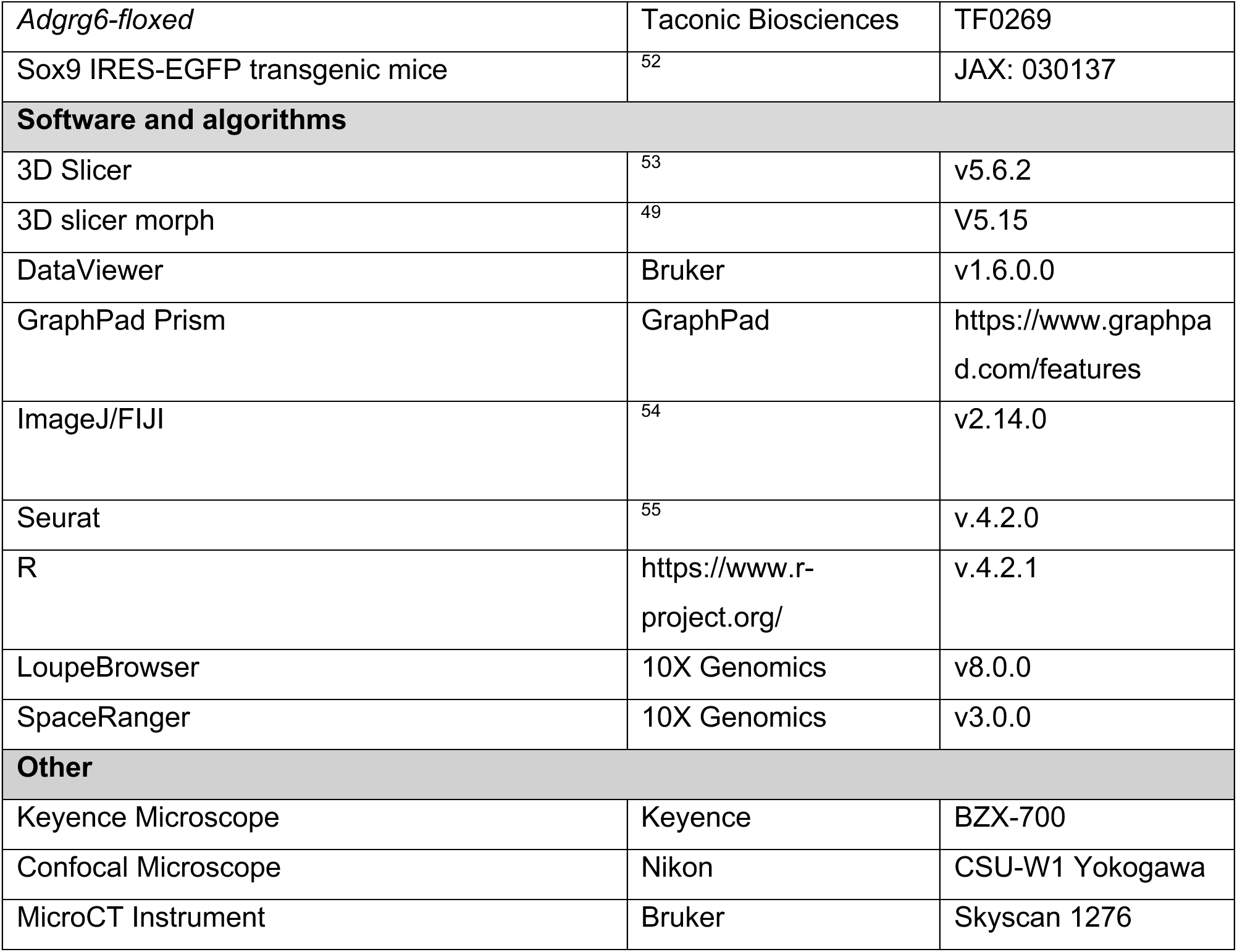

## Supporting Documents

